# Colon-delivered multivitamin supplementation enhances working memory-related fMRI responses in older adults: a randomized, placebo-controlled trial

**DOI:** 10.64898/2026.03.10.709744

**Authors:** Lianne B. Remie, Mark R. van Loenen, Mechteld M. Grootte Bromhaar, Noortje M.P. Overwater, Janne van Overbeek, Andrea Anesi, Urska Vrhovsek, Ateequr Rehman, Robert E. Steinert, Jurriaan J. Mes, Guido J.E.J. Hooiveld, Wilma T. Steegenga, Joukje M. Oosterman, Mara P.H. van Trijp, Esther Aarts

## Abstract

**Background:** Vitamins are important modulators of intestinal health and may affect the gut-brain axis through microbial metabolites such as short-chain fatty acids (SCFAs). However, the neurocognitive effects of colon-delivered vitamins in older adults remain unexplored – a critical gap given the gut-brain axis’s emerging role in cognitive aging.

**Objective:** We investigated the effect of a colon-delivered multivitamin (CDMV) supplement on intestinal health and neurocognitive outcomes in older adults at risk of cognitive decline.

**Methods:** Within the double-blind randomized placebo-controlled trial COMBI (ClinicalTrials.govID: NCT05675007), we included 75 older adults (60-75 years) at risk of cognitive decline based on lifestyle-related factors. Participants consumed a colon-delivered capsule with vitamins B2, B3, B6, B9, C and D3, or a placebo, daily for six weeks. Pre- and post-intervention, we employed neuroimaging, feces- and blood collection. Primary outcomes were fecal SCFA concentrations, working memory (WM)-related fMRI responses, and WM performance measured with the n-back task.

**Results:** After adjusting for baseline values, we found no significant between-group differences in total fecal SCFA levels (p=0.30) and WM performance (p=0.50). Post-intervention WM-related fMRI responses in the hippocampus (p=0.01; ηp²=0.09) and dorsolateral prefrontal cortex (dlPFC) (p=0.06; ηp²=0.04), driven by the right dlPFC (p=0.02), were higher in the CDMV group compared to placebo. Independent of intervention group, post-pre increases in fecal SCFA levels were significantly correlated to increases in dlPFC fMRI responses (ρ=0.31; p=0.02) and WM performance (ρ=0.43; p=0.001).

**Conclusions:** Our findings suggest that CDMV supplementation increases WM-related responses of the dlPFC and hippocampus in older adults, but this effect was not accompanied by changes in fecal SCFA levels or WM performance. The positive correlation of within-subject changes in fecal SCFAs with changes in WM dlPFC responses and performance across intervention groups provides human evidence for gut-brain communication in cognitive aging beyond cross-sectional associations.

## Introduction

The gut and its microbiome are increasingly recognized as key contributors to cognitive aging. Over the lifespan, our microbiome undergoes age-related changes (1). Compared to younger individuals, older adults exhibit an altered microbiota composition (2) and diversity (3), which both have been linked to cognitive performance. Moreover, gut microbiota of aged individuals with severe cognitive decline show substantial differences compared to healthy aged adults (4).

Given the complexity of the microbial ecosystem (5), metabolites are gaining more attention as functional microbiome indicators (6). Short-chain fatty acids (SCFAs), such as acetate, butyrate, and propionate, are abundant bacterial fermentation products known for their anti-inflammatory effects (7). Moreover, SCFAs are suggested to play a pivotal role in gut-brain communication (8). Studies in mice demonstrate that SCFAs can reduce neuroinflammation (9), maintain blood-brain barrier integrity (10), improve cognition (11), and might be key factors in age-associated cognitive decline (12). Importantly, cross-sectional studies show that human fecal SCFAs levels decrease with age (13, 14) and that these reduced SCFA levels become more pronounced in patients with severe cognitive decline compared to healthy aged individuals (4, 15, 16). However, causal evidence for involved human gut-brain pathways is scarce and neurocognitive mechanisms remain unresolved.

Overall, the gut microbiome and its metabolites represent promising intervention targets for cognitive aging (17, 18). Such interventions (e.g., administering pro- or prebiotics) could elevate fecal SCFAs (19) and might influence cognition in aging, although current evidence is mixed. While probiotics primarily benefit cognitively impaired individuals based on meta-analyses (20, 21), prebiotics show - despite limited research - greater potential in healthy older adults (22). Traditional prebiotics are predominantly carbohydrate-based (i.e., fermentable fibers) (23). Notably, novel approaches considering the complete microbial ecosystem tend to broaden and complement this spectrum (24, 25).

Vitamins are important shapers of the intestinal environment and essential microbial micronutrients. They act as co-factors in fermentation, can affect gut barrier integrity, have antioxidant capacities, and could hereby modulate the gut microbiome (26). Especially B-vitamins (27), but also vitamin C (28) and D3 (29), have been proposed as substances that lie adjacent to the current boundaries of prebiotics and may modulate the gut microbiome in a prebiotic-like effect (24). Although some gut bacteria can produce (B-)vitamins *de novo* and cross-feed other bacteria, the influx of micronutrients from the diet is essential (30). Yet, most dietary vitamins are absorbed in the small intestine, and only reach the colon when consumed regularly as present in slow-fermentable foods (31). For this reason, colon-targeted vitamin administration could be a promising, novel gut intervention (24, 32). *In vitro* digestion and fermentation showed that vitamin B2 in a colon-targeted encapsulation led to increased bacterial SCFA production, and validated the colonic delivery method (33). Moreover, a pilot study in healthy participants (*n*=96, 7 arms) demonstrated that a 4-week colon-delivered supplementation of vitamins B2, C, and D could modulate bacterial composition (B2, C, D), increase alpha diversity (C) and elevate fecal SCFA levels (C) versus placebo (34). Finally, high-dose oral vitamin B2 supplementation – reaching the colon through systemic overdosing – increased fecal butyrate levels as well in a two-week trial with healthy volunteers (35).

Notably, inadequate vitamin intake is common in older adults (36). Especially individuals with a suboptimal lifestyle (e.g., Western diet) might have low vitamin availability in the colon (37). Crucially, these same suboptimal lifestyle factors are important predictors of cognitive decline (38), positioning at-risk older adults as a relevant target group for preventive interventions. Yet, despite the relevance of this target group, gut-brain neurocognitive mechanisms in cognitive aging remain particularly understudied. Only a handful of studies have assessed probiotic effects on functional neuroimaging in younger volunteers (39), and just one recent study has examined this in older adults, focusing solely on resting-state functional connectivity (40). Therefore, we investigated the effect of a 6-week colon-delivered multivitamin supplementation (containing vitamin B2, B3, B6, B9, C, and D3) versus placebo on intestinal health (fecal SCFA levels) and neurocognition (working memory-related fMRI responses and performance) in at-risk older adults.

We selected fecal SCFA levels as primary intestinal outcome, based on their demonstrated sensitivity to colon-targeted vitamins (33–35). We selected working memory (WM) as our primary neurocognitive outcome given its age-related decline (41), role in cognitive variability (42), and demonstrated responsiveness to gut interventions across species and age groups (11, 22, 39, 43). Our fMRI regions-of-interest (ROIs) – the dorsolateral prefrontal cortex (dlPFC) and hippocampus – were chosen for their vulnerability to aging (44, 45), involvement in WM (46, 47), and prior associations with gut interventions or microbiome alterations (39, 48). In addition to our primary outcomes, we examined various secondary intestinal-, immune- and neurocognitive outcomes related to gut-brain communication pathways that may underlie both cognitive aging (18) and vitamin effects on neurocognition (49).

## Methods

### Study design

This study was a randomized, double-blind, placebo-controlled trial involving two research centers: the Donders Centre for Cognitive Neuroimaging, Radboud University, Nijmegen, The Netherlands (RU-DCCN) and the Division of Human Nutrition and Health, Wageningen University & Research, Wageningen, The Netherlands (WUR-HNH). At baseline (T0) and after the 6-week intervention period (T1), participants visited both research centers within one week (**Figure 1**). At the RU-DCCN, participants underwent neuroimaging and neuropsychological testing. In the week prior to their WUR-HNH visit, participants collected fecal samples and completed questionnaires. At the WUR-HNH, blood samples were drawn in a fasted state. To ensure no wash-out effects in blood measurements, both WUR-HNH visits were planned directly before and after the intervention period. After baseline visits were completed, participants were randomly assigned to one of two intervention groups: colon-delivered multivitamin (CDMV) or placebo. Subsequently, they started their 6-week intervention period. The study was conducted in compliance with the Declaration of Helsinki for research involving human participants, and the complete procedure was approved by the local Ethics Committee (METC Oost-Nederland, NL80063.091.22) and registered in the Clinical Trial Register (ClinicalTrials.gov ID: NCT05675007).

**Figure 1.**
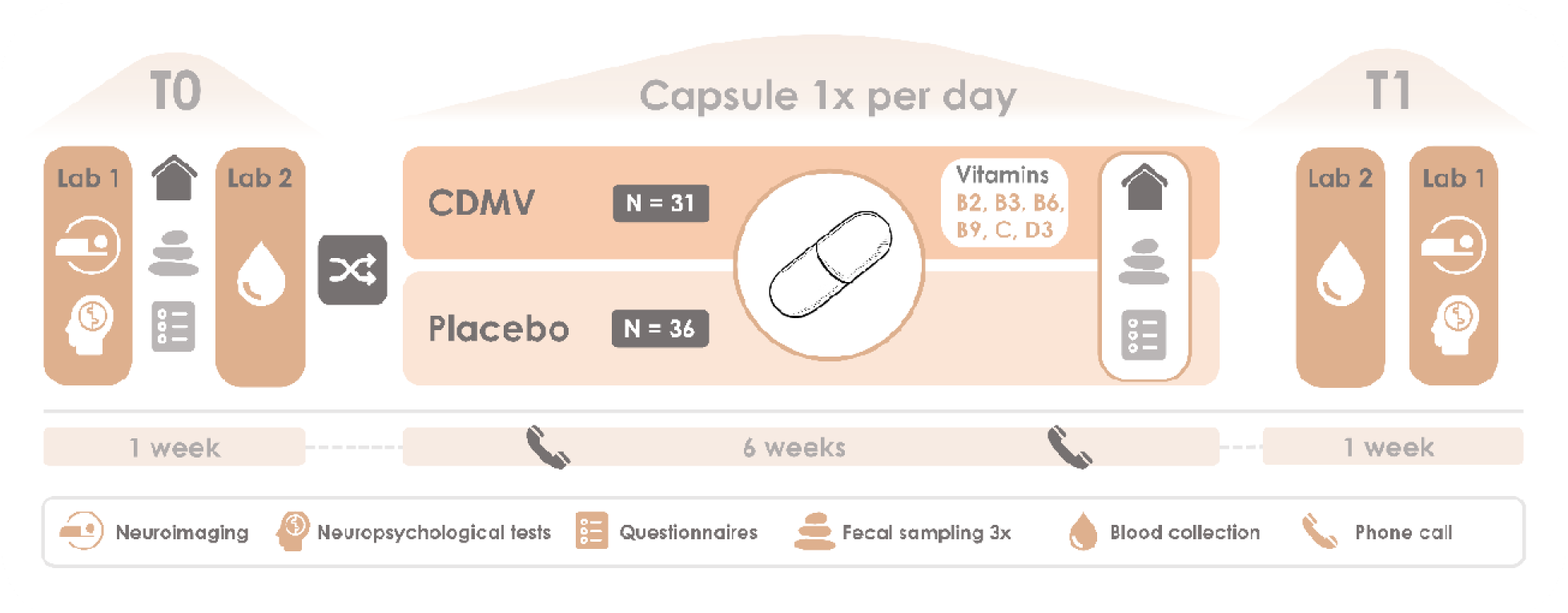
Schematic overview of study visits, measurements, and intervention arms. Before (T0) and after (T1) the intervention period of 6 weeks, participants visited the RU-DCCN (Lab 1) and WUR-HNH (Lab 2) within 1 week. Measurements included neuroimaging and neuropsychological tests (Lab 1), fecal sampling and questionnaires (at home), and blood collection (Lab 2). Participants were randomized into one of the two intervention arms: CDMV (*n* = 31) or placebo (*n* = 36). At week 1 and week 5 of the intervention period, participants received a phone call to assess compliance, check potential problems, and share instructions on the post-intervention fecal sampling (week 5 only). CDMV = colon-delivered multivitamin. *Created with elements from thenounproject.com*

### Participants

In total, 75 Dutch participants aged between 60-75 years enrolled between December 2022 and March 2024, of which 67 completed the trial and were included in the analysis. Participants were recruited via social media, research center participant databases, newsletters of Dutch brain foundations (e.g., Hersenstichting, Alzheimer Nederland), and local newspapers. Individuals that applied for participation were assessed for eligibility via a telephonic screening. Potential participants were included if they were fluent Dutch speakers, and had at least two of the following lifestyle-related risk factors for cognitive decline (50): BMI ≥ 25, physical inactivity as defined by the 2020 WHO guidelines, type 2 diabetes, hypercholesterolemia, (untreated) hypertension, and non-symptomatic cardiovascular disease. Exclusion criteria were: 1) clinical diagnosis of neurological pathology, cerebrovascular events, revascularization surgery in the last 12 months at moment of screening, current malignant diseases, current psychiatric disorders, symptomatic cardiovascular disease, gastro-intestinal diseases or history of gastro-intestinal surgical event, visual impairment, and hearing or communicative impairment, 2) cognitive impairment as determined by Telephone Interview for Cognitive Status (TICS-M1) (51), performed during pre-screening before inclusion and defined as a score <23, 3) use of antibiotics within 3 months before start of the study, 4) use of proton pump inhibitors within the study period, 5) food allergies or other issues with the vitamins present in the supplement, 6) not willing to refrain from taking other supplements (containing vitamin B2, B3, B6, B9, or C, prebiotic, or probiotic) that could interfere with the study outcomes, from at least 2 weeks before start of the intervention till the end of the intervention period, 7) any MRI contraindications, and 8) concurrent participation in other intervention trials. Written informed consent was obtained from each participant during the first study visit.

### Study capsules

Participants consumed a study capsule, either the CDMV supplement or placebo, daily in the morning for six weeks. Study capsules were coated with an Eudragit S 100 layer (52), resulting in targeted release in the colon. Multivitamin capsules contained vitamin B2 (10 mg riboflavin), vitamin B3 (4.0 mg nicotinic acid), vitamin B6 (1.4 mg pyridoxin hydrochloride), vitamin B9 (400 μg folic acid), vitamin C (200 mg ascorbic acid) and vitamin D3 (15ug cholecalciferol 100 SD/S). These doses were selected based on prior evidence (34), safety (53), and technical feasibility. Both placebo and multivitamin capsules were filled with microcrystalline cellulose and magnesium stearate up to 200 mg. Study capsules were identical in appearance, color, taste, smell, and package in order to maintain treatment allocation concealed from participants and researchers. Study capsules jars (seven jars, one for each of the six intervention weeks, and one extra with reserve capsules) were distributed and collected at the WUR-HNH research center. Study capsules were provided by dsm-firmenich.

### Randomization and blinding

This study was performed in a double-blind setting. Participants were randomized 1:1 over the two intervention arms using stratified (2,4) block-randomization. Randomization was stratified by the factors sex (male vs. female), risk factor profile (medium risk 2-3 points vs. high risk ≥4 points) and a combination of age and education level (70+ years or low education vs. 60-69 years and high education, specified as university of applied science associate degree (Dutch: HBO associate degree) or higher). The combined factor of age and education (54) was chosen as (2,4) block-randomization with our sample size allowed for a maximum of three strata. After completion of the baseline visits, participants were randomized automatically in Castor EDC. An independent researcher at the WUR-HNH received a notification each time a participant was randomized, and prepared jars containing the correct study capsules.

### Compliance and safety

Participants completed a daily study diary to track intervention adherence and report potential adverse events. At the post-intervention study visit, participants handed in jars with remaining capsules for compliance verification. Phone calls were scheduled after week one and week five to assess both compliance and potential adverse effects or other intervention issues. Additionally, at pre- and post-intervention study visits, participants completed a brief wellbeing questionnaire to systematically assess adverse events. All reported adverse events were documented and evaluated for severity and potential relatedness to the intervention.

### Study procedures

#### Preparation study visits

The evening before the study visits at the WUR-HNH, participants consumed a standard evening meal provided by the research team, consisting of a pasta wheat meal with cheese and various legumes, containing 610 kJ of energy, 1.70 g of fat, 23.9 g of carbohydrates, 5.30 g of fiber, and 6.00 g of protein per 100 g. After 20:00 hours, the participants were not allowed to eat or drink, except for water, until their blood was collected at the WUR-HNH. Furthermore, participants were asked to refrain from alcohol the day before the study visits.

#### Feces collection

Participants collected the first three consecutive fecal samples after the first study visit at RU-DCCN (T0) and at home after a phone call one week before the WUR-HNH visit (T1) (**Figure 1)**. Feces collection was facilitated by a fecal collection paper (Fecesvanger, Tag Hemi VOF, The Netherlands) and 55 × 44 mm polypropylene containers with scoop-integrated screw caps (Sarstedt, Nümbrecht, Germany). Participants were instructed to sample from three different topographical subsampling locations within the feces. After collection, the samples were stored at -20 °C at home for a maximum of seven days.

The gastrointestinal (GI) transit time was assessed using muffins containing blue dye (55). Each serving of two muffins included 1.5 g of royal blue food coloring (E133 and E122), providing 1753 kJ of energy, 15.9 g of total fat, 63.1 g of carbohydrates, 0.9 g of fiber, and 4.9 g of protein. Participants consumed the muffins during the first study visit at RU-DCCN (T0) or at home after the phone call (T1). During feces collection, participants were instructed to record the date, time of collection, and whether the blue coloring was visible. Participants also assessed stool consistency using the Bristol Stool Scale (BSS). The BSS consists of visualizations of different types of stools, ranging from 1 to 7 with decreasing solidity (56). GI complaints were assessed using 8 items from the validated 15-item gastrointestinal symptom rating scale (GSRS), measured on a 7-point Likert scale where scores ranged from 1 (no symptoms) to 7 (very troublesome symptoms) (57–59). The items evaluated symptoms related to stomach acid, bloating, flatulence, stomach rumbling, constipation, diarrhea, and loose or hard stools. Samples were transported under frozen conditions to the research facility and stored at -80 °C upon arrival.

#### Blood collection

Blood was drawn via a finger-prick for direct measurements, and intravenously from the arm. In total, 30 mL blood was collected: 10 mL in a serum tube, 8 mL in two lithium heparin plasma tubes, 2 mL in a natrium fluoride plasma tube, and 10 mL in four K2EDTA tubes.

#### Neuroimaging data acquisition

Participants were scanned in a Siemens Healthineers Magnetom Skyra 3 Tesla MR scanner at the RU-DCCN, using a 32-channel MR head coil.

##### Anatomical scan

A whole-brain high-resolution T1-weighted anatomical scan was acquired using a magnetization-prepared 2 rapid gradient echo (MP2RAGE) sequence (TR/TE = 6000/2.34 ms; voxel size = 1.0 × 1.0 × 1.0 mm3; FoV = 256 mm; flip angle = 6°; 176 sagittal slices; interleaved slice acquisition; 1.0 mm slice thickness).

##### fMRI acquisition

To examine WM fMRI responses and performance, we used task-related fMRI during a numerical n-back task. The n-back task is a widely-used WM paradigm that requires participants to continuously monitor a series of stimuli (60), numbers ranging from 1-9 in our case. Participants were asked to respond when the presented number was the same as the one presented n trials before. We used a 15-min block design with three conditions (8 blocks per condition): 0-back (control condition, press when number is 1), 1-back (press when number is same as one back), 2-back (press when number is same as two back) (**Figure 4a**). Before performing the actual task within the MR scanner, participants received instructions outside the scanner and had the opportunity to practice the task. Images with blood-oxygen-level-dependent (BOLD) contrast were acquired during the task, using a whole-brain T2*-weighted gradient-echo single-echo multiband acceleration factor 4 echo planar imaging (EPI) sequence (TR/TE = 1500/ 33.40 ms; voxel size 2.0×2.0×2.0 mm; FoV = 213 mm; flip angle = 75°; 68 slices per volume; interleaved slice acquisition).

##### ^1^H-MRS acquisition

Proton magnetic resonance spectroscopy (^1^H-MRS) was used for the quantification of neuroinflammation-related metabolites in the left dlPFC. ^1^H-MRS metabolite spectra were acquired with a Point RESolved Spectroscopy (PRESS) sequence (TR/TE = 2000/35 ms; flip angle = 90°) with chemically selective water suppression (CHESS) (61) in a 20 x 20 x 20 mm isotropic voxel. Water-unsuppressed reference data were acquired next (TR/TE = 2000/35 ms; flip angle = 90°). Advanced shimming was performed before each scan, to minimize field inhomogeneities within the voxel location. A shorter T1 weighted scan was acquired for voxel selection prior to MRS acquisition (TR/TE = 6.3/3.2 ms; voxel size = 1.0 × 1.0 × 1.0mm; FoV = 256 mm; flip angle = 11°). Voxel location in the left dlPFC was performed manually, based on a reference image (see **Supplementary methods Figure M1**). To ensure data quality, minor participant-specific voxel adjustments were made to minimize cerebrospinal fluid and exclude non-brain tissue, accounting for individual anatomical differences in the cortical region. Due to insufficient data quality, hippocampal neuroinflammation indices measured by MRS (pre-registered) are not reported.

##### ASL acquisition

Arterial spin labeling (ASL) was used for non-intravenous quantification of cerebral blood flow (CBF). Arterial blood water in the carotid arteries was labelled using an inversion pulse before the blood flows into the brain. We visualized the carotid arteries using a vessel scout (TR/TE = 47.90/8.15 ms, voxel size = 1.2 × 1.2 × 6.0 mm; flip angle = 9°) and manually placed the inversion pulse plane perpendicular to the feeding arteries according to anatomical landmarks. We acquired perfusion maps, using a Hadamard-encoded multi-PLD pseudocontinuous ASL (pCASL) sequence, and an equilibrium magnetization (M_0_) calibration map. Due to an MR scanner software update and subsequent sequence update in May 2023, we had to change our pCASL sequence during the study period. From December 2022 to May 2023, we used Siemens-based sequences for the pCASL perfusion maps (TR/TE = 4000/16.28 ms, voxel size = 1.8 × 1.8 × 3.5 mm, flip angle = 120°, PLD = 7, repeats = 2) and the M_0_ calibration maps (TR/TE = 3000/16.28 ms, voxel size = 1.8 × 1.8 × 3.5 mm, flip angle = 120°). From May 2023 to May 2024, we used gammaSTAR sequences from the Fraunhofer Institute for Digital Medicine MEVIS for the pCASL perfusion maps (TR/TE = 4000/16.28 ms, voxel size = 1.8 × 1.8 × 3.5 mm, flip angle = 120°, PLD = 7, repeats = 2) and the M_0_ calibration maps (TR/TE = 3000/16.28 ms, voxel size = 1.8 × 1.8 × 3.5 mm, flip angle = 120°). We corrected for potential differences on CBF quantification by including a scanner update covariate in the ASL analyses.

#### Neuropsychological test battery

After neuroimaging, participants underwent a neuropsychological test battery, measuring executive functioning, working memory, episodic memory, and processing speed. The neuropsychological test battery consisted of: the Digit Span Test (DST) (62), the Digit Symbol Substitution Test (DSST) (62), the Rey Auditory Verbal Learning Test (RAVLT) (63), the Trail Making Test (TMT) (64), and the Verbal Fluency Test (VFT) (65).

#### Questionnaires

Participants filled in a short wellbeing questionnaire during the baseline and follow-up visits. Participants were asked not to change their usual dietary habits and health-related behaviors during the period of intervention. To monitor habitual dietary intake over the last month, a food frequency questionnaire (FFQ) was completed before and after the intervention period (66, 67). To assess effects on stress, anxiety, and depression scores, the Perceived Stress Scale (PSS) and Hospital Anxiety and Depression Scale (HADS) were completed before and after the intervention period.

### Fecal laboratory analyses

#### Feces hammering and mill-homogenizing

Fecal samples were processed under frozen conditions in two phases to minimize variability in gut health markers (5). First, fecal samples were crushed into smaller particles (∼0.5-1 cm²) using a dead-blow hammer (Performance Tool Dead-Blow Hammer, Wilmar Corporation) while kept frozen with dry ice and liquid nitrogen. The hammered samples were then stored at -80°C. Second, a portion of the crushed feces was further processed using the IKA® Tube Mill 100 control (IKA-Werke GmbH & Co. KG, Staufen, Germany). Approximately 10 grams of the crushed sample was placed in a 40 mL disposable milling chamber (art. IKAA20001173, IKA) along with two pieces of dry ice. Feces were milled under frozen conditions for approximately 20 sec at 25000 rpm, pulverizing it into a fine, homogeneous powder. The homogenized fecal samples were stored at -80°C until further analysis.

#### Fecal fatty acids

Fecal SCFAs (acetic acid, propionic acid, butyric acid, valeric acid), medium-(M)CFAs (heptanoic acid, hexanoic acid) and branched-(B)CFAs (isobutyric acid, isovaleric acid, 4-methylvaleric acid) levels were quantified using gas chromatography with flame ionization detection (GC-FID) using an Agilent 8690 GC system (Agilent, Amstelveen, the Netherlands) equipped with a Restek Stabilwax-DA column (Restek Corporation, United States), as previously described (68). Briefly, 200 mg of mill-homogenized feces was weighed and mixed with 1 mL of ultra-high performance liquid chromatography mass spectrometry water (ULC/MS). The samples were put on a shaker for 20 min (Universal shaker SM-30, Edmund Bühler GmbH, Bodelshausen, Germany) and centrifuged for 5 min at 9000 g. Subsequently, 500 µL of the supernatant was transferred to a new tube and combined with 250 µL of internal standard solution. This internal standard contained 0.45 mg/ml 2-ethylbutyric acid (Aldrich, 109959), 0.3 M HCl (VWR, 20252.244) and 0.9 M oxalic acid (Merck, 1.00495.0500), all dissolved in ULC/MS-grade water. A quality control reference mix was prepared using 50 µL ready-made volatile free acid mixture (CRM46975, Sigma-Aldrich), 450 µL ULC/MS water, and 250 µL internal standard solution. Additionally, a standard stock solution containing 0.45 mg/mL each of acetic acid (Biosolve, 0001074131), propionic acid (Aldrich, 402907), isobutyric acid (Aldrich, 58360), butyric acid (Aldrich, 103500), isovaleric acid (Aldrich, 129542), valeric acid (Aldrich, 240370), 4-methyl valeric acid (Aldrich, 277827), hexanoic acid (Aldrich, 153745), heptanoic acid (Aldrich, W334812) was prepared in ULC/MS-water. From this stock solution, a five-point dilution series (1:1 dilutions) was generated, producing six standard solutions with concentrations from 0.045 to 0.45 mg/mL. 500 µL of each standard solution was mixed with 250 µL of the internal standard solution. All samples, controls, and standards were vortexed, centrifuged at 9000g for 5 min, and 150 µL of the supernatant was transferred to GC injection vials, of which 2 µl was injected into the GC-FID. The GC run settings were as follows: (i) 1 min at 80 °C, (ii) raised by 10 °C /min to 200 °C, (iii) hold time of 5 min, (iv) raised by 25 °C/min, to a temperature of 230 °C, (v) 5 min hold time. The carrier gas was nitrogen at a flow rate of 1.32 mL/min. Data were analyzed using Openlabs CDS software version 2.6 (Agilent Technologies, Santa Clara, California, USA). SCFA concentrations were expressed as μmol/g of wet weight using the formula: ((compound concentration (mg/mL)/sample weight (g))/molar mass acid (g/mol))*1000.

#### Fecal microbiota composition

Bacterial DNA was extracted from approximately 250 mg feces using the DNeasy® PowerSoil® Pro Kit (QIAGEN, Hilden, Germany), following the manufacturer’s instructions. Cell lysis and bead beating were performed using the TissueLyser adapter set 2 (QIAGEN, Hilden, Germany). DNA concentrations were quantified using a NanoDrop ND-1000 spectrophotometer (Thermo Fisher Scientific, Waltham, MA, USA). The extracted DNA was used to prepare libraries for sequencing with 2× Phanta Max Master Mix (VAZYME, China) polymerase. The V3–V4 variable region of bacterial 16S rDNA was amplified by PCR using the primers 338F: 5’-ACTCCTACGGGAGGCAGCAG-3’ and 806R: 5’-GGACTACHVGGGTWTCTAAT-3’. PCR enrichment was performed in a 50 µL reaction containing 30 ng of DNA template and fusion PCR primers. The PCR cycling conditions were as follows: 95°C for 3 min; 30 cycles of 95°C for 15 sec, 56°C for 15 sec, 72°C for 45 sec and final extension at 72°C for 5 min. PCR products were purified by DNA magnetic beads (BGI, LB00V60). The purified products were used to generate single-stranded library products through denaturation, followed by circularization to produce single-strand circular DNA molecules. Single strand linear DNA was removed using digestion. The final single strand circularized library is amplified with phi29 and rolling circle amplification (RCA) to generate DNA nano balls (DNB) carrying multiple copies of the initial single stranded library molecule. DNBs were loaded into the patterned nanoarray. Sequencing reads of PE300 bases length were generated with the DNBSEQ-G400 platform (BGI-Shenzhen, China), with a coverage of 100k reads (clean tags per sample). Raw sequencing data were processed using the QIIME2R-DADA2 workflow (69), an amplicon sequence variant (ASV)-based approach. Taxonomy was assigned using the SILVA database version 138.2 (https://www.arb-silva.de/documentation/release-1382).

#### Other fecal measurements

Other fecal measurements, including feces characteristics (water content, pH, and redox potential) and fecal inflammatory markers (calprotectin, lipocalin-2 (LCN-2), and secretory immunoglobulin A (sIgA)) can be found in **Supplementary Methods**.

### Blood laboratory analyses

#### Blood vitamins

To validate colon-delivery and detect potential absorption, concentrations of vitamins present in the supplement were measured in blood. Blood tubes (Vacutainers, BD) containing EDTA (K2E) were used to analyze vitamin B2, B3 and B6. Tubes were inverted eight times, and processed as follows: 500 µL of whole blood for vitamin B2 and B6 analyses was stored at -20°C until analysis, while 2 mL of whole blood for vitamin B3 analysis was kept at 2–8°C and analyzed within 48 hours. Vitamin B2, B3, and B6 levels were measured chromatographically using HPLC with spectrophotometry fluorescence detection at a medical laboratory (Reinier Medical Diagnostic Center, Delft, The Netherlands). Blood samples for vitamin C analysis were collected in lithium-heparin tubes (Vacutainers, BD), inverted eight times, and centrifuged at 3000 × g for 8 min at 20°C. 500 µL of plasma was mixed with 2 mL of pre-cooled metaphosphoric acid. The mixture was stored at 2-8°C and analyzed within 48 hours using HPLC and spectrophotometry UV detection at the same laboratory (Reinier Medical Diagnostic Center, Delft, The Netherlands). For vitamins B9 and D, serum collection tubes (Vacutainers, BD) were used. These tubes were inverted eight times, left to clot for a minimum of 30 min, and afterwards centrifuged (3000 × g) for 8 min at 20°C. For vitamin B9, 500 µL serum was analyzed at the Clinical Chemistry and Hematology Laboratory of Gelderse Vallei Hospital (Ede, The Netherlands) using standard clinical laboratory assays. For vitamin D, including 25-hydroxyvitamin D3 (25OHD3) and 25-hydroxyvitamin D2 (25OHD2), 600 µL of serum was analyzed at the Clinical Chemistry and Hematology Laboratory Dicoon (Canisius-Wilhelmina Ziekenhuis, Nijmegen, The Netherlands), as described previously (70).

#### Other blood measurements

Other blood laboratory measurements, including blood SCFAs, MCFAs and BCFAs, intestinal permeability markers (lipopolysaccharide-binding protein (LBP) and zonulin), and systemic inflammation markers (high-sensitivity C-reactive protein (hs-CRP), total white blood cell (WBC) count, interferon-γ (IFN-γ), tumor necrosis factor-α (TNF-α), interleukin (IL)-6, IL-8, and IL-10) can be found in **Supplementary Methods**. Blood antioxidant status profile, metabolic profile, and brain health profile were preregistered on clinical trial registration, but eventually not measured.

### Neuroimaging data analyses

#### Working memory fMRI responses and performance (n-back task)

##### Preprocessing & Quality control

Anatomical and fMRI data were preprocessed using fMRIPrep (23.2.0; RRID:SCR_016216) (71). Functional scans were realigned, coregistered to the participant’s T1-weighted anatomical scan, susceptibility distortion corrected, and spatially normalized to MNI152 space. A detailed description of all preprocessing steps with fMRIPrep can be found in the **Supplementary methods**. We used Statistical Parametric Mapping 12 (SPM12; https://www.fil.ion.ucl.ac.uk/spm/software/spm12/) running in MATLAB R2024a (Mathworks Inc.; https://nl.mathworks.com/products/matlab.html) to spatially smooth the preprocessed BOLD time series with a 6 mm FWHM kernel. We assessed the quality of individual fMRI datasets and task performance before including participants in the group analysis. Participants that obviously misunderstood the n-back task, indicated by 1) a target accuracy lower than 30% for the 0-back condition and/or 2) a non-target accuracy lower than 65% for the 0-back, 1-back or 2-back condition, were excluded from analysis. This led to exclusion of *n*=2 datasets. We evaluated fMRI dataset quality using the fMRIPrep reports and motion parameters. For datasets that were deemed of doubtful quality (framewise displacement mean >0.5 mm or max >3 mm), the first level 2-back – 0-back map (thresholded at p=0.001 uncorrected) was visually inspected for frontoparietal activation typical for WM by two researchers, blinded for intervention allocation. In case the map was lacking any frontoparietal activation, the dataset was excluded. This led to inspection of 18 datasets and exclusion of *n*=3 participants, for whom one of the two fMRI datasets did not have sufficient quality. In addition, for *n*=6 participants fMRI data was lacking for one of the sessions due to scanner issues. As a result, *n*=62 participants were included for n-back performance analysis, of which *n*=56 participants were included for n-back fMRI responses analysis.

##### First level analysis

fMRI analyses were performed in SPM12. For each participant, we used one general linear model (GLM) to model block-related BOLD responses for both the pre- and post-intervention session. Per session we had three conditions (0-back, 1-back, 2-back), resulting in six unique task parameters for each session-condition combination. In addition to these task parameters, we included session-specific confound regressors in the model: framewise displacement, six realignment parameters, six anatomical principal component noise regressors (aCompCor), six temporal principal component noise regressors (tCompCor), all cosine regressors, and all independent components labeled as noise by Independent Component Analysis-based Automatic Removal Of Motion Artifacts (ICA-AROMA). ICA-AROMA regressors were retrieved from an earlier version of fMRIPrep (23.0.2; RRID:SCR_002502) and included as regressors instead of using AROMA-denoised data, to prevent removal of shared variance between noise regressors and task parameters. ICA-AROMA regressors that had a covariation higher than 5% with the task design were removed from the model. For our block design, we modeled each experimental condition as a separate regressor by convolving the duration of each block with the SPM standard hemodynamic response function. Instead of using a high-pass filter, we controlled for low-frequency drift through cosine regressors in the model. We used 2-back – 0-back as primary WM contrast. We created this contrast pre-intervention (2-back pre – 0-back pre), post-intervention (2-back post – 0-back post), and subtracted ([2-back post – 0-back post] – [2-back pre – 0-back pre]) to model the effect of time on WM at the first level. Statistical parametric difference maps per contrast were generated for each participant.

##### ROI analysis

Independent ROI masks were created for the dlPFC (**Figure 4d**) and hippocampus (**Figure 4f**). The dlPFC mask was based on dlPFC regions in the WM activation map retrieved from a meta-analysis of 1091 studies (Neurosynth; https://neurosynth.org/analyses/terms/working) overlaid with peak dlPFC voxels of aging-related n-back effects retrieved from a meta-analysis of 96 studies (72). The hippocampus mask was retrieved from the Harvard-Oxford subcortical atlas (73). Mean ROI beta values were extracted from first level pre- and post-intervention statistical parametric maps using SPM12, and used for the group analysis.

##### Whole-brain analysis

We also performed whole-brain group analyses using a two-sample T-test. Within the second level model, the neurocognitive covariates reported within the **Statistical analyses** section below were used. We explored results at a significance cluster-defining threshold of p<0.001, uncorrected for multiple comparisons.

##### gPPI analysis

In addition to the fMRI activation analysis, we assessed task-related functional connectivity between the dlPFC and hippocampus by conducting a generalized psychophysiological interaction (gPPI) analysis. An 8 mm sphere in the dlPFC was used as seed region. Using our primary WM contrast (2-back – 0-back), we searched for the average peak activation voxel across all participants and all sessions within the dlPFC (BA9) and selected it as center of the sphere (x=40, y=36, z=32) (**Figure 5d**). gPPI analyses were performed in SPM12 using the Generalized PPI Toolbox (v13; RRID:SCR_009489). For each participant, we combined the pre- and post-intervention session in one GLM to predict the seed region’s BOLD response with psychological, physiological, and psychophysiological interaction (PPI) regressors. The following regressors were included in the model: six psychological task regressors for each unique session-condition combination, two physiological regressors derived from the first eigenvariate of the BOLD signal in the dlPFC seed region for each session, and six condition-specific PPI terms. These PPI terms were generated by multiplying each psychological regressor with the deconvolved physiological signal, followed by reconvolution with the canonical hemodynamic response function. We kept the other steps within our first level gPPI model (confound regressors, contrasts) similar to the fMRI activation first level model described above. Mean ROI beta values for the hippocampus were extracted from first level pre- and post-intervention statistical parametric maps using SPM12, and used for the group analysis. In addition, we also explored whole-brain effects in a second level model, as described above.

##### Performance

We used the dprime as a measure of n-back performance for each condition. The dprime [d’ = z(H) – z(F)] calculates the difference between z-scores of hit rate (H, correct detections) and false alarm rate (F, false alarms). Higher d’ values indicate better discrimination between signal and noise, independent of response bias.

#### Other neuroimaging analyses

Other neuroimaging data analyses, including determination of neuroinflammatory metabolites using ^1^H-MRS and cerebral blood levels flow using ASL, can be found in **Supplementary Methods**.

### Statistical analyses

#### Sample size

Sample size was based on a combination of previous trials (available when the study was designed in 2021) using a similar gut intervention, target group, or measuring method, and assessing effects on one of our primary outcomes: SCFAs, WM performance or fMRI responses (34, 43, 74, 75). Based on these previously published effect sizes, we calculated that at least 60 participants (30 per arm) would be needed to reach a power of 80% with a significance level of 0.05. Accounting for 15% data loss in neuroimaging studies, a total of 70 participants had to be included. We only analyzed complete cases and did not use drop-out data, according to the per protocol principle. Drop-outs were replaced to a maximum of 10% of the participants, leading to a maximal allowed inclusion number of 76 participants.

#### Group analyses

To assess group effects, a linear model predicting post-intervention values was performed in R (version 4.4.0), using the lm function from the stats package. The basic linear model included baseline value and intervention group as covariate: lm(“post value” ∼ “pre value” + ‘’intervention group”). In the final linear model, age and sex (76) were added as covariates, along with other covariates depending on the outcome measurement group (gut, immune, or neurocognitive). For gut outcomes, habitual dietary fiber intake (from FFQ reflecting last month at T1) (77) and BMI (78) were added as covariates. For immune outcomes, BMI (79) was added as covariate. For neurocognitive outcomes, education group (80) was added as covariate. Additionally, we controlled for potential systemic effects, should any unexpected vitamin absorption occur. Specifically, if post-intervention blood concentrations of any vitamin present in the supplement were significantly elevated in the multivitamin group compared to placebo, that vitamin’s post-intervention level was included as a covariate in neurocognitive outcome models to ensure that observed effects were attributable to colon-targeted mechanisms.

Data validity was assessed through comprehensive quality checks for neuroimaging data (explained in detail for each neuroimaging outcome) and the interquartile range (IQR) method for identifying outliers in all other outcomes. Outliers were retained in analyses unless there were serious reasons to suspect erroneous data (e.g., substantial protocol deviations or abnormalities reported on participant wellbeing questionnaire). Missing data for primary and secondary outcomes were not imputed. Analyses were conducted separately for each outcome measure, including only participants with available and valid pre- and post-intervention data for that specific outcome. Sample sizes for each outcome analysis are specified in the corresponding tables and figures. For selected covariates, we performed imputation where we deemed this methodologically sound. This decision was made to maximize statistical power and prevent exclusion of participants solely due to missing covariate data. Missing values were present for covariates habitual dietary fiber intake (*n*=6) and blood vitamin B6 (*n*=1). Imputation was applied by substituting missing values with the participant’s baseline value (*n*=5 habitual fiber intake), as habitual dietary fiber intake was not expected to change over the course of the 6-week trial. In case the baseline value was also missing, we used the mean end value of all participants of the same sex (*n*=1 habitual fiber intake), or of the same sex within the same intervention group (*n*=1 blood vitamin B6).

Reported p-values represent between-group differences at week 6, adjusted for baseline values and – for the final linear models – also for other covariates. The normality of residuals was assessed using Q-Q plots, and homoscedasticity (equal variance) was evaluated by plotting residuals against the fitted values. When residuals deviated from normality, data were logarithmically transformed. In these cases, back-transformed adjusted means were reported as well, in addition to logarithmic adjusted means.

#### Gut-brain correlations analysis

To explore gut-brain relationships, correlations between changes over time (Δ post-pre) of primary gut- and neurocognitive outcomes were assessed. Delta values were corrected for baseline value, and residuals were used to create a false discovery rate (FDR) corrected Spearman correlation heatmap. Correlations of interest were further inspected. Additionally, changes over time (Δ post-pre) correlations between primary neurocognitive outcomes and secondary gut outcomes showing significant group effects were explored in a similar way.

#### Microbiota composition and diversity analyses

Microbiota analyses were performed in R (version 4.4.0). Archaeal sequences, chloroplasts, mitochondria, singletons and doubletons were filtered from the dataset. Only bacterial taxa with a relative abundance exceeding 0.25% were retained for downstream analyses (81). Alpha-diversity (within-sample diversity) metrics, including Shannon Diversity (species richness and evenness), Phylogenetic Diversity (diversity accounting for the evolutionary relationships among species), and Chao1 (species richness), were calculated based on non-rarefied ASV counts using the microbiome and picante R packages. Beta-diversity (between sample diversity) was calculated by Bray-Curtis dissimilarity, as implemented in the phyloseq package, on relative abundances of ASVs to determine overall microbiota differences between groups over time. Differences were visualized using Principal Coordinate Analysis (PCoA). For microbiota alpha-diversity, the linear model, as described above, was also applied. To test for differences in the overall microbiota community composition between groups over time, permutational multivariate analysis of variance (PERMANOVA) was performed on the Bray-Curtis distances accounting for repeated sampling within individuals using the vegan package. For microbiota differential abundance testing on genus level, MaAsLin3 (Microbiome Multivariable Associations with Linear Models) was applied, which is specifically developed for highly sparse and compositional data (82). A prevalence filter was applied ensuring that the bacteria on genus level were present in >10% of samples. MaAsLin3 fitted linear regression models on non-rarefied, log-transformed relative abundances for differential abundance testing, as well as logistic regression models for prevalence testing. The basic MaAsLin3 model included the interaction between group and time, sequencing read depth as a covariate, and a random intercept for individuals. The extensive MaAsLin3 model also included sex, age, BMI, habitual dietary fiber intake, sequencing read depth, the interaction between group and time, and a random intercept for individuals. Multiple testing correction was performed using the Benjamini-Hochberg FDR.

#### Other analyses

For neuropsychological test battery analysis see **Supplementary Methods**.

## Results

### Baseline characteristics

A total of 187 volunteers were recruited and screened, of which 75 individuals were included in the study. After one drop-out during the baseline visit, 74 participants were randomly allocated to an intervention arm (35 received CDMV supplements and 39 received placebo). During the intervention period, seven participants dropped out due to various reasons not related to the study intervention (see **Figure 2** for more details), leading to a total drop-out number of eight participants. A total of 67 participants completed the study and were included in the analysis, of which 31 were in the CDMV group and 36 in the placebo group (**Figure 2**). The mean age of our study population was 65.3 ± 3.9 years and 60% of the participants were female. Baseline characteristics of the study population per intervention arm can be found in **Table 1**.

**Figure 2.**
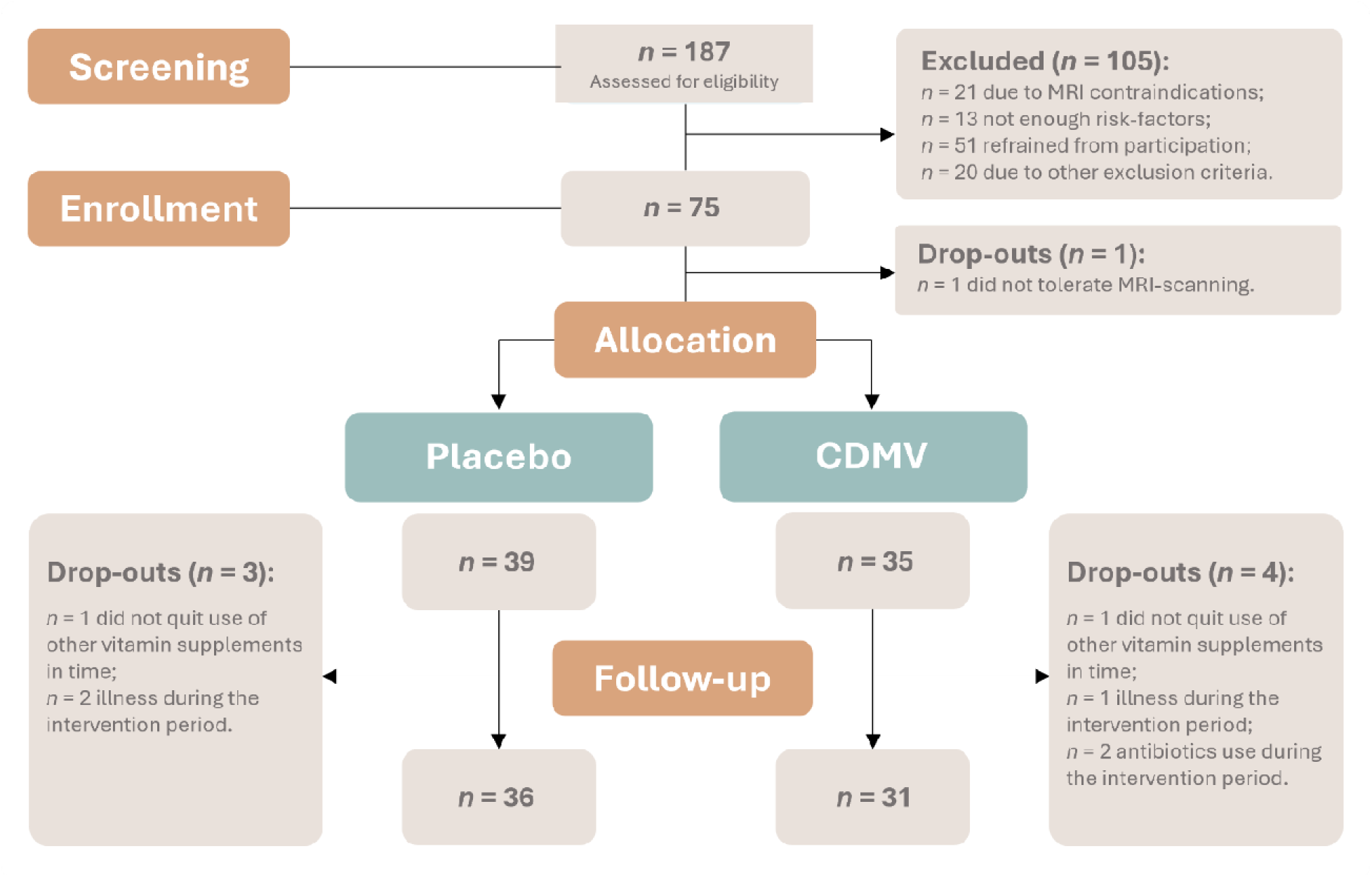
CONSORT flow diagram. CONSORT flow diagram with screening, enrollment, allocation, and follow-up participant numbers. CDMV = colon-delivered multivitamin.

**Table 1.**
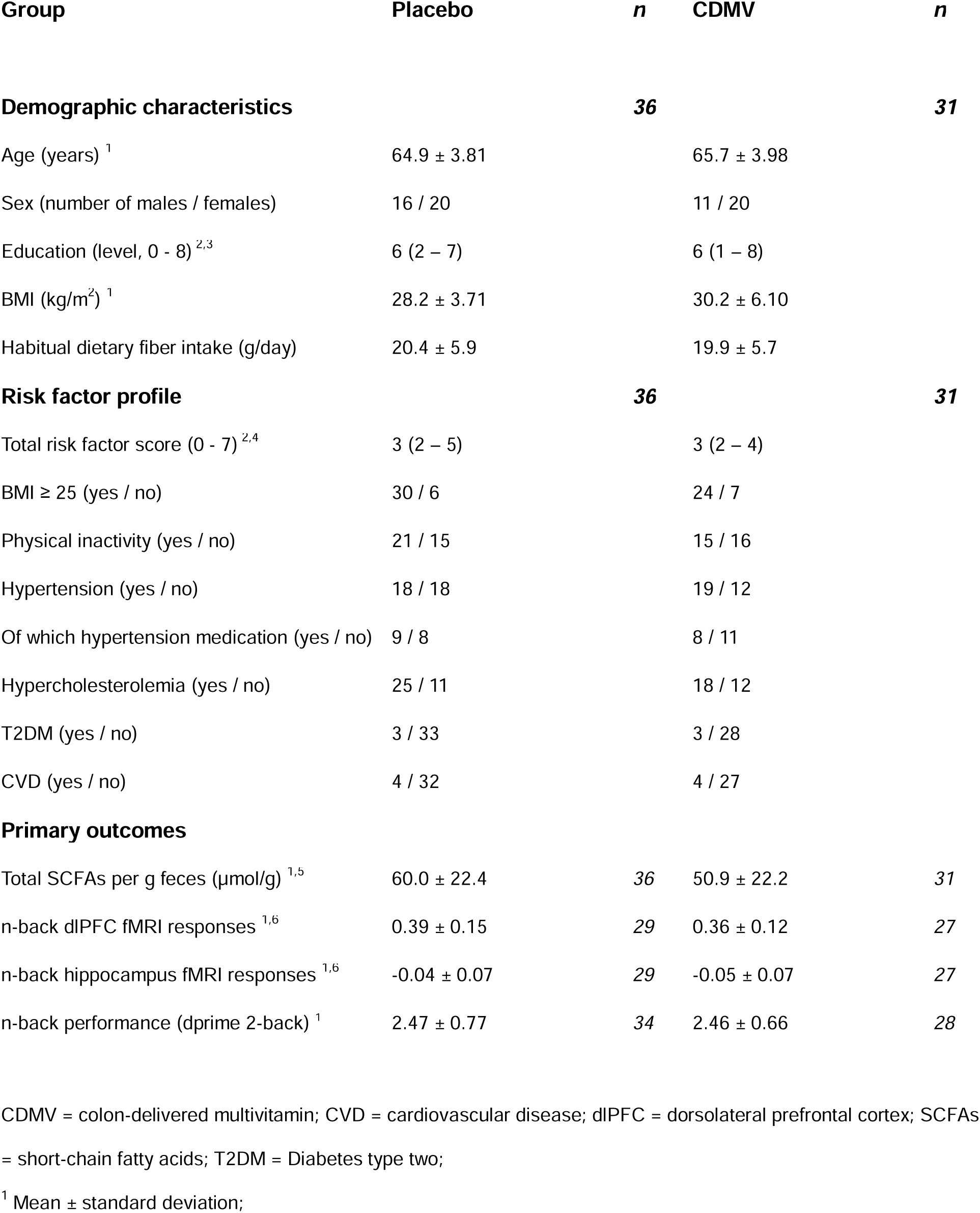
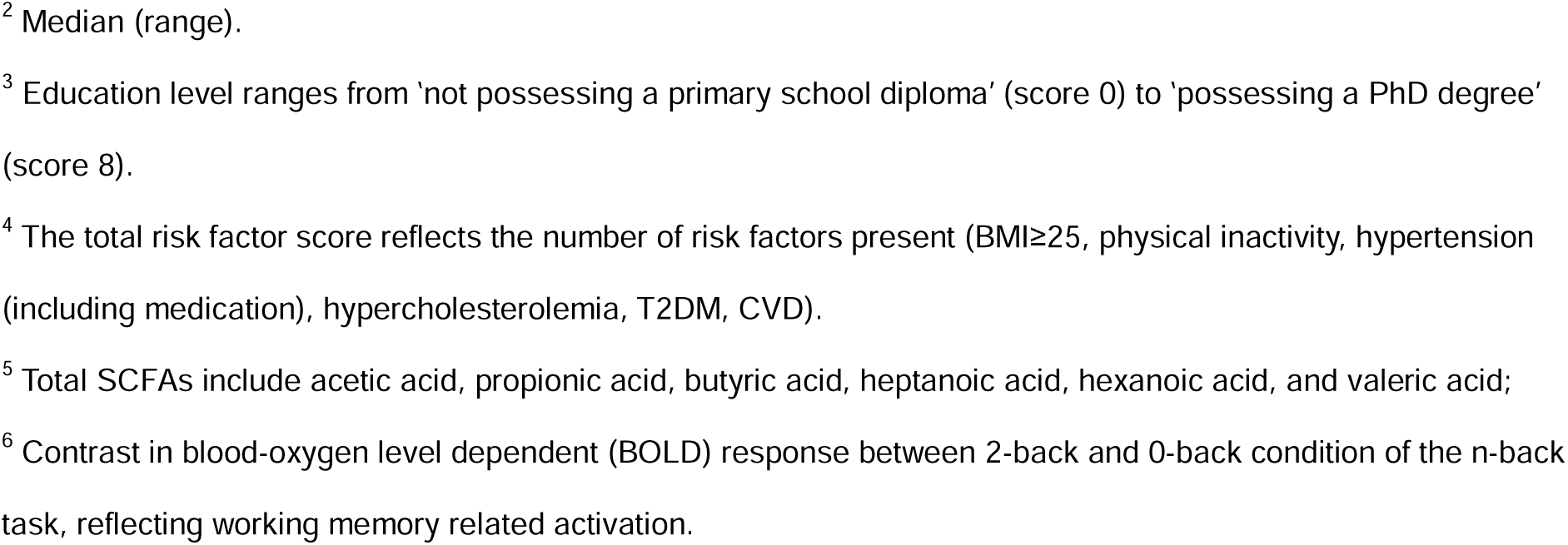
Baseline demographics of the COMBI participants.

| Group | Placebo | *n* | CDMV | *n* |
| --- | --- | --- | --- | --- |
| **Demographic characteristics** |  | **36** |  | **31** |
| Age (years) <sup>1</sup> | 64.9 ± 3.81 |  | 65.7 ± 3.98 |  |
| Sex (number of males / females) | 16 / 20 |  | 11 / 20 |  |
| Education (level, 0 - 8) <sup>2,3</sup> | 6 (2 – 7) |  | 6 (1 – 8) |  |
| BMI (kg/m<sup>2</sup>) <sup>1</sup> | 28.2 ± 3.71 |  | 30.2 ± 6.10 |  |
| Habitual dietary fiber intake (g/day) | 20.4 ± 5.9 |  | 19.9 ± 5.7 |  |
| **Risk factor profile** |  | **36** |  | **31** |
| Total risk factor score (0 - 7) <sup>2,4</sup> | 3 (2 – 5) |  | 3 (2 – 4) |  |
| BMI ≥ 25 (yes / no) | 30 / 6 |  | 24 / 7 |  |
| Physical inactivity (yes / no) | 21 / 15 |  | 15 / 16 |  |
| Hypertension (yes / no) | 18 / 18 |  | 19 / 12 |  |
| Of which hypertension medication (yes / no) | 9 / 8 |  | 8 / 11 |  |
| Hypercholesterolemia (yes / no) | 25 / 11 |  | 18 / 12 |  |
| T2DM (yes / no) | 3 / 33 |  | 3 / 28 |  |
| CVD (yes / no) | 4 / 32 |  | 4 / 27 |  |
| **Primary outcomes** |  |  |  |  |
| Total SCFAs per g feces (μmol/g) <sup>1,5</sup> | 60.0 ± 22.4 | 36 | 50.9 ± 22.2 | 31 |
| n-back dlPFC fMRI responses <sup>1,6</sup> | 0.39 ± 0.15 | 29 | 0.36 ± 0.12 | 27 |
| n-back hippocampus fMRI responses <sup>1,6</sup> | -0.04 ± 0.07 | 29 | -0.05 ± 0.07 | 27 |
| n-back performance (dprime 2-back) <sup>1</sup> | 2.47 ± 0.77 | 34 | 2.46 ± 0.66 | 28 |

CDMV = colon-delivered multivitamin; CVD = cardiovascular disease; dlPFC = dorsolateral prefrontal cortex; SCFAs
= short-chain fatty acids; T2DM = Diabetes type two;
<sup>1</sup> Mean ± standard deviation;
<sup>2</sup> Median (range). <sup>3</sup> Education level ranges from 'not possessing a primary school diploma' (score 0) to 'possessing a PhD degree' (score 8). <sup>4</sup> The total risk factor score reflects the number of risk factors present (BMI≥25, physical inactivity, hypertension (including medication), hypercholesterolemia, T2DM, CVD). <sup>5</sup> Total SCFAs include acetic acid, propionic acid, butyric acid, heptanoic acid, hexanoic acid, and valeric acid; <sup>6</sup> Contrast in blood-oxygen level dependent (BOLD) response between 2-back and 0-back condition of the n-back task, reflecting working memory related activation.

### Compliance and user experiences

The CDMV group showed a consumption compliance of 99.8%, and the placebo group demonstrated a consumption compliance of 99.5%. In both groups, instances of missed capsules occurred sporadically across different participants. Participants complied with instructions to maintain their habitual diet, as dietary intake remained comparable between groups throughout the intervention period, with no significant between-group differences (**Supplementary table 3** and **Supplementary figure 1**).

Within the CDMV group, 16.7% of the participants experienced side effects and 20.0% of the participants experienced effects on stool and/or intestines. Within the placebo group, 11.1% of the participants experienced side effects and 19.4% of the participants experienced effects on stool and/or intestines. These numbers are comparable between both intervention groups. For more user experiences, see **Supplementary table 5** and **Supplementary figure 3**. No serious adverse effects (SAEs) occurred during the study. For a summary of all AEs, see **Supplementary Data**.

### Effects of colon-delivered multivitamin supplementation on fecal SCFAs

No significant between-group differences were found for total fecal SCFAs (b=5.38 (CI - 4.95; 15.7), p=0.30) or individual fecal SCFAs (**Table 2** and **Figure 3**). In other words, the CDMV group did not show the expected increase in fecal SCFAs compared to the placebo group.

**Figure 3.**
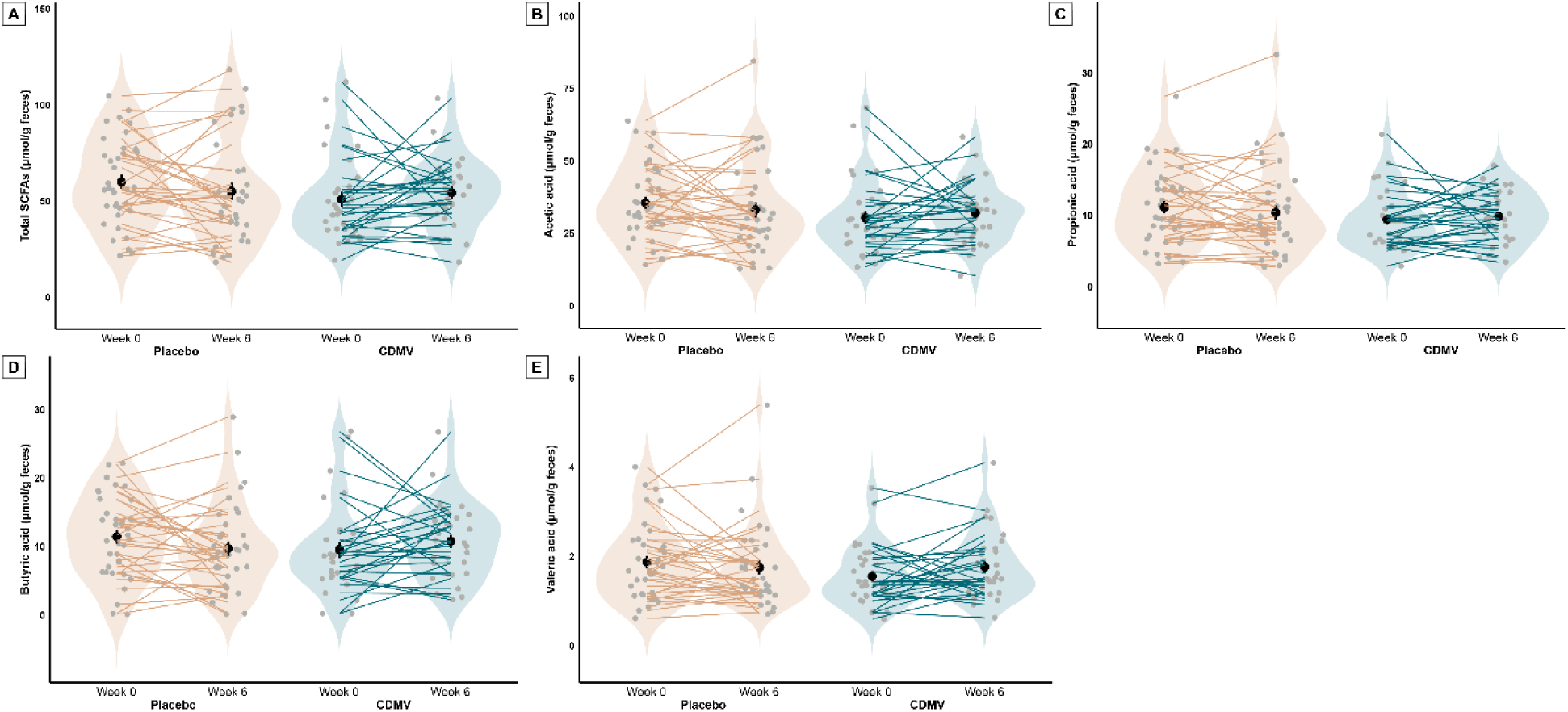
Fecal SCFA levels at baseline and after the 6-weeks intervention for placebo (*n*=36) and CDMV group (*n*=31). Total SCFAs (**A**), acetic acid (**B**), propionic acid (**C**), butyric acid (**D**), and valeric acid (**E**) levels are reported. Individual paired samples are connected by a line. The width of the violin shapes indicates the sample density, the circled shape inside indicates the group mean and SE. CDMV = colon-delivered multivitamin; SCFAs = short-chain fatty acids.

**Table 2.** Intervention-induced effects in placebo and CDMV group on the fecal short-chain fatty acids of older adults at risk of cognitive decline.

| Outcome | Placebo | | CDMV | | Estimate group (CI) | Partial $\eta^2$ | P-value |
| --- | --- | --- | --- | --- | --- | --- | --- |
|  | Mean<sub>adj</sub> (SE) | n | Mean<sub>adj</sub> (SE) | n |  | (CI) |  |
| <u>Total fecal short-chain fatty acids</u><br><u>(μmol/g feces)</u> | 54.5 (3.48) | 36 | 59.9 (3.79) | 31 | 5.38<br>(-4.95; 15.7) | 0.01<br>(0; 0.11) | 0.30 |
| Acetic acid (μmol/g feces) | 32.1 (2.03) | 36 | 34.7 (2.21) | 31 | 2.54<br>(-3.56; 8.63) | 0.00<br>(0; 0.09) | 0.41 |
| Propionic acid (μmol/g feces) | 10.1 (0.68) | 36 | 10.9 (0.74) | 31 | 0.821<br>(-1.22; 2.86) | 0.00<br>(0; 0.09) | 0.42 |
| Butyric acid (μmol/g feces) | 9.52 (0.90) | 36 | 11.6 (0.99) | 31 | 2.06<br>(-0.650; 4.77) | 0.03<br>(0; 0.16) | 0.13 |
| Valeric acid (μmol/g feces) * | 0.19 (0.03) | 36 | 0.24 (0.03) | 31 | 0.06<br>(-0.03; 0.14) | 0.02<br>(0; 0.14) | 0.17 |
|  | 1.58 (0.10) |  | 1.81 (0.12) |  |  |  |  |

CDMV = colon-delivered multivitamin. Estimated marginal means with standard error at week 6 are reported for the final linear model (adjusted for baseline value, age, sex, BMI and habitual dietary fiber intake). Parameter estimate and confidence interval for CDMV group are reported. Partial $\eta^2$ and confidence interval are reported as measure of effect size; lower CI capped at theoretical minimum (0) and upper CI capped at theoretical maximum (1). Primary outcomes are underlined, full linear model statistics of primary outcomes can be found in **Supplementary table 1**. Outcomes indicated with an asterisk (\*) are log-transformed (log10) for normal distribution; original (upper) and back-transformed (lower) adjusted means and standard errors are reported for these outcomes. Results for the basic model (adjusted for baseline value only) can be found in **Supplementary table 2**. Non-adjusted means at week 0 and week 6 can be found in **Supplementary** **table 10**.

#### Other gut and immune-related outcomes

Microbiome analysis at genus level showed that three genera were significantly higher post-intervention in the CDMV group compared to placebo: *Lachnospiraceae FCS020 group* (b=9.07, SD=2.32, p=0.03), *Eubacterium ventriosum group* (b=3.98, SD = 0.02, p=0.03), and *Ruminococcus gauvreauii group* (b=1.52, SD<0.001, p<0.001) (**Supplementary table 11**, **Supplementary figure 14**). We found no significant between-group effects on fecal microbiota alpha diversity (phylogenetic diversity, Shannon diversity, and Chao1). In addition, no clustering effect was found between pre- and post-intervention within and between groups based on the overall microbiota profiles based on Bray-Curtis dissimilarity beta-diversity (PERMANOVA p=0.83, **Supplementary figure 13**). This absence of pre-post clustering was comparable between the CDMV and placebo group, indicating that the observed variation reflected natural temporal fluctuations rather than intervention-induced changes in microbiota composition.Complete microbiota composition changes per group can be found in **Supplementary table 12.**

Post-intervention fecal levels of the MCFA hexanoic acid were significantly higher in the CDMV group compared to placebo (b=0.15 (CI 0.00; 0.29), p=0.04, ηp²=0.07). No significant group effects were found on other gut- and immune-related secondary outcomes, including other fecal MCFAs and BCFAs, feces characteristics (water content, pH, and redox potential), fecal intestinal inflammation markers (LCN-2; due to technical issues calprotectin and sIgA have too many missing values and are not reported), blood SCFAs, MCFAs and BCFAs, blood intestinal permeability markers (zonulin and LBP), blood systemic inflammation markers (hs-CRP, total WBC count, IFN-γ, TNF-α, IL-6, IL-8, and IL-10), stool consistency, and gut transit time (see **Supplementary table 6 and 7,** and **Supplementary figures 4, 5, 6, 7, 8, and 9**).

### Effects of colon-delivered multivitamin supplementation on working memory-related fMRI responses and performance

Among the vitamins measured in blood, vitamin B6 was the only vitamin that showed significant between-group differences post-intervention, and increased in the CDMV group (see **Supplementary table 4** and **Supplementary figure 2**). Therefore, blood vitamin B6 was added as covariate in all neurocognitive analyses.

First, we checked whether the WM task showed the predicted behavioral and BOLD responses (across participants, intervention groups, and sessions). As expected, overall n-back performance (dprime) was significantly lower for the 2-back condition compared to the 1-back and 0-back conditions (p<0.001) (**Figure 4b**). The main overall n-back task-related brain activity map for the 2b-0b contrast elicited the predicted fronto-parietal responses, as shown in **Figure 4c**. In line with these whole-brain responses, the dlPFC fMRI responses were highest for the 2b-0b contrast, compared to the 1b-0b and 2b-1b contrasts (**Figure 4d-e**). Hippocampal fMRI responses were comparable for all three n-back contrasts (**Figure 4f-g**).

**Figure 4.**
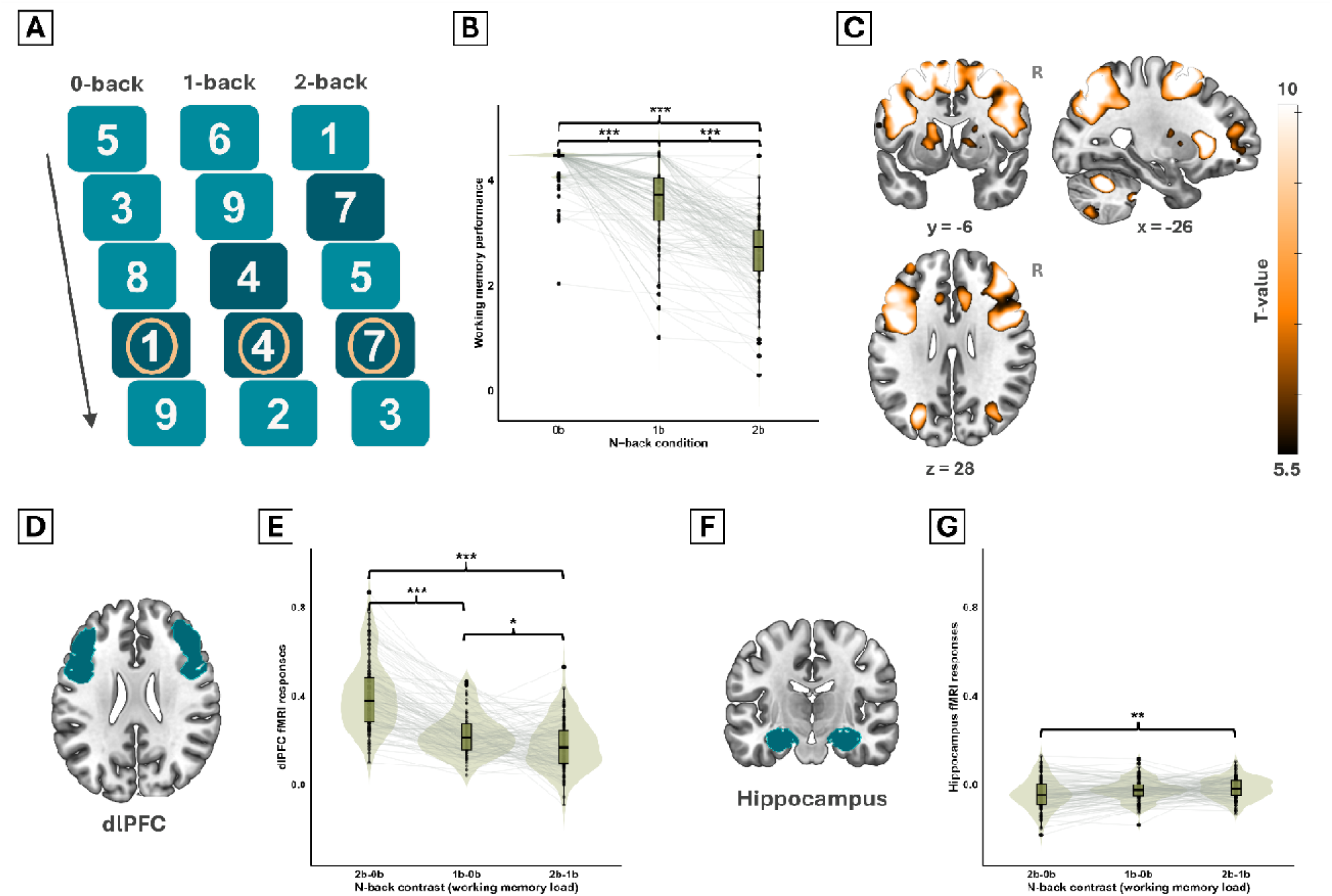
Working memory n-back task design and overall fMRI and performance results. Overview of the n-back task design, with circles indicating target stimuli per condition (**A**). Overall working memory performance dprime scores per n-back condition (**B**). Overall working memory-related fMRI brain activity, contrast in BOLD response between 2-back and 0-back condition of the n-back task thresholded at p=0.05 FWE corrected (**C**). Independent ROI map of dlPFC (**D**) and overall dlPFC fMRI responses per n-back contrast (**E**). Independent ROI map of hippocampus (**F**) and overall hippocampal fMRI responses per n-back contrast (**G**). In violin plots (**B,E,G**), individual paired samples are connected by a line and the width of the violin shapes indicates the sample density. Boxplots within violins indicate IQR and median. Significant differences (Tukey’s range test) between n-back conditions and contrasts are indicated with asterisks (‘***’ 0.001, ‘**’ 0.01, ‘*’ 0.05). BOLD = blood-oxygen level dependent; dlPFC = dorsolateral prefrontal cortex; FWE = family-wise error; IQR = inter quartile range; ROI = region of interest.

All group effects on WM-related fMRI responses and performance can be found in **Table 3**. No significant between-group differences were found for WM performance (b=0.10 (CI -0.19; 0.40), p=0.50) (**Figure 5a**). Post-intervention WM-related fMRI responses in the dlPFC were marginally significantly higher in the CDMV group compared to placebo (b=0.08 (CI 0.00; 0.16), p=0.06, ηp²=0.04), which was driven by the right dlPFC (p=0.02, ηp²=0.04) (**Figure 5b**). Post-intervention WM-related fMRI responses in the hippocampus were significantly higher in the CDMV group compared to placebo (b=0.05 (CI 0.01; 0.08), p=0.01, ηp²=0.09) (**Figure 5c**). The gPPI analysis showed that WM-related hippocampal connectivity with the dlPFC was also higher post-intervention in the CDMV group compared to placebo (b=0.04 (CI 0.00; 0.08), p=0.03, ηp²=0.07) (**Figure 5e**). Exploratory whole-brain fMRI group results can be found in **Supplementary table 13** and **Supplementary figure 15**. Overall, the CDMV group showed higher WM-related fMRI responses compared to the placebo group, but no increased WM performance.

**Table 3.**
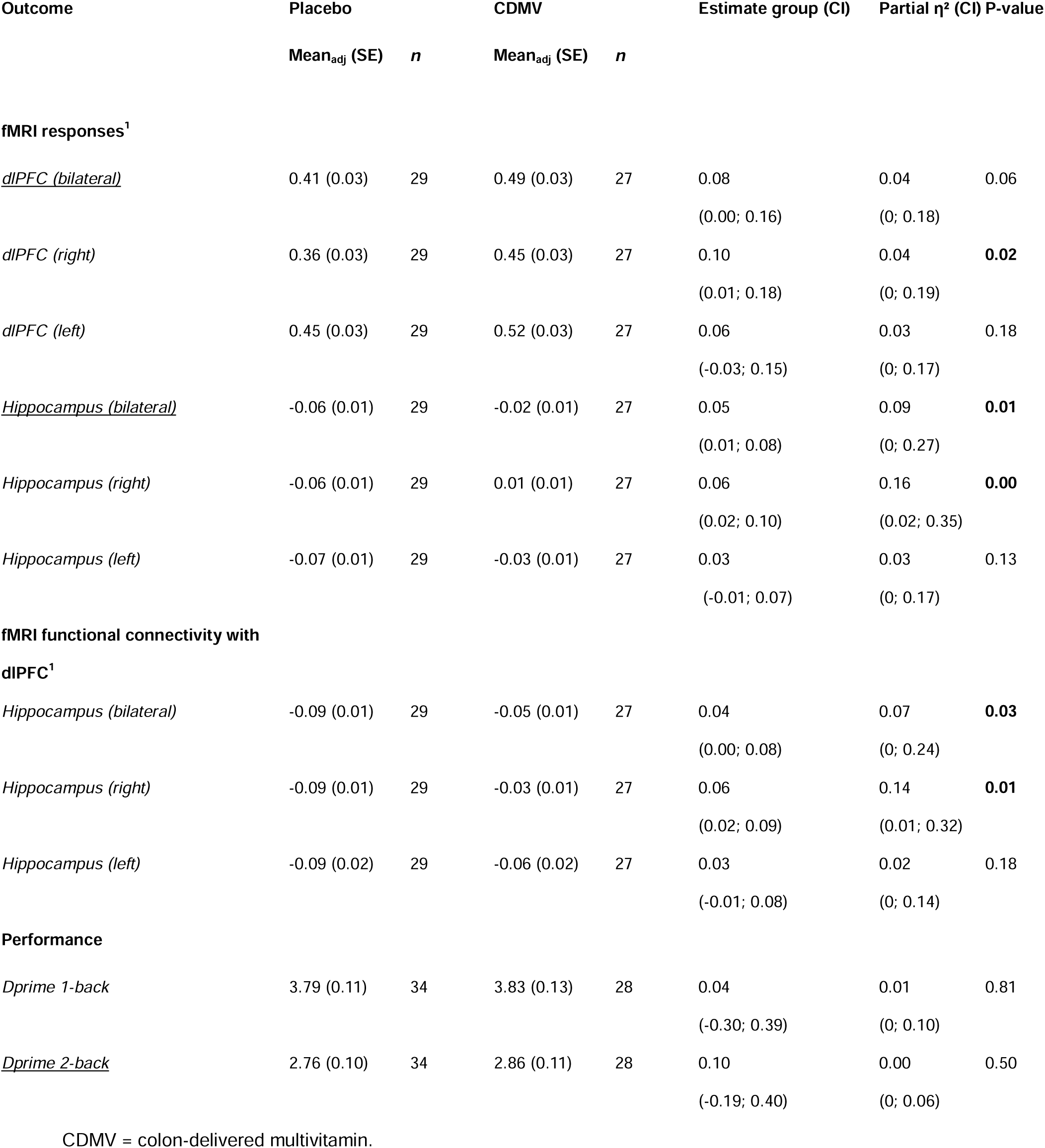
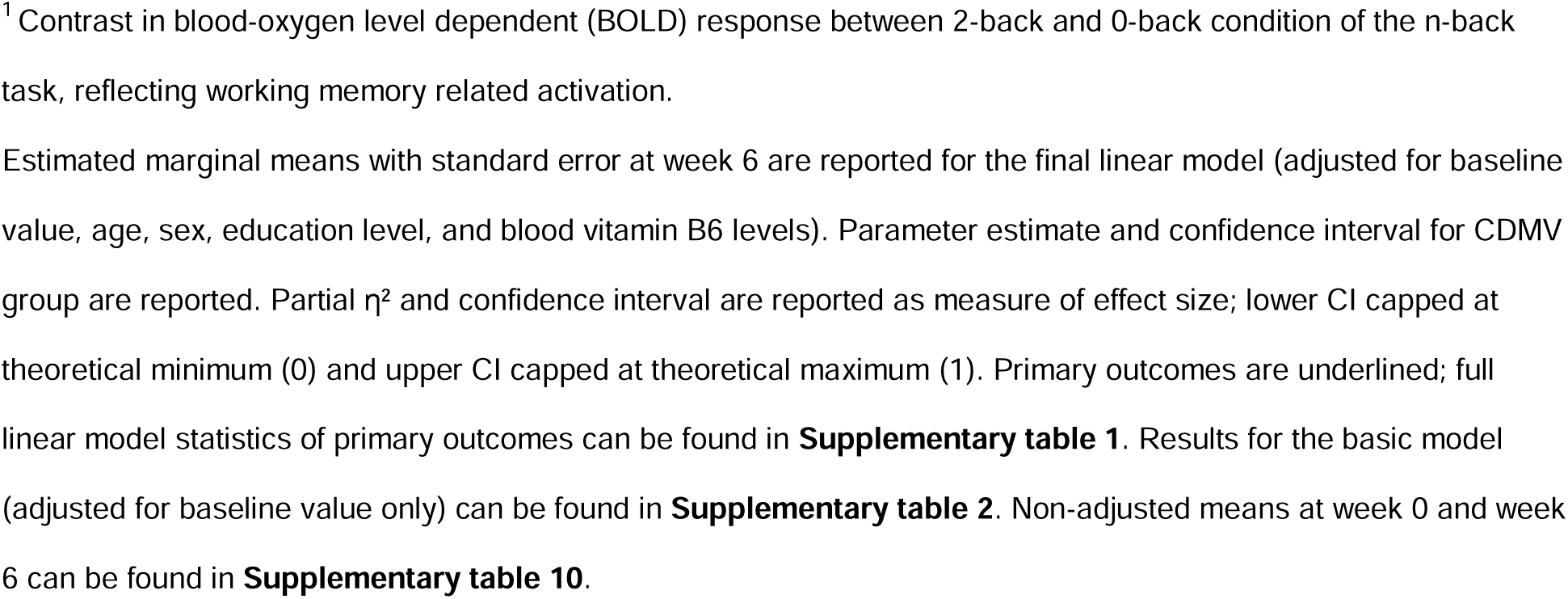
Intervention-induced effects in placebo and CDMV group at week 6 on fMRI working memory n-back task in older adults at risk of cognitive decline.

| Outcome | Placebo | | CDMV | | Estimate group (CI) | Partial $\eta^2$ (CI) | P-value |
| --- | --- | --- | --- | --- | --- | --- | --- |
|  | Mean<sub>adj</sub> (SE) | *n* | Mean<sub>adj</sub> (SE) | *n* |  |  |  |
| fMRI responses<sup>1</sup> |  |  |  |  |  |  |  |
| *dIPFC (bilateral)* | 0.41 (0.03) | 29 | 0.49 (0.03) | 27 | 0.08<br>(0.00; 0.16) | 0.04<br>(0; 0.18) | 0.06 |
| *dIPFC (right)* | 0.36 (0.03) | 29 | 0.45 (0.03) | 27 | 0.10<br>(0.01; 0.18) | 0.04<br>(0; 0.19) | **0.02** |
| *dIPFC (left)* | 0.45 (0.03) | 29 | 0.52 (0.03) | 27 | 0.06<br>(-0.03; 0.15) | 0.03<br>(0; 0.17) | 0.18 |
| *Hippocampus (bilateral)* | -0.06 (0.01) | 29 | -0.02 (0.01) | 27 | 0.05<br>(0.01; 0.08) | 0.09<br>(0; 0.27) | **0.01** |
| *Hippocampus (right)* | -0.06 (0.01) | 29 | 0.01 (0.01) | 27 | 0.06<br>(0.02; 0.10) | 0.16<br>(0.02; 0.35) | **0.00** |
| *Hippocampus (left)* | -0.07 (0.01) | 29 | -0.03 (0.01) | 27 | 0.03<br>(-0.01; 0.07) | 0.03<br>(0; 0.17) | 0.13 |
| fMRI functional connectivity with dIPFC<sup>1</sup> |  |  |  |  |  |  |  |
| *Hippocampus (bilateral)* | -0.09 (0.01) | 29 | -0.05 (0.01) | 27 | 0.04<br>(0.00; 0.08) | 0.07<br>(0; 0.24) | **0.03** |
| *Hippocampus (right)* | -0.09 (0.01) | 29 | -0.03 (0.01) | 27 | 0.06<br>(0.02; 0.09) | 0.14<br>(0.01; 0.32) | **0.01** |
| *Hippocampus (left)* | -0.09 (0.02) | 29 | -0.06 (0.02) | 27 | 0.03<br>(-0.01; 0.08) | 0.02<br>(0; 0.14) | 0.18 |
| Performance |  |  |  |  |  |  |  |
| *Dprime 1-back* | 3.79 (0.11) | 34 | 3.83 (0.13) | 28 | 0.04<br>(-0.30; 0.39) | 0.01<br>(0; 0.10) | 0.81 |
| *Dprime 2-back* | 2.76 (0.10) | 34 | 2.86 (0.11) | 28 | 0.10<br>(-0.19; 0.40) | 0.00<br>(0; 0.06) | 0.50 |

CDMV = colon-delivered multivitamin.
<sup>1</sup> Contrast in blood-oxygen level dependent (BOLD) response between 2-back and 0-back condition of the n-back task, reflecting working memory related activation.
Estimated marginal means with standard error at week 6 are reported for the final linear model (adjusted for baseline value, age, sex, education level, and blood vitamin B6 levels). Parameter estimate and confidence interval for CDMV group are reported. Partial $\eta^2$ and confidence interval are reported as measure of effect size; lower CI capped at theoretical minimum (0) and upper CI capped at theoretical maximum (1). Primary outcomes are underlined; full linear model statistics of primary outcomes can be found in **Supplementary table 1**. Results for the basic model (adjusted for baseline value only) can be found in **Supplementary table 2**. Non-adjusted means at week 0 and week 6 can be found in **Supplementary table 10**.

**Figure 5.**
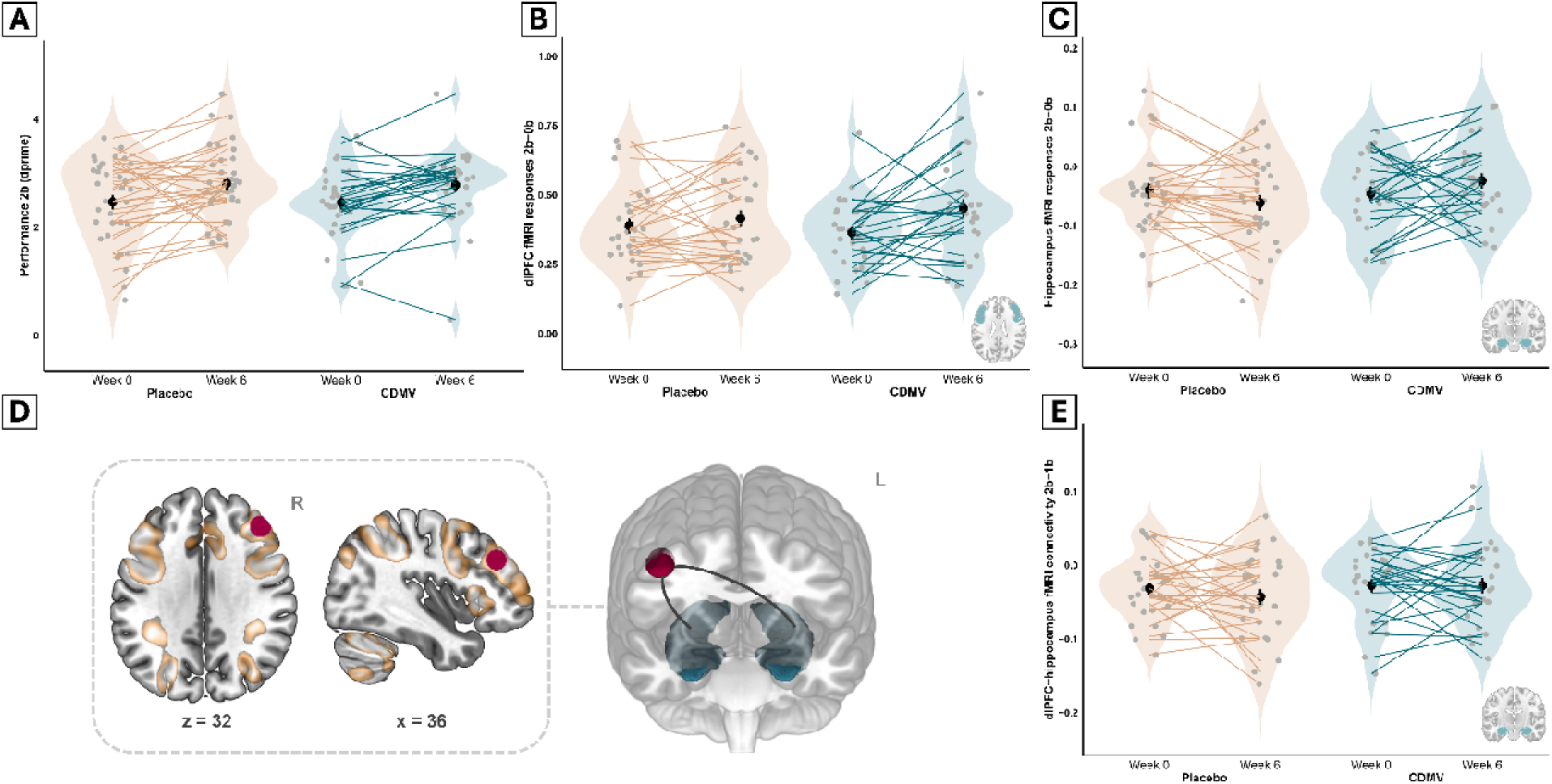
Working memory n-back task fMRI responses and performance at baseline and after the 6-weeks intervention for placebo and CDMV group. Working memory performance (2-back dprime) at baseline and after the 6-weeks intervention for placebo (*n*=34) and CDMV group (*n*=28) (**A**). Working memory dlPFC (**B**) and hippocampus (**C**) fMRI responses (2b-0b) at baseline and after the 6-weeks intervention for placebo (*n*=29) and CDMV group (*n*=27). Selected gPPI seed region in the dlPFC (red) on 2D map with overall working memory related fMRI brain activity (orange, high opacity) and 3D map with independent hippocampus ROI (blue) (**D**). Working memory task-related connectivity of hippocampus with seed region dlPFC at baseline and after the 6-weeks intervention for placebo (*n*=29) and CDMV group (*n*=27) (**E**). In violin plots (**A-C, E**), individual paired samples are connected by a line and the width of the violin shapes indicates the sample density. Circled shapes with lines within the violin indicate the group mean and SE. CDMV = colon-delivered multivitamin; dlPFC = dorsolateral prefrontal cortex; gPPI = generalized psychophysiological interactions; ROI = region of interest.

#### Other neurocognitive outcomes

No significant group effects were found on other neurocognitive secondary outcomes, including cerebral perfusion, intracranial neuroinflammatory metabolites (myo-inositol, total choline, and total creatine) and neuropsychological tests (see **Supplementary table 8** and **Supplementary figures 10, 11 and 12**). No significant group effects were found on perceived stress, anxiety, and depression scores (**Supplementary table 9**).

### Gut-brain correlations of within-subject changes over time

To explore gut-brain correlations of within-subject changes over time, we employed Spearman correlation analyses on the change values (Δ post-pre) of our primary outcomes across both intervention groups. Corrected for baseline values, changes (Δ post-pre) in total fecal SCFA correlated positively with changes in both WM-related dlPFC fMRI responses (Spearman’s Rho = 0.31, p=0.02) and WM performance (Spearman’s Rho = 0.43, p=0.001) (**Figure 6**). These correlations were driven by acetic acid, propionic acid, butyric acid individually (see **Supplementary Figure 16** for a gut-brain heatmap). For both changes in WM-related dlPFC fMRI responses (Fisher’s Z = 0.38, p=0.71) and WM performance (Fisher’s Z = 1.39, p = 0.17), the correlation with changes in total fecal SCFAs did not differ significantly between the multivitamin and placebo groups.

**Figure 6.**
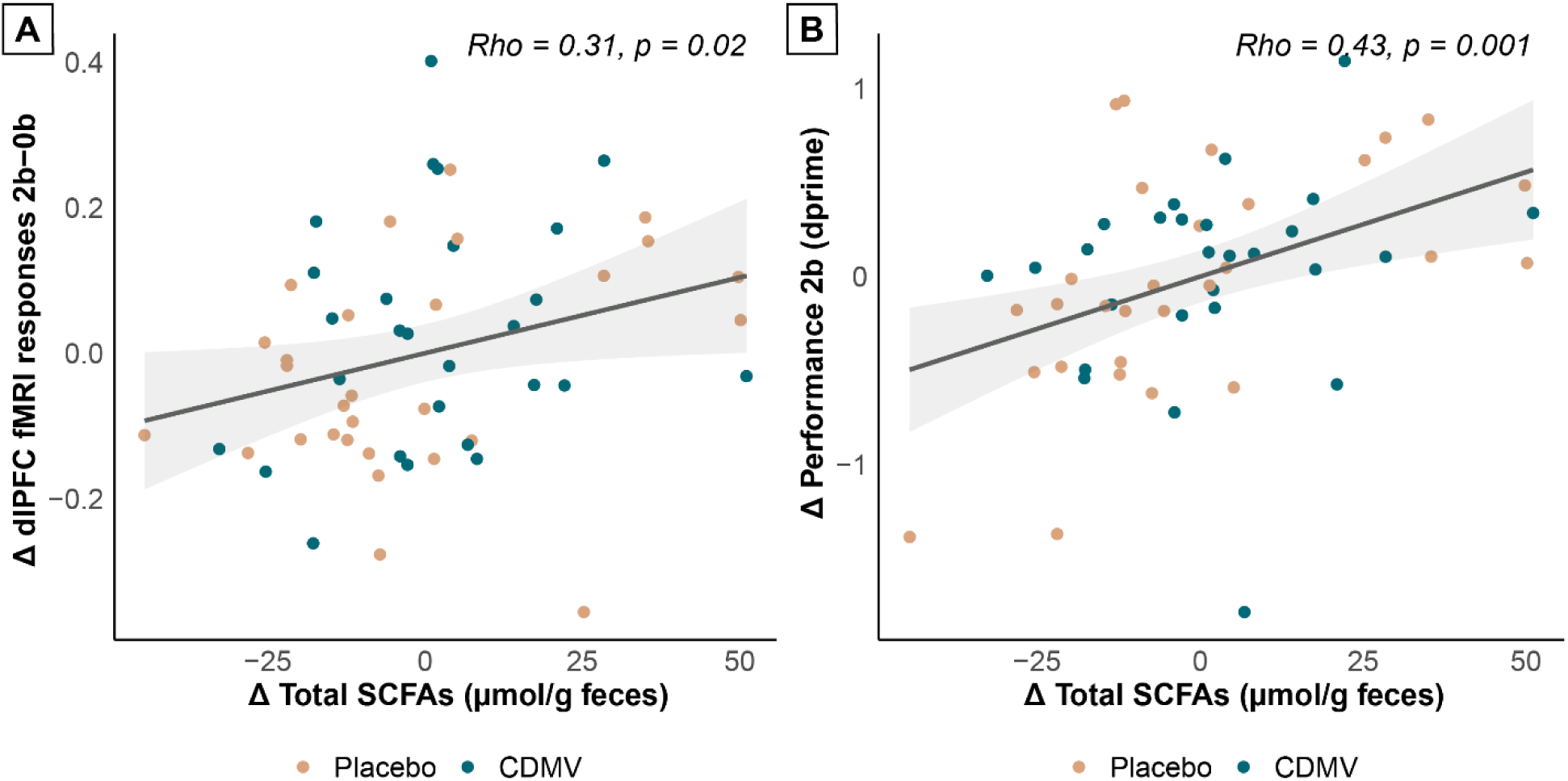
Scatterplots delta SCFAs and delta working memory for both intervention groups combined. Scatterplots representing correlation between delta score (post-value minus pre-value) of total SCFAs and working memory-related fMRI responses in dlPFC (fMRI n-back task 2-back minus 0-back contrast) (**A**) and total SCFAs and working memory performance (n-back task 2-back condition) (**B**). Dots represent individual participants. Orange color represents the placebo group, blue color represents the CDMV group. Trendline and confidence intervals are displayed in black and grey. Spearman’s rho and p-value are corrected for baseline value and reported top right. CDMV = colon-delivered multivitamin; dlPFC = dorsolateral prefrontal cortex; SCFAs = short-chain fatty acids.

Additionally, we explored gut-brain correlations over time between primary WM outcomes and secondary gut outcomes showing significant group effects: *Lachnospiraceae FCS020 group*, *Eubacterium ventriosum group*, *Ruminococcus gauvreauii group* and the fecal MCFA hexanoic acid. No significant gut-brain correlations were found for these secondary gut outcomes (see **Supplementary Figure 17**). Moreover, Fisher r-to-z tests did not show differential correlations per intervention group.

## Discussion

The goal of this study was to investigate effects of a 6-week CDMV supplementation versus placebo on the gut-brain axis in older adults, with fecal SCFAs, WM-related fMRI responses, and WM performance as primary outcomes. In addition, we examined multiple secondary gut-, immune-, and brain-outcomes that are hypothesized to play a role in gut-brain communication in aging. Our results demonstrate that CDMV supplementation enhances WM-related recruitment of the (right) dlPFC and hippocampus. However, we observed no effects on fecal SCFA levels or WM performance. Among our secondary outcomes, we only found significant increases of fecal levels of three microbial genera and the MCFA hexanoic acid within the multivitamin group. Finally, we also assessed gut-brain correlations over time across both intervention groups, and found a positive correlation between change in SCFAs, WM-related dlPFC fMRI responses and WM performance.

### Effects on fecal SCFAs

We hypothesized that CDMV supplementation would increase fecal SCFA levels compared to placebo, but we found no significant between-group differences. Previous *in vitro* (33, 34) and clinical (34, 35) work showed that individual colon-targeted vitamins (specifically vitamin B2 and C) could increase fecal SCFA levels. Given that baseline fecal SCFA levels of our participants were comparable with levels in other clinical studies (34, 35), a ceiling effect is unlikely to account for our deviating results. Instead, various factors could explain why we could not replicate these results in our population of older adults. To start with, we were the first to investigate this specific combination of colon-delivered vitamins (B2, B3, B6, B9, C, D) at dosages selected based on prior evidence, safety, and technical feasibility. In the previous mentioned pilot study of Pham et al. (34), individual vitamins (A, B2, C, D, E), some overlapping with our formulation, were tested at higher doses (ranging from 2.5- to 7.5-fold), identifying B2 and C as most promising. However, they also demonstrated that a combination of vitamin B2 and C did not enhance SCFA levels compared to placebo, suggesting that potential interactions between vitamins may have affected the observed effects on SCFAs in our study. Additionally, our supplement lacked vitamin B1, which has only very recently been identified as a rate-limiting dietary component for SCFA production (83). Secondly, Pham et al. (34) used healthy young volunteers, whereas we examined CDMV supplementation in older adults. Age-related changes of the gastro-intestinal tract in our older population - including altered pH levels, transit times, and dysbiosis (84) - could have modified both colon-delivery kinetics and microbial responsiveness, even though we excluded proton-pump inhibitor and antibiotic users. Moreover, the six-week duration may have been insufficient for this population, even though a four-week intervention led to increases in SCFAs by vitamin C in healthy young volunteers (34). Finally, although habitual dietary fiber consumption was included as a covariate, it may still have constrained the vitamin’s co-factor effects on SCFA fermentation, given that SCFA production requires fermentable fiber as a substrate. While total fiber intake was comparable across our study (20 g/day) and the study of Pham et al. (22 g/day) (34), our dietary assessment did not distinguish between fermentable and non-fermentable fiber types, and such potential differences may have better explained SCFA production capacity.

Of crucial importance, fecal SCFA concentrations represent a proxy measure for colonic SCFA production rather than a direct assessment, as most SCFAs are produced in the proximal colon and rapidly absorbed (85). Thus, the absence of effects on fecal SCFAs does not preclude the possibility of altered colonic SCFA production by CDMV supplementation. Although measuring circulating SCFAs could theoretically provide insight into more proximal colonic production, blood SCFA levels are influenced by intestinal absorption, portal vein transport, and hepatic first-pass metabolism (86), rendering them an indirect measure as well. Previous research demonstrated that colon-delivered SCFAs increased blood concentrations without affecting fecal levels (87, 88), illustrating the dissociation between these measurements. However, both fecal and circulating SCFA concentrations showed no significant between-group effects in our study. Nevertheless, while the absence of CDMV-induced changes in both markers reduces the likelihood of substantial SCFA effects, it does not entirely preclude subtle alterations in colonic fermentation, given that both measures are indirect.

### Effects on working memory

We were the first to assess neurocognitive effects of CDMV supplementation, by examining WM-related fMRI responses in the dlPFC and hippocampus, and WM performance. We showed that CDMV supplementation led to higher WM-related recruitment of both the (right) dlPFC and hippocampus compared to placebo, as measured by a numerical n-back task. We observed a trend towards increased dlPFC responses (small-to-medium effect size), which was predominantly driven by the right dlPFC. This finding can be interpreted in light of two competing theoretical frameworks. Aged individuals typically exhibit reduced WM-related fMRI responses in the dlPFC compared to younger adults (72), which further declines in mild cognitive impairment (89), suggesting that the increased fMRI responses observed in our study are beneficial. However, alternative theories propose that aging can lead to neural over-recruitment as a compensatory mechanism to maintain WM performance (90), complicating the interpretation of increased activation patterns. Given the absence of intervention effects on behavioral WM performance, both theoretical perspectives remain plausible. Nevertheless, our findings demonstrate that a relatively brief 6-week intervention with CDMV supplements can modulate neural correlates of WM. It is conceivable that the neural reorganization we observed requires additional time to translate into detectable behavioral improvements, and that a longer intervention period, such as 12 weeks instead of six (22), may be needed for the observed changes in fMRI responses to manifest as measurable gains in WM performance. This interpretation is further supported by our correlational findings showing that within-subject increases in SCFA concentrations over time positively correlate with increases in both dlPFC responses and WM performance, suggesting that enhanced fMRI responses may indeed be beneficial rather than compensatory in our study population. While no previous studies have examined effects of colon-targeted interventions on WM-related neuroimaging in older adults, limited evidence supporting such an effect exists in younger populations (39, 43). Papalini et al. (43) demonstrated that a 4-week probiotic intervention in young adults improved WM performance after acute stress, which correlated with right PFC recruitment during cognitive control. These findings align with our results, particularly regarding the right-driven PFC effects.

In addition to effects on dlPFC recruitment, we observed a significant effect on WM-related hippocampal responses (medium-to-large effect size). Although the hippocampus showed minimal overall task activation – consistent with its less pronounced supportive role in WM (47) – post-intervention responses were significantly higher in the multivitamin group compared to the placebo group. Using task-based functional connectivity analyses, we demonstrated that higher hippocampal responses in the multivitamin group were associated with dlPFC connectivity, suggesting modulation of the dlPFC-hippocampal circuit during WM (91). The stronger hippocampal effect compared to dlPFC may reflect its smaller, more confined ROI, potentially increasing statistical sensitivity. Given the hippocampus’s vulnerability to aging and neuroinflammation and its central role in cognitive decline (42), these findings are of clinical relevance.

Notably, post-intervention blood vitamin B6 levels were significantly elevated in the multivitamin group, while all other vitamins remained unchanged. Given that only vitamin B6 showed systemic elevation, this strongly suggests colonic absorption of this specific vitamin (92) rather than general failure of the colon-delivery coating resulting in full small intestine absorption of all vitamins. As vitamin B6 can directly influence brain health through mechanisms beyond the gut-brain axis (93), we included post-intervention blood vitamin B6 as a covariate in the neurocognitive analyses. Vitamin B6 was not a significant predictor in these models, suggesting minimal confounding of our primary outcomes. Nevertheless, future studies employing colon-targeted vitamin formulations should account for potential colonic absorption of vitamin B6.

### Effects on secondary peripheral outcomes

Given the neurocognitive effects of CDMV supplementation without concomitant changes in fecal SCFAs, the question remains which peripheral-central pathways mediated these central effects. To explore alternative peripheral mechanisms, we assessed a broad array of secondary outcomes, including gut integrity markers, blood SCFA concentrations, and immune markers. However, the majority of these measures likewise showed no significant between-group differences, despite our liberal approach to multiple testing corrections. Specifically, we applied corrections only within high-dimensional datasets (microbiome composition, whole-brain fMRI), but not for individual markers within other secondary outcome domains or across the different outcome categories. Given the exploratory, hypothesis-generating nature of this mechanistic study, this approach was chosen to maintain sensitivity for detecting potential biological mechanisms underlying the observed effects.

Notably, we did find three bacterial genera that were significantly higher post-intervention in the multivitamin group compared to placebo: *Lachnospiraceae FCS020 group*, *Eubacterium ventriosum group*, and *Ruminococcus gauvreauii group*. Effects on *Lachnospiraceae* are of particular interest, as *Lachnospiraceae* are well-known SCFA producers (94). Moreover, a pilot study in healthy individuals found substantial increases in fecal *Lachnospiraceae* as result of vitamin C supplementation (95), aligning with our results, even though supplementation in this study was not colon-delivered. Similarly, in vitro digestion and fermentation of colon-targeted vitamin B2 also increased *Lachnospiraceae* (33). Both *Eubacterium ventriosum group* and *Ruminococcus gauvreauii group* are also involved in SCFA production, predominantly butyrate (96). Nevertheless, determining the health implications of such shifts in specific genera remains challenging, as the functional consequences can only be understood within the context of the complete bacterial ecosystem. And, importantly, within-subject changes of these bacterial genera did not correlate with changes in WM outcomes.

In addition, we also found a significant group difference in fecal hexanoic acid levels, a MCFA, with higher concentrations in the multivitamin group. However, these changes likewise did not correlate with within-subject changes in WM outcomes over time. Moreover, since MCFAs are not exclusively produced by bacterial fermentation but can also originate from the diet (97), and given that absolute concentrations were relatively low compared to other fatty acids, future studies should confirm its clinical relevance.

### Gut-brain correlations

Beyond group-level comparisons, we examined associations between changes in our primary gut and neurocognitive outcomes across both intervention groups. Importantly, such within-subject change correlations offer greater sensitivity than cross-sectional approaches and provide mechanistic insights. We observed that increases in fecal SCFAs positively correlated with increases in both WM-related dlPFC responses and WM performance, independent of intervention group. These findings support the hypothesis that fecal SCFAs are relevant predictors of neurocognitive function and emphasize the clinical potential of SCFA-targeted interventions in aging populations. However, it remains unclear to what extent CDMV supplementation contributed to the observed SCFA changes and which other (lifestyle-related) factors may have played a role. Moreover, the involved peripheral-to-central pathways remain unresolved as well, warranting further mechanistic investigation.

### Recommendations

Our study provides valuable insights into the effects of CDMV supplementation on the gut-brain axis in aging by incorporating a broad range of neurocognitive, blood, and fecal measurements. Yet, as discussed throughout this paper, several limitations and methodological considerations remain.

Altogether, the unresolved questions regarding involved gut-brain mechanisms provide important starting points for future research. To better characterize these mechanisms in human studies, we recommend that future studies incorporate additional markers and employ complementary innovative techniques to elucidate how CDMV supplements exert their neurocognitive effects. For instance, using a smart capsule (98) or naso-intestinal catheter (99) could help visualize vitamin release patterns and enable direct measurement of proximal colonic SCFA concentrations. Moreover, to comprehensively assess functional microbiome activity, other microbial metabolites beyond SCFAs, such as secondary bile acids and tryptophan metabolites could be measured in feces and blood, or untargeted metabolomic approaches could be applied (6). Finally, we propose that future research also examines blood-brain barrier integrity (100), for example using gadolinium-enhanced MRI or diffusion-weighted arterial spin labeling, and vagal nerve activity (101) as potential mediators of gut-brain communication (102).

Beyond these methodological considerations for measuring gut-brain mechanisms, and further addressing the study’s limitations, we propose several recommendations for the design and conduct of future clinical trials targeting the gut-brain axis in aging. First, the CDMV formulation requires optimization for older adults through systematic preclinical evaluation of vitamin selection, dosing, and combinations. Particular attention should be given to vitamin B1, given its potential key role in SCFA production (83). Additionally, in vitro kinetic studies accounting for age-related gastrointestinal changes (e.g., altered pH, transit times) would be valuable to optimize colonic release and microbial responsiveness in older adults. Second, longer intervention durations would enable evaluation of behavioral cognitive outcomes and sustained SCFA effects. Third, enriching CDMV supplements with prebiotic fibers or combining them with a diet high in fermentable fibers might enhance microbiome-mediated benefits, as SCFA production may have been constrained by insufficient fermentable fiber intake. Moreover, detailed monitoring of dietary fiber intake, distinguishing between fermentable, non-fermentable, and prebiotic fibers, would enable a more nuanced interpretation of the effects. While comprehensive fiber questionnaires are still under development, tools such as the FiberTAG (103) could provide more precise fiber intake assessment for analytical adjustments. Finally, directly administering colon-delivered SCFAs (87, 88) might be of clinical relevance, given our findings on the positive association between fecal SCFAs and working memory increases over time.

### Conclusions

Collectively, our results show that a 6-week CDMV supplementation affects neurocognitive functioning by increasing working memory-related fMRI responses. Although our comprehensive assessment of peripheral markers provides valuable insights, the specific peripheral mechanisms mediating the neurocognitive effects of CDMV supplements remain to be further elucidated. We recommend future studies to build upon these findings using complementary markers and additional innovative techniques to gain a deeper mechanistic understanding. Moreover, we demonstrated that, independent of intervention group, within-subject changes in fecal SCFA levels correlated positively with changes in WM outcomes. These correlations of changes over time provide human evidence for the gut-brain axis in cognitive aging beyond cross-sectional associations and emphasize the potential of SCFA-increasing interventions in older adults. Future studies should determine the role of CDMV supplementation within this gut-brain relation.

## Acknowledgements

We would like to thank all the participants of the COMBI study for volunteering. We would like to express our gratitude to Diede Booltink and Nele Ziegler for their practical contributions in MRI scanning; Paul Gaalman for assistance in the neuroimaging lab and setting up the MRI protocol; Karin Mudde for assistance in the laboratory; Marcel Zwiers, Ruben Doornbosch, and Lisa-Katrin Kaufmann for their advice on fMRI analyses; Nick Puts, Viola Hollestein and Jilly Naaijen for their advice on ^1^H-MRS analyses; Gerben de Gier and Marlies Diepeveen-de Bruin for their help with fecal SCFA analyses; Cesare Lotti for performing the blood SCFA analyses; Nhien Ly for performing the blood cytokine analyses; Els Oosterink for conducting the permeability marker blood analyses; and Cindy van der Schaaff for performing the ddPCR analyses. Special thanks to Joao Caldas Paulo for advice on statistical analyses, and Jos Boekhorst for their help with pre-processing the 16S rRNA gene sequencing data. We also acknowledge all MSc thesis students that assisted within the COMBI study for their practical contributions. Lastly, we extend our appreciation to the WUR-HNH Human Nutrition Research Unit team and the RU-DCCN research assistant pool for their support in the practical execution of the study.

## Author Contributions

LBR, MRvL, JJM, WTS, JMO, MPHvT, and EA designed research. LBR, MRvL, MMGB, NMPO, and JvO conducted research. AR and RES provided essential materials. LBR, MPHvT, MRvL, MMGB and JvO analyzed data. LBR and MPHvT performed statistical analysis. LBR, MPHvT, MRvL, JMO, and EA wrote the manuscript. LBR had primary responsibility for final content. AA and UV analyzed blood SCFAs. WTS deceased before manuscript was finalized, and has not reviewed and approved final manuscript. All other authors (LBR, MRvL, MMGB, NMPO, JvO, AA, UV, AR, RES, JJM, GJEJH, JMO, MPHvT, and EA) have reviewed the manuscript. All other authors read and approved the final manuscript.

## Ethical Statement and Consent

The COMBI study was conducted in compliance with the Declaration of Helsinki for research involving human participants and the Medical Research Involving Human Subjects Act (WMO; ‘Wet Medisch-wetenschappelijk Onderzoek met mensen’). The complete procedure was approved by the local Ethics Committee (METC Oost-Nederland, NL80063.091.22) on August 22^nd^ 2022 and registered in the Clinical Trial Register (ClinicalTrials.gov ID: NCT05675007) on October 24^th^ 2022. Written informed consent was obtained from each participant during the first study visit.

## Data availability

Data available upon request. Data will be made publicly and freely available without restriction at Radboud Data Repository after publication.

## Abbreviations

^1^H-MRS: Proton magnetic resonance spectroscopy
AE: Adverse event
BCFA: Branched-chain fatty acid
BOLD: Blood-oxygen level dependent
BSS: Bristol Stool Scale
CBF: Cerebral blood flow
CHESS: Chemically selective water suppression
CRP: C-reactive protein
CVD: Cardiovascular disease
dlPFC: Dorsolateral prefrontal cortex
DST: Digit Span Test
DSST: Digit Symbol Substitution Test
EPI: Echo planar imaging
FDR: False discovery rate
FWE: Family-wise error
FFQ: Food Frequency Questionnaire
FWHM: Full width at half maximum
GC-FID: Gas chromatography with flame ionization detection
GI: Gastrointestinal
GLM: Generalized linear model
(g)PPI: (generalized) Psychophysiological interaction
GSRS: Gastrointestinal Symptom Rating Scale
HADS: Hospital Anxiety and Depression Scale
hs-CRP: High sensitivity C-reactive protein
ICA-AROMA: Independent Component Analysis-based Automatic Removal Of Motion Artifacts
IFN: Interferon
IL: Interleukin
IQR: Interquartile range
LBP: Lipopolysaccharide Binding Protein
LCN-2: Lipocalin-2
MCFA: Medium-chain fatty acid
MP2RAGE: Magnetization-prepared 2 rapid gradient echo
(pC)ASL: (pseudo-continuous) Arterial spin labeling
PRESS: Point RESolved Spectroscopy
PSS: Perceived Stress Scale
RAVLT: Rey Auditory Verbal Learning Test
ROI: Region-of-interest
RU-DCCN: Radboud University, Donders Centre for Cognitive Neuroimaging
SAE: Serious adverse event
SCFA: Short-chain fatty acid
sIgA: Secretory Immunoglobulin A SNR Signal-to-noise ratio
SPM12: Statistical Parametric Mapping, version 12 TE Echo time
TICS-M1: Telefonisch Interview voor Cognitieve Status(TICS-M) version 1
TMT: Trail Making Test
TNF: Tumor necrosis factor
TR: Repetition time
VFT: Verbal Fluency Test
WBC: White blood cell
WM: Working memory
WUR-HNH: Wageningen University & Research, Division of Human Nutrition and Health

