## Supplement for "Colon-delivered multivitamin supplementation enhances working memory-related fMRI responses in older adults: a randomized, placebo-controlled trial"

**Supplementary Methods** **2**

Other fecal laboratory analyses 2

Other blood laboratory analyses 3

Other neuroimaging analyses 6

Neuropsychological test battery analysis 9

Preprocessing with fMRIPrep 10

**Supplementary Data** **15**

Multiple linear regression models (primary outcomes) 15

Basic model (reported outcomes) 16

Habitual dietary intake and blood vitamin concentrations 18

User experiences and adverse effects 23

Secondary outcomes 26

Non-adjusted group means 45

Microbiome analysis 51

Whole-brain fMRI analysis 69

Gut-brain correlations over time 74

**References** **78**

### Supplementary Methods

#### Other fecal laboratory analyses

##### Fecal water content

For fecal water content measurement, approximately 1 gram of frozen, mill-homogenized fecal sample was weighed, and freeze-dried using a freeze-dryer (FD8518, Ilshin Biobase, Korea) according to manufacturer's instructions. The empty tube weight was also determined for calculation purposes. After 48 hours of freeze-drying, the samples were retrieved and weighed. The water content percentage was calculated using the formula: 100-[(dry weight (g)/wet weight (g))*100].

##### Fecal pH and redox potential

For the measurement of fecal pH and redox potential, 500 mg of a frozen, mill-homogenized fecal sample was diluted in 5 mL of MQ water. The samples were homogenized for 15 min on a horizontal shaker at a speed of 300/min (Universal Shaker SM-30, Edmund Bühler GmbH, Bodelshausen, Germany) and centrifuged (5292 g, 10 min, at 21°C). Fecal pH and redox potential were measured using a ConeFET pH/EC/ORP/T probe (Sentron Europe B.V., Leek, The Netherlands) according to the manufacturer's instructions. The device was calibrated using standard pH solutions covering the range of pH 2–12.

##### Fecal inflammatory markers

Fecal inflammatory markers calprotectin, lipocalin-2 (LCN-2), and secretory immunoglobulin A (sIgA) were measured in hammered-only fecal samples. Calprotectin and sIgA were quantified using the IDK® Calprotectin ELISA kit and IDK® sIgA ELISA kit, respectively (#K6927 and #K8870, Immundiagnostik AG, Bensheim, Germany), according to the manufacturer's protocols. LCN-2 was measured with the Human LCN-2/NGAL ELISA kit (RD191102200R, BioVendor, Brno, Czech Republic), following the manufacturer's instructions. Before the ELISA assays were performed, fecal extractions took place. For fecal extraction, 60 mg of hammered-only feces was weighed into a 15 mL Falcon tube. For calprotectin and sIgA, the samples were diluted 1:100 in extraction buffer (IDK). For LCN-2, the samples were diluted 1:50 using extraction buffer (C005821, BioVendor). The tubes were vortexed for 30 sec, placed on a horizontal shaker at 300 rpm for 30 min, and subsequently centrifuged at 3000g for 10 min at RT. The ELISA plates were read using a VANTAstar® Microplate Reader (BMG Labtech, Ortenberg, Germany) at absorbance wavelengths of 450 nm and 690 nm. Optical density values were converted to concentrations using GraphPad Prism software (version 8.0.1).

#### Other blood laboratory analyses

##### Blood fatty acids

Plasma SCFAs (acetic acid, propionic acid, butyric acid, valeric acid), MCFAs (heptanoic acid, hexanoic acid, octanoic acid, decanoic acid, dodecanoic acid) and BCFAs (isobutyric acid, isovaleric acid, 2-methyl butyric acid) were quantified using gas chromatography-mass spectrometry (GC-MS) according to an optimized protocol adapted from Lotti et al. (1). Briefly, 100 µL of plasma was added to a 1.5 mL Eppendorf tube containing 10 µL of

15% $H_3PO_4$, 10 µL of internal standard mixture (dissolved in methyl tert-butyl ether; MTBE), and 140 µL of MTBE. Analytes were extracted for 5 minutes using an orbital shaker, followed by centrifugation at 15,000 rpm at 5°C for 5 minutes. The supernatant was transferred into a 250 µL insert placed in a labeled 2 mL vial and sealed. Samples were stored at -20°C for a maximum of 3-4 weeks prior to analysis to prevent solvent evaporation.

GC-MS analysis was performed using an upgraded version of the previously published method (1) with a total run time of 6.5 minutes in multiple reaction monitoring (MRM) mode. Data processing was conducted using Agilent quantitative analysis software.

**Blood intestinal permeability markers**

Biomarkers indicating bacterial translocation and/or intestinal permeability were measured. ELISAs were used to quantify zonulin and LPS-binding protein (LBP) in 25 µL of serum each. Zonulin was measured using the K5601 assay kit (Immundiagnostik AG, Bensheim, Germany), and LBP was measured using the HK315-02 assay kit (Hycult Biotech, Uden, Netherlands), following manufacturers' instructions. All samples were analyzed in duplicate.

**Blood systemic inflammation markers**

The high-sensitivity C-reactive protein (hs-CRP) levels and white blood cell (WBC) counts were determined from blood droplets collected via finger prick. For hs-CRP concentration measurement, 20 µL of blood was collected in a capillary tube, directly transferred into a cuvette with buffer solution, and analyzed using the QuikRead Go analyzer (Orion Diagnostics, Ghodbunder, India) according to the manufacturer’s instructions. The measurement range was between 0.5–200 mg/L. For total white blood cell counts and differential counts of neutrophils, lymphocytes, monocytes, eosinophils, and basophils, 10 μL of blood was collected in a microcuvette, and analyzed using the HemoCue® WBC DIFF (HemoCue AB, Ängelhom, Sweden) following manufacturer's instructions. The measurement range was between 0.3-30.0×$10^9$/L. Plasma IFN-γ (range 0.33-1350 pg/mL), IL-6 (range 0.2-809 pg/mL), IL-8 (range 0.14-591 pg/mL), IL-10 (range 0.09-375 pg/mL), TNF-α (range 0.09-360 pg/mL) were quantified using a mesoscale discovery V-PLEX Pro-inflammatory Panel 1 (human) kit (K151A9H-1, Mesoscale Discovery, Gaithersburg, MD) according to the manufacturer's instructions.

#### Other neuroimaging analyses

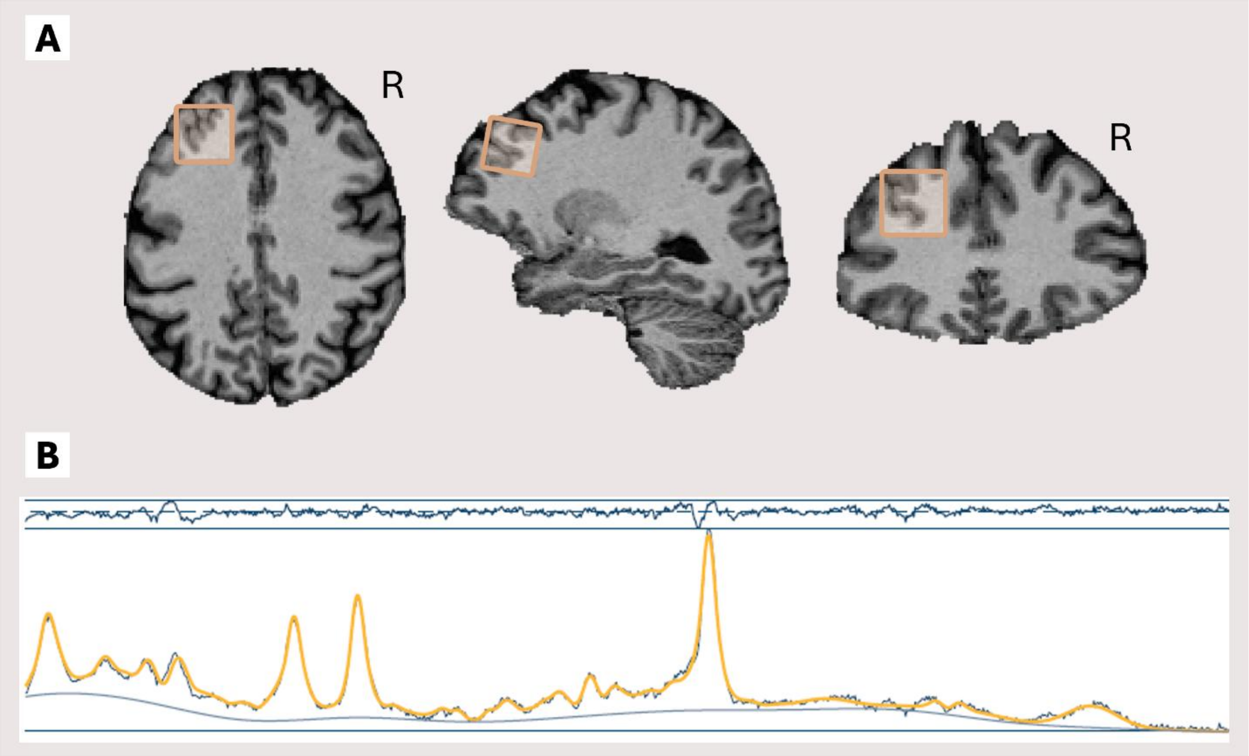


**Supplementary Figure M1. $^{1}$H-MRS example voxel and spectrum.** Transversal, sagittal and coronal image of voxel location in the left dlPFC retrieved from example participant (**A**). Fitted spectrum of example participant (**B**). The oscillating dark blue line on top represents the residuals. The yellow line represents the fitted spectrum (i.e. the modelled spectrum that is used for quantification of metabolites) and the blue line represents the observed spectrum (aligned and averaged). The blue line below the observed and fitted spectra represents a regressor for chemical shift drift.

##### Neuroinflammation ($^{1}$H-MRS)

To assess neuroinflammation, we measured intracranial levels of three neuroinflammation-associated metabolites using $^{1}$H-MRS: myo-inositol, total choline, and total creatine (3). Initially $^{1}$H-MRS was measured in both our ROIs: dlPFC and hippocampus. However, since general spectral quality of hippocampal data was insufficient, we only used dlPFC spectral data for the quantification of neuroinflammation metabolites. $^{1}$H-MRS data processing was conducted with Osprey 2.4.0 (4), an open-source software implemented in MATLAB R2024a (Mathworks Inc.; https://nl.mathworks.com/products/matlab.html) that provides automated and standardized preprocessing, linear-combination modeling, tissue correction, and quantification. Metabolite, water-unsuppressed reference, and anatomical MP2RAGE data were preprocessed, generating averaged spectra. Eddy-current correction (5) using water-unsuppressed reference data were applied, if necessary. The processed spectra were fitted using Osprey's linear combination model with default settings. Spectra were fitted across a frequency range of 0.5 to 4.0 ppm with a knot spacing of 0.4 ppm. Water data was fitted in the range of 2.0 to 7.4 ppm. Subsequently, spectroscopy data was coregistered to the participant's T1-weighted anatomical scan, and tissue segmentation was performed on the anatomical data using SPM12 to determine fractional tissue volumes within each region of interest. Tissue (relaxation time) corrected and water-scaled metabolite measures were calculated, resulting in molar concentration estimates of myo-inositol, total choline, and total creatine. Pre-aligned, post-aligned, averaged, and fitted spectra were manually inspected to evaluate spectral quality. Quality metrics included: adequate water suppression (3-4 ppm) and shimming, no lipid contamination (1-2 ppm) or chemical shift drift (indicating motion), and detection of creatine (3.03 ppm), choline (3.20 ppm), and myo-inositol (3.5 and 4.06 ppm) peaks. Besides, Osprey's reported data quality measures including creatin signal-to-noise (SNR) ratio and full width at half maximum (FWHM) values were consulted. In case of insufficient spectral quality, datasets were excluded from analysis. This led to exclusion of *n*=10 datasets. In addition, for *n*=6 participants $^1$H-MRS data was lacking for one of the sessions due to scanner issues. In the end, *n*=51 participants were included for analysis, see **Supplementary Table M1** for a summary of data quality measures and tissue fractions in our dataset. Spectral quality measure RelResA was used as covariate in models with $^1$H-MRS outcomes, to correct for small differences in data quality.

**Supplementary Table M1. Data quality measures and tissue fractions of $^1$H-MRS data (*n*=51)**

| | Cr SNR | Cr FWHM | Water FWHM | relResA | GM % | WM % | CSF % |
|---|---|---|---|---|---|---|---|
| ***Mean ± SD*** | 67.45 ± 11.79 | 6.90 ± 1.38 | 8.04 ± 1.37 | 2.86 ± 2.07 | 30 ± 7 | 67 ± 9 | 4 ± 3 |

*Cr = Creatin; SNR = signal-to-noise ratio; FWHM = full width at half maximum; GM = grey matter; WM = white matter; CSF = cerebrospinal fluid; SD = standard deviation*

##### Cerebral blood flow (ASL)

ASL data were analyzed using the FSL BASIL asl_gui toolset (6). We processed the acquired perfusion maps and calibration images to create quantified CBF images. Perfusion images were Hadamard-decoded, motion corrected, and filtered to only keep voxels with ASL signal. We analyzed the 14 volumes (7 PLDs x 2 repeats) of each pre-substracted perfusion image grouped by repeats, using a constant sub-bolus duration of 0.4 seconds (0.5-2.9 seconds) and 3D GRASE readout. Quantified CBF images were coregistered to the participant's T1-weighted anatomical scan using the FSL_ANAT pipeline, and calibrated using the $M_0$ image as a proton-density calibration image (sequence TR = 1000, calibration gain = 10). We extracted CBF values of whole-brain grey matter, white matter, and our ROIs: the dlPFC and hippocampus. Native space bilateral dlPFC masks were created for each participant, using *antsApplyTransforms* on the independent MNI dlPFC mask (used for fMRI analysis) with fMRIprep generated MNI to native space transform parameters (*antsRegistration*). Native space bilateral hippocampus masks were created using Freesurfer ASEG structural parcellation. Quantified CBF maps of all participants were visually inspected to examine data quality, and checked for acquisition errors and excessive motion artifacts. Based on these assessments, *n*=1 participant was excluded. In addition, for *n*=12 participants ASL data was lacking for one of the sessions due to scanner issues. In the end, *n*=54 participants were included for analysis.

#### Neuropsychological test battery analysis

For the neuropsychological test battery analysis, we used the following outcomes from individual tests: DST total score (forward+backward), DSST total score, RAVLT delayed recall score, TMT-B/TMT-A ratio score, and VFT total score. A composite cognition Z-score was created by combining Z-scores of the individual cognitive tests. Z-scores were reversed when necessary, meaning that higher values indicate a better cognitive performance. For *n*=1 participant, neuropsychological tests were missing for the post-intervention (T1) visit.

#### Preprocessing with fMRIPrep

Results included in this manuscript come from preprocessing performed using *fMRIPrep* 23.2.0 (Esteban et al. (2019); Esteban et al. (2018); RRID:SCR_016216), which is based on *Nipype* 1.8.6 (K. Gorgolewski et al. (2011); K. J. Gorgolewski et al. (2018); RRID:SCR_002502).

##### Preprocessing of $B_0$ inhomogeneity mappings

A total of 2 fieldmaps were found available within the input BIDS structure for this particular subject. A $B_0$ nonuniformity map (or *fieldmap*) was estimated from the phase-drift map(s) measure with two consecutive GRE (gradient-recalled echo) acquisitions. The corresponding phase-map(s) were phase-unwrapped with prelude (FSL None).

##### Anatomical data preprocessing

A total of 2 T1-weighted (T1w) images were found within the input BIDS dataset. Each T1w image was corrected for intensity non-uniformity (INU) with N4BiasFieldCorrection (Tustison et al. 2010), distributed with ANTs 2.5.0 (Avants et al. 2008, RRID:SCR_004757). The T1w-reference was then skull-stripped with a *Nipype* implementation of the antsBrainExtraction.sh workflow (from ANTs), using OASIS30ANTs as target template. Brain tissue segmentation of cerebrospinal fluid (CSF), white-matter (WM) and gray-matter (GM) was performed on the brain-extracted T1w using fast (FSL (version unknown), RRID:SCR_002823, Zhang, Brady, and Smith 2001). An anatomical T1w-reference map was computed after registration of 2 T1w images (after INU-correction) using mri_robust_template (FreeSurfer 7.3.2, Reuter, Rosas, and Fischl 2010). Brain surfaces were reconstructed using recon-all (FreeSurfer 7.3.2, RRID:SCR_001847, Dale, Fischl, and Sereno 1999), and the brain mask estimated previously was refined with a custom variation of the method to reconcile ANTs-derived and FreeSurfer-derived segmentations of the cortical gray-matter of Mindboggle (RRID:SCR_002438, Klein et al. 2017). Volume-based spatial normalization to two standard spaces (MNI152NLin6Asym, MNI152NLin2009cAsym) was performed through nonlinear registration with antsRegistration (ANTs 2.5.0), using brain-extracted versions of both T1w reference and the T1w template. The following templates were were selected for spatial normalization and accessed with *TemplateFlow* (23.1.0, Ciric et al. 2022): *FSL's MNI ICBM 152 non-linear 6th Generation Asymmetric Average Brain Stereotaxic Registration Model* [Evans et al. (2012), RRID:SCR_002823; TemplateFlow ID: MNI152NLin6Asym], *ICBM 152 Nonlinear Asymmetrical template version 2009c* [Fonov et al. (2009), RRID:SCR_008796; TemplateFlow ID: MNI152NLin2009cAsym].

**Functional data preprocessing**

For each of the 2 BOLD runs found per subject (across all tasks and sessions), the following preprocessing was performed. First, a reference volume was generated, using a custom methodology of *fMRIPrep*, for use in head motion correction. Head-motion parameters with respect to the BOLD reference (transformation matrices, and six corresponding rotation and translation parameters) are estimated before any spatiotemporal filtering using mcflirt (FSL , Jenkinson et al. 2002). The estimated *fieldmap* was then aligned with rigid-registration to the target EPI (echo-planar imaging) reference run. The field coefficients were mapped on to the reference EPI using the transform. The BOLD reference was then co-registered to the T1w reference using bbregister (FreeSurfer) which implements boundary-based registration (Greve and Fischl 2009). Co-registration was configured with six degrees of freedom. Several confounding time-series were calculated based on the *preprocessed BOLD*: framewise displacement (FD), DVARS and three region-wise global signals. FD was computed using two formulations following Power (absolute sum of relative motions, Power et al. (2014)) and Jenkinson (relative root mean square displacement between affines, Jenkinson et al. (2002)). FD and DVARS are calculated for each functional run, both using their implementations in *Nipype* (following the definitions by Power et al. 2014). The three global signals are extracted within the CSF, the WM, and the whole-brain masks. Additionally, a set of physiological regressors were extracted to allow for component-based noise correction (*CompCor*, Behzadi et al. 2007). Principal components are estimated after high-pass filtering the *preprocessed BOLD* time-series (using a discrete cosine filter with 128s cut-off) for the two *CompCor* variants: temporal (tCompCor) and anatomical (aCompCor). tCompCor components are then calculated from the top 2% variable voxels within the brain mask. For aCompCor, three probabilistic masks (CSF, WM and combined CSF+WM) are generated in anatomical space. The implementation differs from that of Behzadi et al. in that instead of eroding the masks by 2 pixels on BOLD space, a mask of pixels that likely contain a volume fraction of GM is subtracted from the aCompCor masks. This mask is obtained by dilating a GM mask extracted from the FreeSurfer's *aseg* segmentation, and it ensures components are not extracted from voxels containing a minimal fraction of GM. Finally, these masks are resampled into BOLD space and binarized by thresholding at 0.99 (as in the original implementation). Components are also calculated separately within the WM and CSF masks. For each CompCor decomposition, the *k* components with the largest singular values are retained, such that the retained components' time series are sufficient to explain 50 percent of variance across the nuisance mask (CSF, WM, combined, or temporal). The remaining components are dropped from consideration. The head-motion estimates calculated in the correction step were also placed within the corresponding confounds file. The confound time series derived from head motion estimates and global signals were expanded with the inclusion of temporal derivatives and quadratic terms for each (Satterthwaite et al. 2013). Frames that exceeded a threshold of 0.5 mm FD or 1.5 standardized DVARS were annotated as motion outliers. Additional nuisance timeseries are calculated by means of principal components analysis of the signal found within a thin band (*crown*) of voxels around the edge of the brain, as proposed by (Patriat, Reynolds, and Birn 2017). All resamplings can be performed with *a single interpolation step* by composing all the pertinent transformations (i.e. head-motion transform matrices, susceptibility distortion correction when available, and co-registrations to anatomical and output spaces). Gridded (volumetric) resamplings were performed using nitransforms, configured with cubic B-spline interpolation.

Many internal operations of *fMRIPrep* use *Nilearn* 0.10.2 (Abraham et al. 2014, RRID:SCR_001362), mostly within the functional processing workflow. For more details of the pipeline, see the section corresponding to workflows in *fMRIPrep*'s documentation.

**Copyright Waiver**

The above boilerplate text was automatically generated by fMRIPrep with the express intention that users should copy and paste this text into their manuscripts *unchanged*. It is released under the CC0 license.

### Supplementary Data

#### Multiple linear regression models (primary outcomes)

##### Supplementary Table 1. Linear model results for primary outcomes.

| Models with terms | Estimate | CI (lower) | CI (upper) | Std. Error | t value | Pr(>\|t\|) |
|---|---|---|---|---|---|---|
| **Total SCFAs per g feces** | | | | | | |
| *$R^2$ = 0.325; Adj. $R^2$ = 0.257; F-statistic = 4.81; P = 0.0005* | | | | | | |
| (Intercept) | 79.21 | -10.48 | 168.90 | 44.84 | 1.77 | 0.08 |
| Baseline value | 0.41 | 0.17 | 0.64 | 0.12 | 3.47 | 0.00 |
| Group (intervention) | 5.38 | -4.95 | 15.72 | 5.17 | 1.04 | 0.30 |
| Age | -1.04 | -2.35 | 0.27 | 0.65 | -1.59 | 0.12 |
| Sex (male) | 13.37 | 2.88 | 23.87 | 5.25 | 2.55 | 0.01 |
| BMI | 0.02 | -1.00 | 1.05 | 0.51 | 0.04 | 0.97 |
| Habitual fiber intake | 0.61 | -0.28 | 1.49 | 0.44 | 1.37 | 0.18 |
| **n-back dlPFC fMRI responses [1]** | | | | | | |
| *$R^2$ = 0.363; Adj. $R^2$ = 0.285; F-statistic = 4.66; P = 0.001* | | | | | | |
| (Intercept) | 0.31 | -0.47 | 1.10 | 0.39 | 0.80 | 0.43 |
| Baseline value | 0.38 | 0.05 | 0.71 | 0.16 | 2.30 | 0.03 |
| Group (intervention) | 0.08 | 0.00 | 0.16 | 0.04 | 1.91 | 0.06 |
| Age | 0.00 | -0.01 | 0.01 | 0.01 | -0.32 | 0.75 |
| Sex (male) | 0.13 | 0.04 | 0.22 | 0.04 | 2.89 | 0.01 |
| Education level [2] | 0.01 | -0.01 | 0.04 | 0.01 | 1.22 | 0.23 |
| Blood vitamin B6 (nmol/L) | 0.00 | 0.00 | 0.00 | 0.00 | -1.18 | 0.24 |
| **n-back hippocampus fMRI responses [1]** | | | | | | |
| *$R^2$ = 0.268; Adj. $R^2$ = 0.179; F-statistic = 3.00; P = 0.014* | | | | | | |
| (Intercept) | -0.16 | -0.49 | 0.16 | 0.16 | -1.02 | 0.31 |
| Baseline value | 0.36 | 0.08 | 0.64 | 0.14 | 2.55 | 0.01 |
| Group (intervention) | 0.05 | 0.01 | 0.08 | 0.02 | 2.55 | 0.01 |
| Age | 0.00 | 0.00 | 0.01 | 0.00 | 0.70 | 0.49 |
| Sex (male) | 0.03 | 0.00 | 0.07 | 0.02 | 1.79 | 0.08 |
| Education level [2] | 0.00 | -0.01 | 0.01 | 0.01 | 0.69 | 0.50 |
| Blood vitamin B6 (nmol/L) | 0.00 | 0.00 | 0.00 | 0.00 | -0.86 | 0.40 |
| **n-back performance (dprime 2b)** | | | | | | |
| *$R^2$ = 0.430; Adj. $R^2$ = 0.368; F-statistic = 6.91; P = 0.000* | | | | | | |
| (Intercept) | 4.34 | 1.78 | 6.90 | 1.28 | 3.39 | 0.00 |
| Baseline value | 0.60 | 0.39 | 0.80 | 0.10 | 5.93 | 0.00 |
| Group (intervention) | 0.10 | -0.19 | 0.40 | 0.15 | 0.68 | 0.50 |
| Age | -0.05 | -0.08 | -0.01 | 0.02 | -2.48 | 0.02 |
| Sex (male) | 0.06 | -0.24 | 0.35 | 0.15 | 0.38 | 0.71 |
| Education level [2] | 0.07 | -0.02 | 0.16 | 0.04 | 1.51 | 0.14 |

| | | | | | | |
|---|---|---|---|---|---|---|
| Blood vitamin B6 (nmol/L) | 0.00 | -0.01 | 0.00 | 0.00 | -1.77 | 0.08 |

dlPFC = dorsolateral prefrontal cortex; SCFAs = short-chain fatty acids.

[1] Contrast in blood-oxygen level dependent (BOLD) response between 2-back and 0-back condition of the n-back task, reflecting working memory related activation.

[2] Education level ranges from 'not possessing a primary school diploma' (score 0) to 'possessing a PhD degree' (score 8).

### Basic model (reported outcomes)

**Supplementary Table 2. Intervention-induced effects in placebo and CDMV group on the fecal short-chain fatty acids and fMRI working memory n-back task in older adults at risk of cognitive decline.**

| **Outcome** | **Placebo** | | **CDMV** | | **Estimate group (CI)** | **P-value** |
|---|---|---|---|---|---|---|
| | **Mean$_{adj}$ (SE)** | ***n*** | **Mean$_{adj}$ (SE)** | ***n*** | | |
| **Fecal short-chain fatty acids** | | | | | | |
| Total fecal short-chain fatty acids (µmol/g feces) | 53.9 (3.56) | 36 | 57.7 (3.84) | 31 | 3.74 (-6.68; 14.2) | 0.48 |
| Acetic acid (µmol/g feces) | 32.0 (2.04) | 36 | 33.3 (2.21) | 31 | 1.32 (-4.75; 7.38) | 0.67 |
| Propionic acid (µmol/g feces) | 9.94 (0.71) | 36 | 10.4 (0.77) | 31 | 0.493 (-1.62; 2.61) | 0.64 |
| Butyric acid (µmol/g feces) | 9.36 (0.91) | 36 | 11.2 (0.98) | 31 | 1.85 (-0.852; 4.54) | 0.18 |
| Valeric acid (µmol/g feces) * | 0.18 (0.03) *1.57 (0.10)* | 36 | 0.23 (0.03) *1.75 (0.12)* | 31 | 0.05 (-0.03; 0.13) | 0.25 |
| **Working memory fMRI responses[1]** | | | | | | |
| *dlPFC (bilateral)* | 0.40 (0.03) | 29 | 0.45 (0.03) | 27 | 0.05 (-0.03; 0.13) | 0.22 |
| *dlPFC (right)* | 0.36 (0.03) | 29 | 0.42 (0.03) | 27 | 0.06 | 0.19 |

| | | | | | | |
|---|---|---|---|---|---|---|
| | | | | | (-0.03; 0.14) | |
| *dlPFC (left)* | 0.45 (0.03) | 29 | 0.50 (0.03) | 27 | 0.05<br>(-0.04; 0.14) | 0.27 |
| *Hippocampus (bilateral)* | -0.06 (0.01) | 29 | -0.02 (0.01) | 27 | 0.04<br>(0.00; 0.07) | **0.03*** |
| *Hippocampus (right)* | -0.06 (0.01) | 29 | 0.00 (0.01) | 27 | 0.07<br>(0.02; 0.09) | **0.00*** |
| *Hippocampus (left)* | -0.07 (0.01) | 29 | -0.04 (0.01) | 27 | 0.02<br>(-0.02; 0.06) | 0.23 |
| **Working memory fMRI functional connectivity with dlPFC[1]** | | | | | | |
| *Hippocampus (bilateral)* | -0.09 (0.01) | 29 | -0.06 (0.01) | 27 | 0.04<br>(0.00; 0.07) | 0.06 |
| *Hippocampus (right)* | -0.09 (0.01) | 29 | -0.04 (0.01) | 27 | 0.05<br>(0.02; 0.09) | **0.01*** |
| *Hippocampus (left)* | -0.09 (0.02) | 29 | -0.07 (0.02) | 27 | 0.02<br>(-0.02; 0.06) | 0.38 |
| **Working memory performance** | | | | | | |
| *Dprime 1-back* | 3.83 (0.12) | 34 | 3.74 (0.13) | 28 | -0.09<br>(-0.43; 0.35) | 0.60 |
| *Dprime 2-back* | 2.81 (0.10) | 34 | 2.79 (0.11) | 28 | -0.02<br>(-0.32; 0.27) | 0.87 |

CDMV = colon-delivered multivitamin.

[1]Contrast in blood-oxygen level dependent (BOLD) response between 2-back and 0-back condition of the n-back task, reflecting working memory related activation.

Estimated marginal means with standard error at week 6 are reported for the basic linear model (adjusted for baseline value only). Parameter estimate and confidence interval for CDMV group are reported. Primary outcomes are underlined. Outcomes indicated with an asterisk (*) are log-transformed (log10) for normal distribution; original (upper) and back-transformed (lower) adjusted means and standard errors are reported for these outcomes.

### Habitual dietary intake and blood vitamin concentrations

**Supplementary table 3. Habitual dietary daily intake of macronutrients and vitamins that are also present in the colon-delivery supplement.**

| **Daily intake** | **Placebo** | | | | **CDMV** | | | | **Between groups** |
|---|---|---|---|---|---|---|---|---|---|
| | ***n*** | **Week 0 Mean ± SD** | **Week 6 Mean ± SD** | **Within group change** | ***n*** | **Week 0 Mean ± SD** | **Week 6 Mean ± SD** | **Within group change** | **P-value** |
| Energy (kJ) | 33 | 8108 ± 1990 | 8193 ± 2234 | 84.8 ± 1360 | 28 | 8338 ± 3252 | 7856 ± 4623 | -320 ± 2160 | 0.36 |
| Total carbohydrates (EN%) | 33 | 38.0 ± 5.84 | 38.6 ± 5.75 | 0.444 ± 3.72 | 28 | 38.5 ± 7.54 | 37.9 ± 6.86 | -0.397 ± 4.98 | 0.44 |
| Total fat (EN%) | 33 | 38.9 ± 6.09 | 39.3 ± 6.42 | -0.0538 ± 3.96 | 28 | 40.0 ± 7.49 | 40.2 ± 7.42 | 0.12 ± 5.6 | 0.42 |
| Total protein (EN%) | 33 | 15.9 ± 2.34 | 15.3 ± 2.11 | -0.515 ± 1.91 | 28 | 15.2 ± 2.32 | 15.6 ± 2.51 | 0.332 ± 1.77 | 0.12 |
| Total fiber (EN%) | 33 | 2.25 ± 0.60 | 2.16 ± 0.53 | -0.112 ± 0.434 | 28 | 2.31 ± 0.41 | 2.28 ± 0.49 | -0.0547 ± 0.447 | 0.42 |
| Total fiber (gram) | 33 | 21.6 ± 7.59 | 20.5 ± 6.06 | -1.26 ± 4.36 | 28 | 22.5 ± 7.73 | 19.5 ± 5.16 | -2.78 ± 5.27 | 0.16 |
| Vitamin B2 (mg) | 33 | 1.42 ± 0.64 | 1.48 ± 0.80 | 0.0412 ± 0.392 | 28 | 1.33 ± 0.52 | 1.32 ± 1.05 | 0.00679 ± 0.649 | 0.99 |

| | | | | | | | | | |
|---|---|---|---|---|---|---|---|---|---|
| Vitamin B6 (mg) | 33 | 1.53 ± 0.47 | 1.51 ± 0.50 | 0.00152 ± 0.349 | 28 | 1.47 ± 0.45 | 1.42 ± 0.60 | -0.0204 ± 0.418 | 0.75 |
| Vitamin B9 (µg) | 33 | 250 ± 64.6 | 243 ± 61.1 | -9.78 ± 47 | 28 | 252 ± 87.7 | 246 ± 138.5 | -8.73 ± 77.5 | 0.95 |
| Vitamin C (mg) | 33 | 81.7 ± 37.5 | 80.9 ± 29.8 | -3.33 ± 32.6 | 28 | 91.6 ± 39.8 | 77.4 ± 39.3 | -12.5 ± 35.7 | 0.39 |
| Vitamin D (µg) | 33 | 3.10 ± 1.20 | 2.96 ± 1.12 | -0.093 ± 0.773 | 28 | 3.17 ± 1.46 | 2.92 ± 1.39 | -0.209 ± 0.929 | 0.63 |

CDMV = colon-delivered multivitamin

Outcomes indicated with an asterisk (*) are log-transformed (log10) for normal distribution. Means+SD at week 0 and week 6 are non-adjusted. Between-groups p-values are based on linear model *lm(post-value ~ baseline-value + group)*. It is not possible to quantify vitamin B3 habitual intake using a food frequency questionnaire.

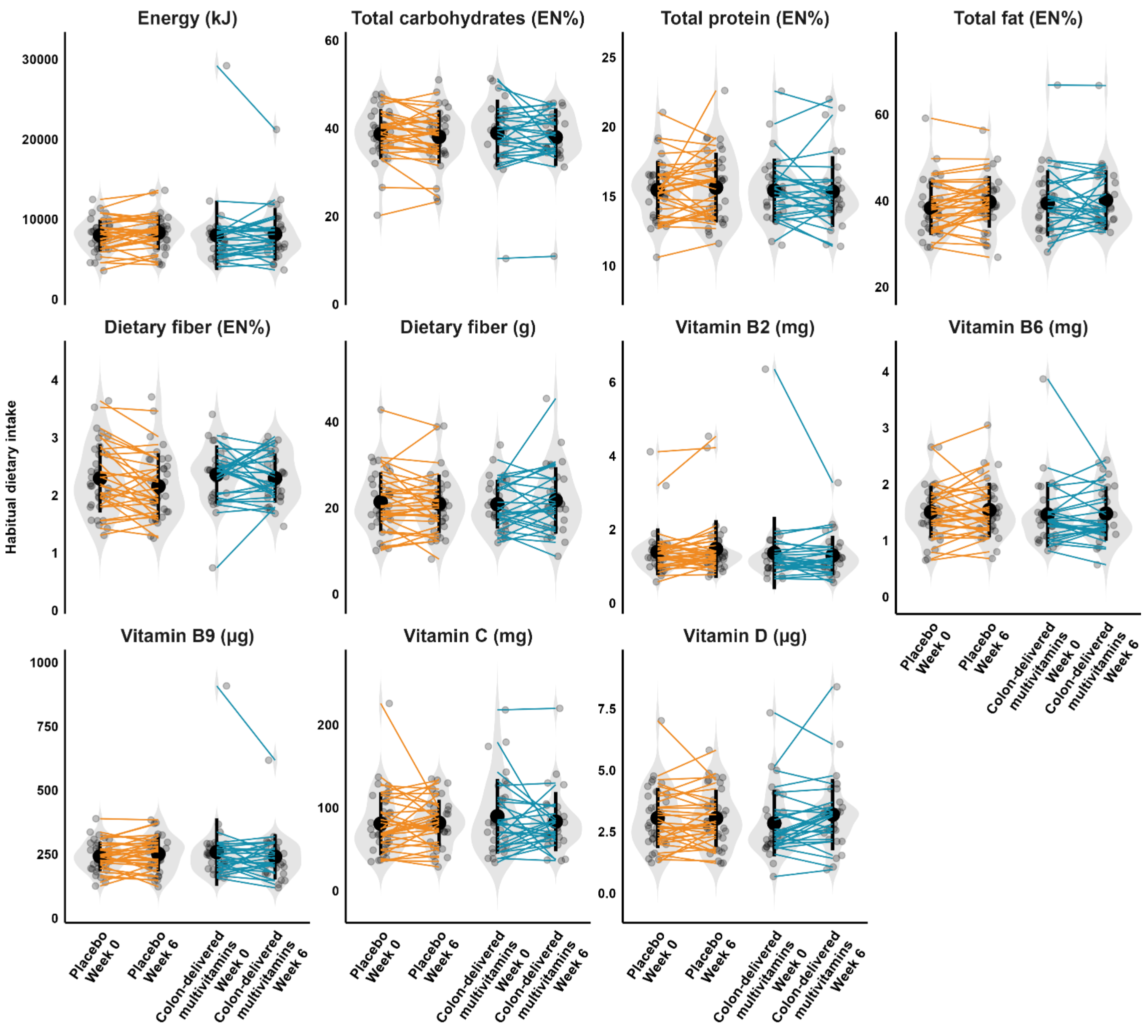


**Supplementary Figure 1**. **Habitual dietary daily intake at baseline and after the 6-weeks intervention for placebo (*n*=33) and CDMV group (*n*=28).** *Habitual dietary intake of macronutrients and vitamins that are also present in the colon-delivery supplement is reported. Individual paired samples are connected by a line. The width of the violin shapes indicates the sample density, the circled shape inside indicates the group mean and SD. CDMV = colon-delivered multivitamin.*

***Supplementary table 4. Blood vitamin concentrations at baseline and after the 6-weeks intervention.***

| **Blood vitamin** | **Placebo** | | | | **CDMV** | | | | **Between groups** |
|---|---|---|---|---|---|---|---|---|---|
| | ***n*** | **Week 0 Mean ± SD** | **Week 6 Mean ± SD** | **Within group change** | ***n*** | **Week 0 Mean ± SD** | **Week 6 Mean ± SD** | **Within group change** | **P-value** |
| Vitamin B2 (nmol/L) * | 35 | 265 ± 29.9 | 269 ± 34.1 | 3.46 ± 21.1 | 31 | 264 ± 40.4 | 266 ± 35.1 | 1.63 ± 17 | 0.82 |
| Vitamin B3 (µmol/L) | 35 | 34.6 ± 5.78 | 35.1 ± 6.55 | 0.436 ± 5.49 | 29 | 32.5 ± 6.27 | 35.5 ± 5.77 | 2.96 ± 5.77 | 0.21 |
| Vitamin B6 (nmol/L) * | 35 | 85.9 ± 14.7 | 90.1 ± 33 | 4.21 ± 32.4 | 31 | 85.2 ± 16.1 | 105 ± 24.4 | 20.2 ± 19.4 | **0.00*** |
| Vitamin B9 (nmol/L) * | 35 | 30.9 ± 12.2 | 29.3 ± 11.5 | -1.69 ± 4.25 | 31 | 26 ± 9.77 | 24.9 ± 9.46 | -1.14 ± 3.63 | 0.67 |
| Vitamin C (µmol/L) | 35 | 64.8 ± 21.9 | 64.9 ± 29.8 | 0.673 ± 20.5 | 31 | 67.8 ± 21.8 | 70.6 ± 32.6 | 2.85 ± 23.6 | 0.70 |
| Total vitamin D (nmol/L) | 34 | 66.5 ± 18.8 | 64.5 ± 18.9 | -1.81 ± 9.57 | 31 | 76.9 ± 22.9 | 72.1 ± 22.9 | -4.77 ± 9.46 | 0.46 |

CDMV = colon-delivered multivitamin.

Outcomes indicated with an asterisk (*) are log-transformed (log10) for normal distribution. Means+SD at week 0 and week 6 are non-adjusted. Between-groups p-values are based on linear model *lm(post-value ~ baseline-value + group)*.

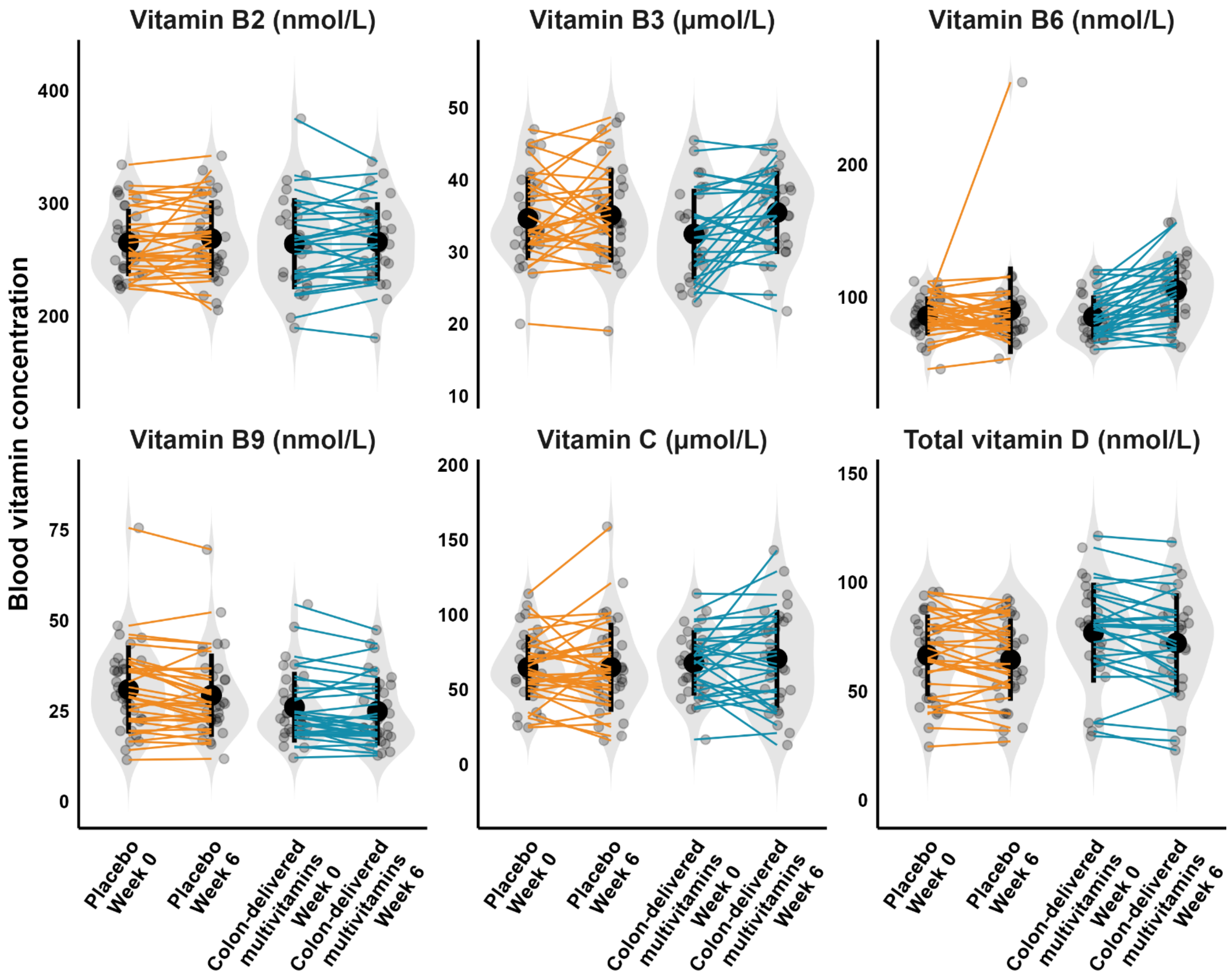


**Supplementary Figure 2. Blood vitamin concentrations at baseline and after the 6-weeks intervention for placebo and CDMV group (for *n* per group, see Supplementary Table 5).** *Individual paired samples are connected by a line. The width of the violin shapes indicates the sample density, the circled shape inside indicates the group mean and SD. CDMV = colon-delivered multivitamin.*

#### User experiences and adverse effects

##### Summary of adverse effects

No serious adverse events (SAEs) have occured within this study. There were 15 adverse events (AEs). Of these 15 AEs, 1 was classified as 'severe', 4 were classified as 'moderate', and 10 as 'mild'. The only 'severe' AE is not related to participation in the intervention. This concerned a participant with sleep apnea, where the sleep apnea equipment did not work properly for one night.

In total, 3 AEs were classified as 'definitely related' to participation, which were all related to events during the MRI scan (claustrophobia 1x and/or coughing 2x). 3 AEs were classified as 'possibly related' to participation, these three AEs may have been related to the intervention capsule (strange taste in mouth 1x, heartburn/bloated feeling 2x). All other AEs were classified as 'not/unlikely related' to participation, and varied in nature (cold 3x, bladder infection 1x, bone fracture 1x, red swollen eye 1x, high glucose values 1x, sleep apnea 1x, bloated feeling 1x).

**Supplementary table 5. User experiences of placebo and CDMV group.**

| **User experiences** | **Placebo** | | | **CDMV** | | |
|---|---|---|---|---|---|---|
| | ***n*** | **Yes (%)** | **No (%)** | ***n*** | **Yes (%)** | **No (%)** |
| *Experienced side effects* | 36 | 11.1 | 88.9 | 30 | 16.7 | 83.3 |
| *Experienced effects on stool and intestines* | 36 | 19.4 | 80.6 | 30 | 20.0 | 80.0 |
| *Experienced effects on memory and cognition* | 36 | 2.8 | 97.2 | 30 | 3.3 | 96.7 |

CDMV = colon-delivered multivitamin.

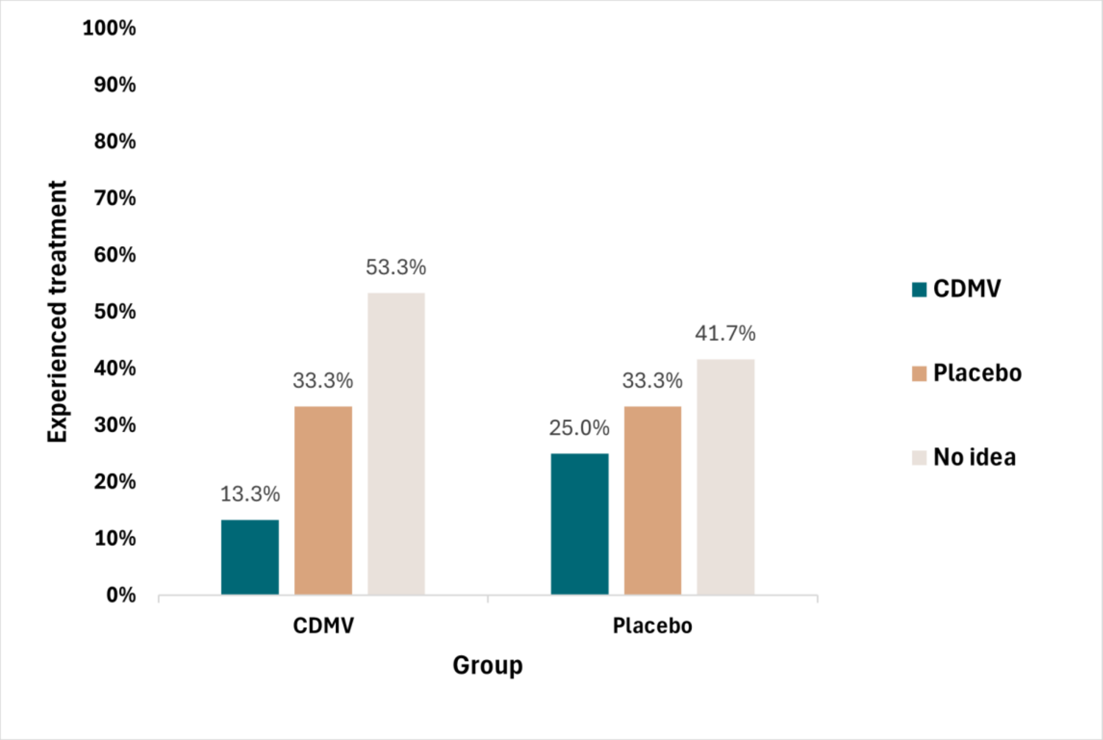


**Supplementary Figure 3. Experienced treatment per group.** Experienced treatment for placebo group (*n*=36) and CDMV group (*n*=31), expressed in percentage (%) of participants per group.

CDMV = colon-delivered multivitamin.

### Secondary outcomes

#### Gut-related outcomes

**Supplementary table 6. Intervention-induced effects in placebo and CDMV group on secondary gut-related outcomes in older adults at risk of cognitive decline.**

| Outcome | Basic linear model | | | | | | Final linear model with covariates | | | | | |
|---|---|---|---|---|---|---|---|---|---|---|---|---|
| | Placebo | | CDMV | | Estimate group (CI) | P-value | Placebo | | CDMV | | Estimate group (CI) | P-value |
| | Mean$_{adj}$ (SE) | *n* | Mean$_{adj}$ (SE) | *n* | | | Mean$_{adj}$ (SE) | *n* | Mean$_{adj}$ (SE) | *n* | | |
| **Fecal medium-chain fatty acids** | | | | | | | | | | | | |
| Total medium-chain fatty acids (µmol/g feces) * | -0.31 (0.04)<br>*0.52 (0.05)* | 36 | -0.19 (0.04)<br>*0.69 (0.07)* | 31 | 0.12<br>(0.00; 0.24) | **0.05** | -0.31 (0.04)<br>*0.53 (0.05)* | 36 | -0.19 (0.05)<br>*0.70 (0.07)* | 31 | 0.12<br>(-0.01; 0.25) | 0.06 |
| Hexanoic acid (µmol/g feces) * | -0.45 (0.06)<br>*0.39 (0.04)* | 36 | -0.31 (0.05)<br>*0.54 (0.04)* | 31 | 0.15<br>(0.01; 0.28) | **0.04** | -0.45 (0.05)<br>*0.39 (0.04)* | 36 | -0.30 (0.05)<br>*0.55 (0.04)* | 31 | 0.15<br>(0.00; 0.29) | **0.04** |
| Heptanoic acid (µmol/g feces) * | -0.93 (0.05)<br>*0.13 (0.02)* | 36 | -0.87 (0.06)<br>*0.15 (0.02)* | 31 | 0.06<br>(-0.10; 0.21) | 0.47 | -0.92 (0.05)<br>*0.13 (0.02)* | 36 | -0.86 (0.06)<br>*0.15 (0.02)* | 31 | 0.06<br>(-0.10; 0.21) | 0.47 |
| **Fecal branched-chain fatty acids** | | | | | | | | | | | | |
| Total branched-chain fatty acids (µmol/g feces) * | 0.59 (0.02)<br>*4.00 (0.21)* | 36 | 0.62 (0.02)<br>*4.27 (0.24)* | 31 | 0.03<br>(-0.04; 0.10) | 0.41 | 0.59 (0.02)<br>*4.00 (0.22)* | 36 | 0.62 (0.03)<br>*4.27 (0.26)* | 31 | 0.03<br>(-0.04; 0.10) | 0.42 |
| Isovaleric acid (µmol/g feces) * | 0.40 (0.02) | 36 | 0.42 (0.02) | 31 | 0.02 | 0.49 | 0.40 (0.02) | 36 | 0.42 (0.03) | 31 | 0.02 | 0.51 |

| | | | | | | | | | | | | |
|---|---|---|---|---|---|---|---|---|---|---|---|---|
| | *2.57 (0.14)* | | *2.72 (0.16)* | | (-0.04; 0.09) | | *2.57 (0.14)* | | *2.72 (0.16)* | | (-0.05; 0.10) | |
| Isobutyric acid (µmol/g feces) * | 0.10 (0.03) | 36 | 0.16 (0.03) | 31 | 0.05 | 0.17 | 0.10 (0.03) | 36 | 0.16 (0.03) | 31 | 0.06 | 0.16 |
| | *1.30 (0.08)* | | *1.47 (0.09)* | | (-0.02; 0.13) | | *1.30 (0.08)* | | *1.48 (0.10)* | | (-0.02; 0.14) | |
| 4-methyl valeric acid (µmol/g | -1.34 (0.10) | 36 | -1.18 (0.11) | 31 | 0.16 | 0.30 | -1.30 (0.11) | 36 | -1.18 (0.12) | 31 | 0.12 | 0.43 |
| feces) * | *0.07 (0.02)* | | *0.10 (0.03)* | | (-0.15; 0.47) | | *0.08 (0.02)* | | *0.10 (0.03)* | | (-0.19; 0.44) | |
| **Blood short-chain fatty acids** | | | | | | | | | | | | |
| Total short-chain fatty acids (µM) * | 2.13 (0.02) | 34 | 2.10 (0.02) | 31 | -0.02 | 0.41 | 2.13 (0.02) | 34 | 2.11 (0.02) | 31 | -0.02 | 0.55 |
| | *135.75 (5.78)* | | *128.99 (5.75)* | | (-0.08; 0.03) | | *135.66 (6.01)* | | *130.45 (6.17)* | | (-0.07; 0.04) | |
| Acetic acid (µM) * | 2.12 (0.02) | 34 | 2.09 (0.02) | 31 | -0.02 | 0.40 | 2.12 (0.02) | 34 | 2.10 (0.02) | 31 | -0.02 | 0.55 |
| | *132.50 (5.67)* | | *125.67 (5.64)* | | (-0.08; 0.03) | | *132.35 (5.90)* | | *127.15 (6.05)* | | (-0.07; 0.04) | |
| Propionic acid (µM) * | 0.30 (0.02) | 34 | 0.29 (0.02) | 31 | -0.01 | 0.72 | 0.31 (0.02) | 34 | 0.29 (0.02) | 31 | -0.02 | 0.53 |
| | *2.02 (0.09)* | | *1.97 (0.09)* | | (-0.07; 0.05) | | *2.05 (0.09)* | | *1.97 (0.09)* | | (-0.08; 0.04) | |
| Butyric acid (µM) * | -0.02 (0.03) | 34 | -0.03 (0.03) | 31 | -0.01 | 0.80 | -0.02 (0.03) | 34 | -0.03 (0.03) | 31 | -0.01 | 0.84 |
| | *0.98 (0.06)* | | *0.96 (0.07)* | | (-0.10; 0.08) | | *0.99 (0.07)* | | *0.97 (0.07)* | | (-0.10; 0.08) | |
| Valeric acid (µM) * | -0.74 (0.13) | 10 | -0.93 (0.10) | 16 | -0.20 | 0.23 | -0.79 (0.15) | 10 | -0.93 (0.13) | 16 | -0.14 | 0.48 |
| | *0.22 (0.06)* | | *0.14 (0.03)* | | (-0.53; 0.14) | | *0.20 (0.07)* | | *0.14 (0.04)* | | (-0.55; 0.27) | |
| **Blood medium-chain fatty acids** | | | | | | | | | | | | |
| Total medium-chain fatty acids | 0.51 (0.02) | 34 | 0.55 (0.02) | 31 | 0.04 | 0.23 | 0.51 (0.02) | 34 | 0.54 (0.02) | 31 | 0.03 | 0.34 |
| (µM) * | *3.28 (0.16)* | | *3.57 (0.18)* | | (-0.02; 0.10) | | *3.30 (0.17)* | | *3.54 (0.19)* | | (-0.03; 0.10) | |
| Hexanoic acid (µM) * | -0.31 (0.02) | 34 | -0.32 (0.02) | 31 | -0.01 | 0.77 | -0.31 (0.02) | 34 | -0.32 (0.02) | 31 | -0.01 | 0.78 |
| | *0.50 (0.02)* | | *0.49 (0.02)* | | (-0.07; 0.05) | | *0.50 (0.02)* | | *0.49 (0.03)* | | (-0.07; 0.05) | |
| Heptanoic acid (µM) * | -1.12 (0.04) | 34 | -1.13 (0.04) | 31 | -0.01 | 0.86 | -1.11 (0.04) | 34 | -1.14 (0.05) | 31 | 0.03 | 0.67 |

| | | | | | | | | | | | | |
|---|---|---|---|---|---|---|---|---|---|---|---|---|
| | *0.08 (0.01)* | | *0.08 (0.01)* | | (-0.13; 0.11) | | *0.08 (0.01)* | | *0.08 (0.01)* | | (-0.16; 0.10) | |
| Octanoic acid (μM) * | -0.22 (0.02)<br>*0.61 (0.03)* | 34 | -0.21 (0.02)<br>*0.63 (0.03)* | 31 | 0.01<br>(-0.05; 0.08) | 0.68 | -0.22 (0.02)<br>*0.61 (0.03)* | 34 | -0.21 (0.02)<br>*0.62 (0.04)* | 31 | 0.01<br>(-0.06; 0.08) | 0.83 |
| Decanoic acid (μM) * | -0.20 (0.02)<br>*0.65 (0.04)* | 34 | -0.14 (0.02)<br>*0.74 (0.04)* | 31 | 0.06<br>(-0.01; 0.13) | 0.09 | -0.19 (0.02)<br>*0.66 (0.04)* | 34 | -0.14 (0.03)<br>*0.75 (0.04)* | 31 | 0.05<br>(-0.02; 0.13) | 0.14 |
| Dodecanoic acid (μM) * | 0.12 (0.03)<br>*1.36 (0.09)* | 34 | 0.18 (0.03)<br>*1.57 (0.11)* | 31 | 0.06<br>(-0.02; 0.15) | 0.15 | 0.12 (0.03)<br>*1.37 (0.10)* | 34 | 0.17 (0.03)<br>*1.55 (0.10)* | 31 | 0.05<br>(-0.04; 0.14) | 0.25 |
| **Blood branched-chain fatty acids** | | | | | | | | | | | | |
| Total branched-chain fatty acids (μM) * | 0.29 (0.02)<br>*2.00 (0.09)* | 34 | 0.26 (0.02)<br>*1.83 (0.09)* | 31 | -0.04<br>(-0.09; 0.02) | 0.20 | 0.30 (0.02)<br>*2.01 (0.09)* | 34 | 0.26 (0.02)<br>*1.84 (0.09)* | 31 | -0.04<br>(-0.10; 0.02) | 0.22 |
| Isovaleric acid (μM) * | 0.03 (0.02)<br>*1.08 (0.06)* | 34 | -0.02 (0.02)<br>*0.97 (0.05)* | 31 | -0.05<br>(-0.11; 0.02) | 0.15 | 0.02 (0.02)<br>*1.08 (0.06)* | 34 | -0.01 (0.02)<br>*0.99 (0.06)* | 31 | -0.04<br>(-0.11; 0.03) | 0.25 |
| Isobutyric acid (μM) * | -0.28 (0.02)<br>*0.53 (0.02)* | 34 | -0.29 (0.02)<br>*0.52 (0.03)* | 31 | -0.01<br>(-0.07; 0.05) | 0.80 | -0.28 (0.02)<br>*0.54 (0.02)* | 34 | -0.30 (0.02)<br>*0.51 (0.02)* | 31 | -0.02<br>(-0.08; 0.04) | 0.50 |
| 2-methyl butyric acid (μM) * | -0.45 (0.03)<br>*0.36 (0.02)* | 34 | -0.50 (0.03)<br>*0.33 (0.02)* | 31 | 0.05<br>(-0.13; 0.04) | 0.26 | -0.45 (0.03)<br>*0.37 (0.02)* | 34 | -0.50 (0.03)<br>*0.33 (0.02)* | 31 | 0.06<br>(-0.14; 0.03) | 0.20 |
| **Fecal microbiota alpha-diversity** | | | | | | | | | | | | |
| Phylogenetic Diversity | 27.9 (0.39) | 36 | 27.8 (0.42) | 31 | -0.0780<br>(-1.22; 1.06) | 0.89 | 27.9 (0.39) | 36 | 27.6 (0.43) | 31 | -0.257<br>(-1.43; 0.920) | 0.66 |
| Shannon Diversity | 4.47 (0.06) | 36 | 4.39 (0.07) | 31 | -0.0864<br>(-0.274; 0.101) | 0.36 | 4.47 (0.06) | 36 | 4.33 (0.07) | 31 | -0.136<br>(-0.317; 0.0440) | 0.14 |

| | | | | | | | | | | | | |
|---|---|---|---|---|---|---|---|---|---|---|---|---|
| Chao1 | 342 (5.99) | 36 | 331 (6.45) | 31 | -10.7<br>(-28.3; 6.92) | 0.23 | 341 (6.02) | 36 | 327 (6.60) | 31 | -14.6<br>(-32.5; 3.36) | 0.11 |
| **Feces characteristics** | | | | | | | | | | | | |
| Fecal pH | 7.16 (0.06) | 36 | 7.03 (0.07) | 31 | -0.13<br>(-0.32; 0.06) | 0.17 | 7.13 (0.06) | 36 | 7.01 (0.07) | 31 | -0.12<br>(-0.31; 0.07) | 0.23 |
| Fecal redox potential (ORP) | 88.4 (8.43) | 36 | 90.6 (9.09) | 31 | 2.23<br>(-22.56; 27.03) | 0.86 | 88.5 (8.59) | 36 | 87.3 (9.43) | 31 | -1.18<br>(-26.81; 24.45) | 0.93 |
| **Intestinal permeability** | | | | | | | | | | | | |
| Blood zonulin (ng/mL) | 32.1 (1.40) | 35 | 30.8 (1.49) | 31 | -1.33<br>(-5.48; 2.82) | 0.52 | 32.5 (1.44) | 35 | 30.1 (1.55) | 31 | -2.45<br>(-6.78; 1.88) | 0.26 |
| Blood LBP (µg/mL) * | 1.06 (0.02)<br>*11.65 (0.48)* | 33 | 1.08 (0.02)<br>*12.22 (0.52)* | 31 | 0.02<br>(-0.03; 0.07) | 0.43 | 1.07 (0.02)<br>*11.83 (0.48)* | 33 | 1.07 (0.02)<br>*11.99 (0.52)* | 31 | 0.01<br>(-0.05; 0.06) | 0.82 |
| **Intestinal inflammation** | | | | | | | | | | | | |
| Fecal lipocalin | 0.39 (0.04) | 28 | 0.39 (0.04) | 24 | 0.00<br>(-0.12; 0.11) | 0.97 | 0.38 (0.04) | 28 | 0.40 (0.04) | 24 | 0.03<br>(-0.09; 0.15) | 0.64 |
| **Stool consistency and transit time** | | | | | | | | | | | | |
| Fecal water content (%) | 71.51 (0.87) | 32 | 71.53 (0.94) | 27 | 0.02<br>(-2.55; 2.59) | 0.99 | 71.36 (0.90) | 32 | 71.46 (1.01) | 27 | 0.09<br>(-2.65; 2.84) | 0.95 |
| Gastrointestinal transit time (hours) * | 1.42 (0.04)<br>*27.54 (2.24)* | 35 | 1.32 (0.04)<br>*22.04 (1.93)* | 30 | -0.10<br>(-0.20; 0.01) | 0.07 | 1.42 (0.04)<br>*27.92 (2.33)* | 35 | 1.32 (0.04)<br>*22.01 (2.02)* | 30 | -0.10<br>(-0.21; 0.01) | 0.06 |

| | | | | | | | | | | | |
|---|---|---|---|---|---|---|---|---|---|---|---|
| Stool consistency (Bristol stool score from 1 to 7) | 3.68 (0.23) | 36 | 3.47 (0.25) | 31 | -0.21 (-0.89; 0.47) | 0.54 | 3.64 (0.22) | 36 | 3.63 (0.24) | 31 | -0.01 (-0.67; 0.65) | 0.98 |

CDMV = colon-delivered multivitamin; LBP = lipopolysaccharide binding protein; ORP = oxidation-reduction potential.

Estimated marginal means with standard error at week 6 are reported for basic linear model (adjusted for baseline value only) and final linear model (adjusted for age, sex, BMI and habitual dietary fiber intake). Parameter estimate and confidence interval for CDMV group are reported. Outcomes indicated with an asterisk (*) are log-transformed (log10) for normal distribution; original (upper) and back-transformed (lower) adjusted means and standard errors are reported for these outcomes. Non-adjusted means at week 0 and week 6 can be found in **Supplementary table 10**.

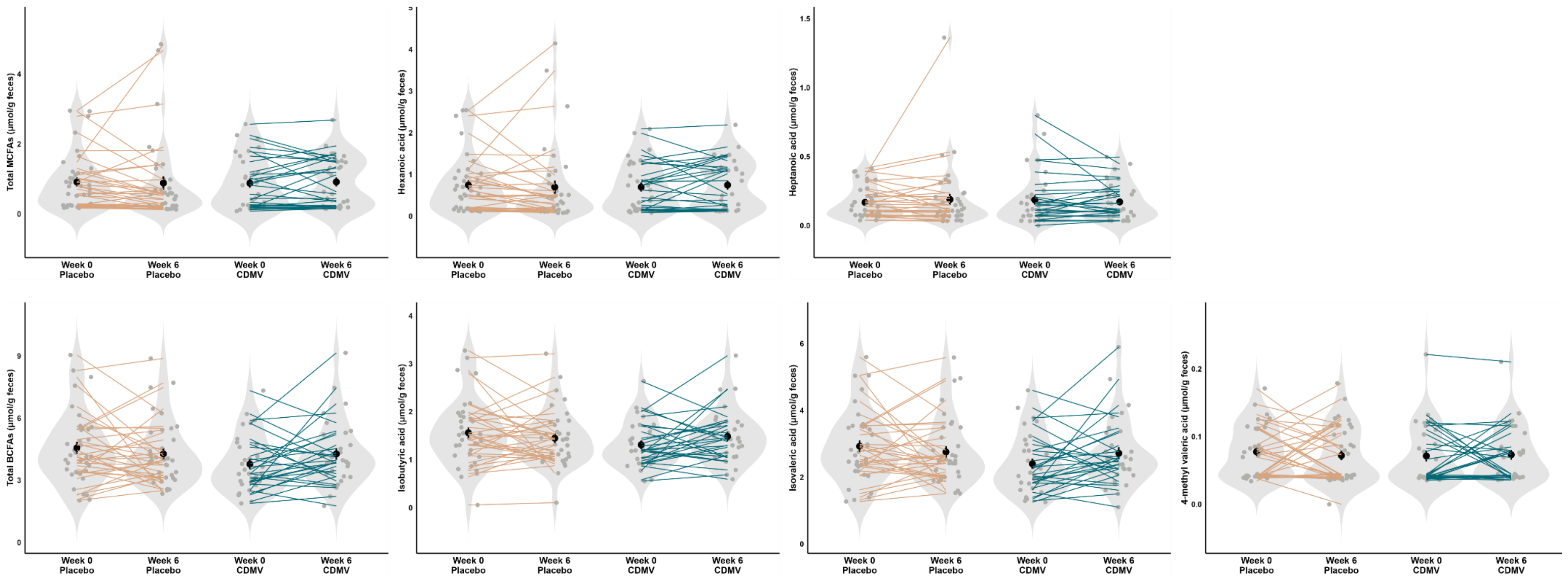


**Supplementary Figure 4. Fecal MCFAs and BCFAs at baseline and after the 6-weeks intervention for placebo (*n*=36) and CDMV group (*n*=31).** *Individual paired samples are connected by a line. The width of the violin shapes indicates the sample density, the circled shape inside indicates the group mean and SE. BCFA = branched-chain fatty acid; CDMV = colon-delivered multivitamin; MCFA = medium-chain fatty acid.*

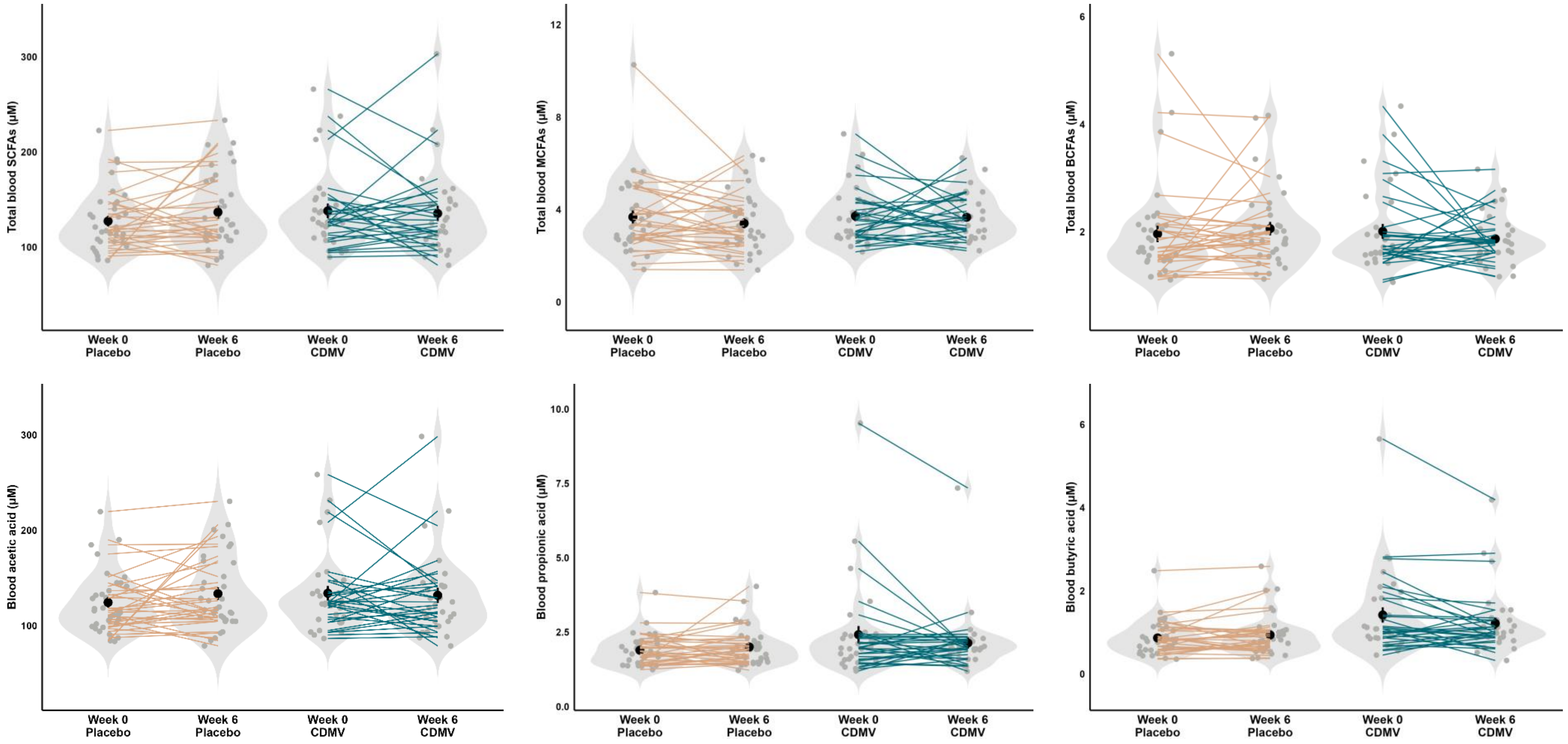


**Supplementary Figure 5. Blood SCFAs, MCFAs, and BCFAs at baseline and after the 6-weeks intervention for placebo (*n*=34) and CDMV group (*n*=31).** *Individual paired samples are connected by a line. The width of the violin shapes indicates the sample density, the circled shape inside indicates the group mean and SE. BCFA = branched-chain fatty acid; CDMV = colon-delivered multivitamin; MCFA = medium-chain fatty acid; SCFA = short-chain fatty acid.*

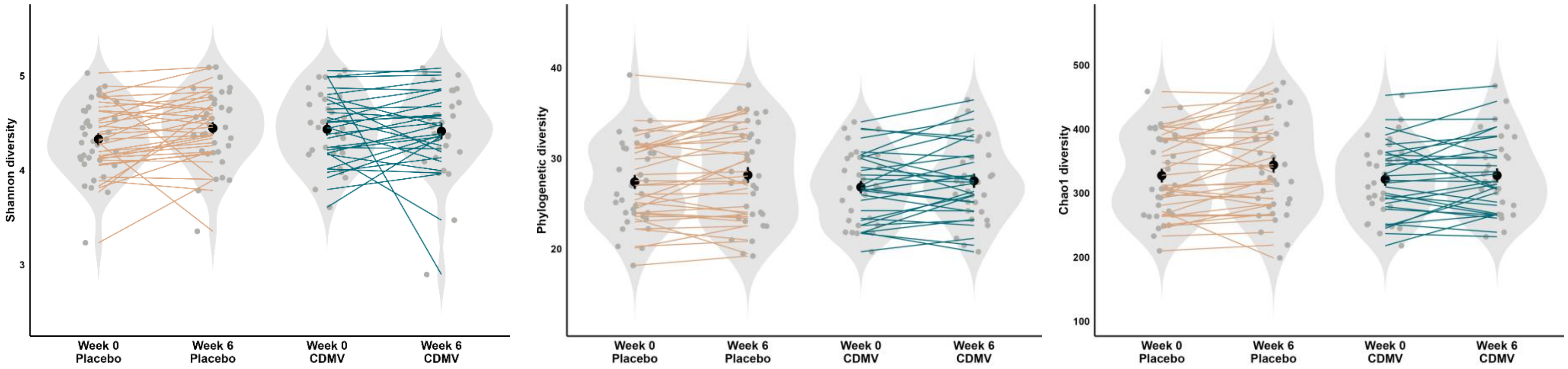


**Supplementary Figure 6**. **Fecal microbiota diversity (Shannon, phylogenetic, and Chao1) at baseline and after the 6-weeks intervention for placebo (*n*=36) and CDMV group (*n*=31)**. *Individual paired samples are connected by a line. The width of the violin shapes indicates the sample density, the circled shape inside indicates the group mean and SE. CDMV = colon-delivered multivitamin.*

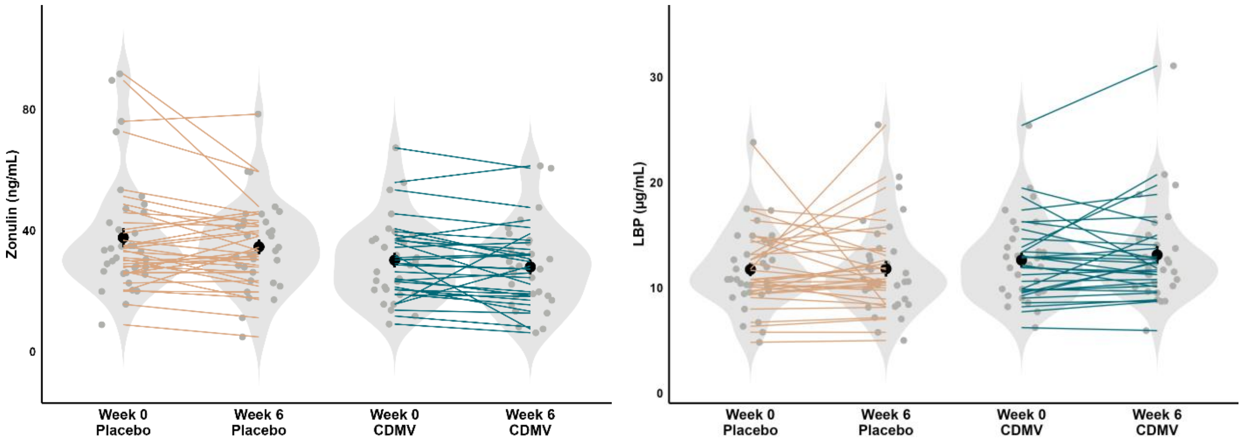


**Supplementary Figure 7**. **Blood intestinal permeability markers (zonulin and LBP) at baseline and after the 6-weeks intervention for placebo and CDMV group (for *n* per group, see Supplementary Table 6).** *Individual paired samples are connected by a line. The width of the violin shapes indicates the sample density, the circled shape inside indicates the group mean and SE. CDMV = colon-delivered multivitamin; LBP = lipopolysaccharide binding protein.*

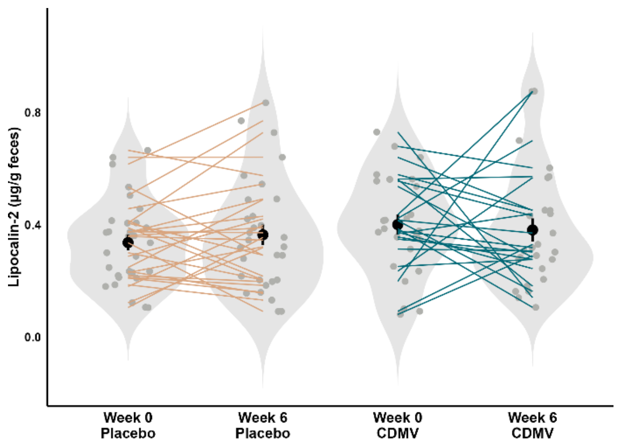


**Supplementary Figure 8. Intestinal inflammation markers measured in feces (lipocalin) at baseline and after the 6-weeks intervention for placebo and CDMV group (for *n* per group, see Supplementary Table 6).** *Individual paired samples are connected by a line. The width of the violin shapes indicates the sample density, the circled shape inside indicates the group mean and SE. CDMV = colon-delivered multivitamin.*

#### Systemic inflammation outcomes

**Supplementary table 7. Intervention-induced effects in placebo and CDMV group on secondary systemic inflammation outcomes in older adults at risk of cognitive decline.**

| Outcome | Basic linear model | | | | | | Final linear model with covariates | | | | | |
|---|---|---|---|---|---|---|---|---|---|---|---|---|
| | Placebo | | CDMV | | Estimate group (CI) | P-value | Placebo | | CDMV | | Estimate group (CI) | P-value |
| | $Mean_{adj}$ (SE) | *n* | $Mean_{adj}$ (SE) | *n* | | | $Mean_{adj}$ (SE) | *n* | $Mean_{adj}$ (SE) | *n* | | |
| Blood hs-CRP (mg/L)* | 0.21 (0.04) | 30 | 0.24 (0.04) | 27 | 0.03 | 0.59 | 0.21 (0.04) | 30 | 0.23 (0.04) | 27 | 0.02 | 0.70 |
| | *1.70 (0.14)* | | *1.82 (0.16)* | | (-0.08; 0.13) | | *1.69 (0.14)* | | *1.77 (0.16)* | | (-0.08; 0.12) | |
| Blood total WBC (counts×$10^9$/L)* | 0.76 (0.02) | 35 | 0.76 (0.02) | 29 | 0.00 | 0.99 | 0.76 (0.02) | 35 | 0.76 (0.02) | 29 | 0.00 | 0.96 |
| | *5.84 (0.24)* | | *5.84 (0.27)* | | (-0.05; 0.05) | | *5.89 (0.25)* | | *5.87 (0.28)* | | (-0.06; 0.05) | |
| Blood IFN-γ (pg/mL)* | 0.55 (0.04) | 31 | 0.57 (0.04) | 30 | 0.02 | 0.67 | 0.56 (0.04) | 31 | 0.58 (0.04) | 30 | 0.02 | 0.76 |
| | *3.75 (0.32)* | | *3.95 (0.35)* | | (-0.08; 0.13) | | *3.85 (0.33)* | | *4.00 (0.35)* | | (-0.09; 0.12) | |
| Blood IL-6 (pg/mL)* | -0.24 (0.04) | 31 | -0.22 (0.05) | 27 | 0.01 | 0.82 | -0.23 (0.04) | 31 | -0.23 (0.05) | 27 | 0.00 | 0.94 |
| | *0.62 (0.06)* | | *0.64 (0.07)* | | (-0.11; 0.14) | | *0.63 (0.06)* | | *0.63 (0.07)* | | (-0.14; 0.13) | |
| Blood IL-8 (pg/mL)* | 0.55 (0.02) | 33 | 0.55 (0.02) | 30 | 0.01 | 0.78 | 0.55 (0.02) | 33 | 0.55 (0.02) | 30 | 0.00 | 0.99 |
| | *3.55 (0.13)* | | *3.60 (0.14)* | | (-0.04; 0.05) | | *3.58 (0.14)* | | *3.58 (0.15)* | | (-0.05; 0.05) | |
| Blood IL-10 (pg/mL)* | -0.71 (0.04) | 33 | -0.65 (0.04) | 31 | 0.06 | 0.26 | -0.70 (0.04) | 33 | -0.64 (0.04) | 31 | 0.06 | 0.23 |
| | *0.21 (0.02)* | | *0.24 (0.02)* | | (-0.04; 0.16) | | *0.21 (0.02)* | | *0.24 (0.02)* | | (-0.04; 0.16) | |
| Blood TNF-α (pg/mL)* | 0.06 (0.01) | 32 | 0.06 (0.01) | 31 | 0.01 | 0.72 | 0.06 (0.01) | 32 | 0.07 (0.01) | 31 | 0.01 | 0.68 |
| | *1.15 (0.04)* | | *1.17 (0.04)* | | (-0.03; 0.05) | | *1.15 (0.04)* | | *1.17 (0.04)* | | (-0.03; 0.05) | |

CDMV = colon-delivered multivitamin; hs-CRP = high-sensitivity C-reactive protein; WBC = white blood cell.

Estimated marginal means with standard error at week 6 are reported for basic linear model (adjusted for baseline value only) and final linear model (adjusted for age, sex, and BMI). Parameter estimate and confidence interval for CDMV group are reported. Outcomes indicated with an asterisk (*) are log-transformed (log10) for normal distribution; original (upper) and back-transformed (lower) adjusted means and standard errors are reported for these outcomes. Non-adjusted means at week 0 and week 6 can be found in **Supplementary table 10**.

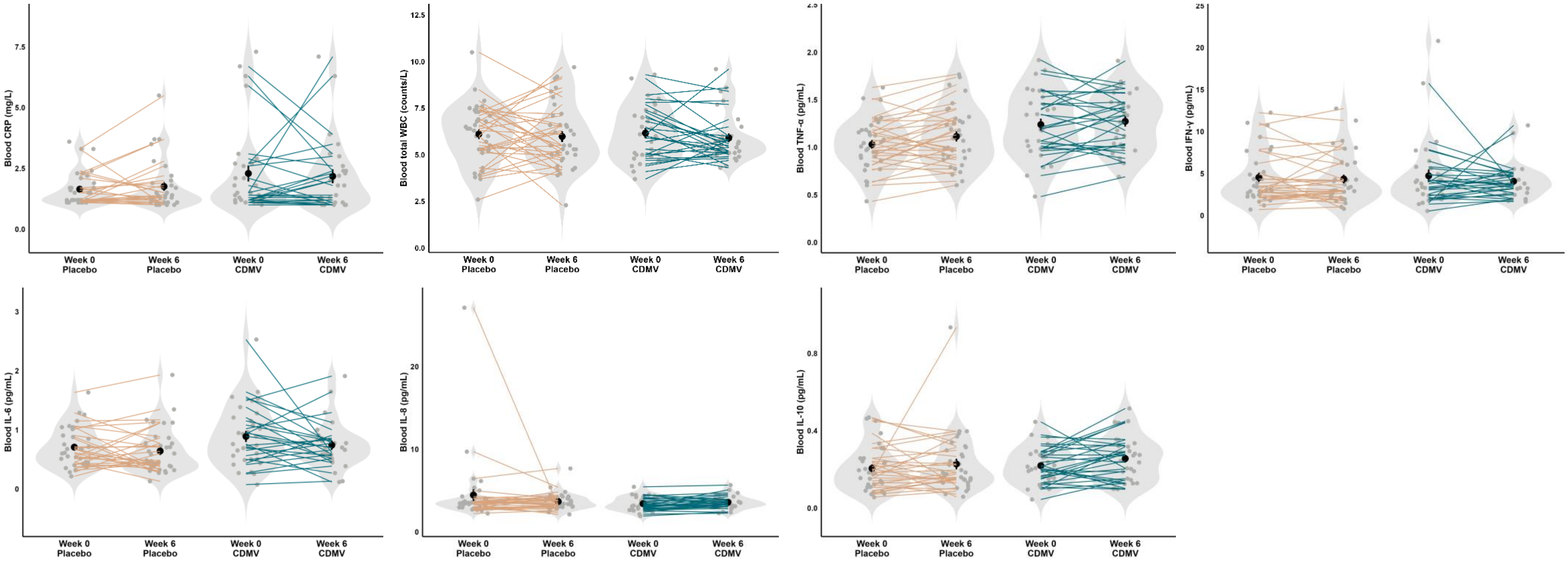


**Supplementary Figure 9. Blood immune markers at baseline and after the 6-weeks intervention for placebo and CDMV group (for *n* per group, see Supplementary Table 7).** *Individual paired samples are connected by a line. The width of the violin shapes indicates the sample density, the circled shape inside indicates the group mean and SE. CDMV = colon-delivered multivitamin; hs-CRP = high-sensitivity C-reactive protein; WBC = white blood cell.*

#### Neurocognitive outcomes

##### Supplementary table 8. Intervention-induced effects in placebo and CDMV group on secondary neuroimaging and neuropsychological outcomes in older adults at risk of cognitive decline

| Outcome | Basic linear model | | | | | | Final linear model with covariates | | | | | |
|---|---|---|---|---|---|---|---|---|---|---|---|---|
| | Placebo | | CDMV | | Estimate group (CI) | P-value | Placebo | | CDMV | | Estimate group (CI) | P-value |
| | $Mean_{adj}$ (SE) | *n* | $Mean_{adj}$ (SE) | *n* | | | $Mean_{adj}$ (SE) | *n* | $Mean_{adj}$ (SE) | *n* | | |
| ***[1]H-MRS neuroinflammation metabolites in the left dlPFC*** | | | | | | | | | | | | |
| *Myo-inositol (mol/kg)* | 10.32 (0.23) | 31 | 10.31 (0.28) | 20 | -0.02 (-0.74; 0.71) | 0.97 | 10.31 (0.23) | 31 | 10.38 (0.29) | 20 | 0.07 (-0.68; 0.83) | 0.83 |
| *Total choline (mol/kg)* | 2.83 (0.04) | 31 | 2.81 (0.05) | 20 | -0.02 (-0.16; 0.12) | 0.78 | 2.84 (0.05) | 31 | 2.80 (0.06) | 20 | -0.04 (-0.19; 0.11) | 0.30 |
| *Total creatine (mol/kg)* | 12.91 (0.19) | 31 | 12.63 (0.23) | 20 | -0.28 (-0.88; 0.33) | 0.36 | 12.90 (0.19) | 31 | 12.57 (0.25) | 20 | -0.34 (-0.98; 0.30) | 0.11 |
| ***ASL Cerebral blood flow*** | | | | | | | | | | | | |
| *CBF whole-brain grey matter* | 41.66 (1.43) | 30 | 42.26 (1.60) | 24 | 0.60 (-3.70; 4.90) | 0.78 | 44.14 (1.76) | 30 | 44.40 (1.89) | 24 | 0.26 (-3.56; 4.09) | 0.89 |
| *CBF whole-brain white matter* | 34.11 (1.19) | 30 | 32.73 (1.33) | 24 | -1.38 (-4.97; 2.21) | 0.44 | 35.28 (1.55) | 30 | 33.49 (1.69) | 24 | -1.79 (-5.23; 1.65) | 0.30 |
| *CBF dlPFC* | 43.84 (1.71) | 30 | 45.23 (1.92) | 24 | 1.39 | 0.59 | 46.92 (2.15) | 30 | 48.16 (2.33) | 24 | 1.24 | 0.59 |

| | | | | | | | | | | | | |
|---|---|---|---|---|---|---|---|---|---|---|---|---|
| | | | | | (-3.78; 6.55) | | | | | | (-3.41; 5.89) | |
| *CBF hippocampus* | 43.94 (1.57) | 30 | 42.44 (1.76) | 24 | -1.50 | 0.53 | 46.14 (2.04) | 30 | 43.21 (2.24) | 24 | -2.93 | 0.20 |
| | | | | | (-6.26; 3.27) | | | | | | (-7.52; 1.65) | |
| ***Neuropsychological test battery*** | | | | | | | | | | | | |
| *DSST score* | 54.91 (1.03) | 35 | 55.72 (1.09) | 31 | 0.81 | 0.59 | 54.78 (1.06) | 35 | 55.32 (1.15) | 31 | 0.55 | 0.73 |
| | | | | | (-2.18; 3.80) | | | | | | (-2.65; 3.74) | |
| *DST total score* | 15.60 (0.35) | 35 | 15.97 (0.37) | 31 | 0.38 | 0.46 | 15.57 (0.36) | 35 | 15.97 (0.40) | 31 | 0.41 | 0.46 |
| | | | | | (-0.64; 1.40) | | | | | | (-0.69; 1.50) | |
| *RAVLT delayed score* | 10.18 (0.39) | 35 | 9.31 (0.41) | 31 | -0.88 | 0.13 | 10.14 (0.39) | 35 | 9.15 (0.44) | 31 | -0.99 | 0.11 |
| | | | | | (-2.03; 0.27) | | | | | | (-2.21; 0.23) | |
| *TMT ratio B/A** | 0.31 (0.02) | 35 | 0.33 (0.02) | 31 | 0.01 | 0.69 | 0.31 (0.02) | 35 | 0.34 (0.02) | 31 | 0.03 | 0.31 |
| | *2.09 (0.09)* | | *2.14 (0.10)* | | (-0.04; 0.06) | | *2.06 (0.09)* | | *2.21 (0.11)* | | (-0.03; 0.09) | |
| *VFT score* | 26.62 (0.64) | 35 | 27.88 (0.68) | 31 | 1.25 | 0.18 | 26.63 (0.65) | 35 | 27.95 (0.72) | 31 | 1.32 | 0.18 |
| | | | | | (-0.61; 3.11) | | | | | | (-0.65; 3.30) | |
| *Composite cognition score (Z)* | 0.14 (0.07) | 35 | 0.21 (0.07) | 31 | 0.07 | 0.46 | 0.14 (0.07) | 35 | 0.18 (0.07) | 31 | 0.05 | 0.65 |
| | | | | | (-0.12; 0.26) | | | | | | (-0.16; 0.25) | |

ASL = arterial spin labelling; CBF = cerebral blood flow; CDMV = colon-delivered multivitamin; dlPFC = dorsolateral prefrontal cortex; DST = digit span test; DSST = digit symbol substitution test; $^{1}$H-MRS = proton magnetic resonance spectroscopy; RAVLT = Rey auditory verbal learning test; TMT = trail making test; VFT = verbal fluency test.

Estimated marginal means with standard error at week 6 are reported for basic linear model (adjusted for baseline value only) and final linear model (adjusted for age, sex, education level, and blood vitamin B6 levels). $^{1}$H-MRS outcomes are also adjusted for quality measure RelResA. ASL outcomes are also adjusted for scanner software version. Parameter estimate and confidence interval for CDMV group are reported. Outcomes indicated with an asterisk (*) are log-transformed (log10) for normal distribution; original (upper) and back-transformed (lower) adjusted means and standard errors are reported for these outcomes. Non-adjusted means at week 0 and week 6 can be found in **Supplementary table 10**.

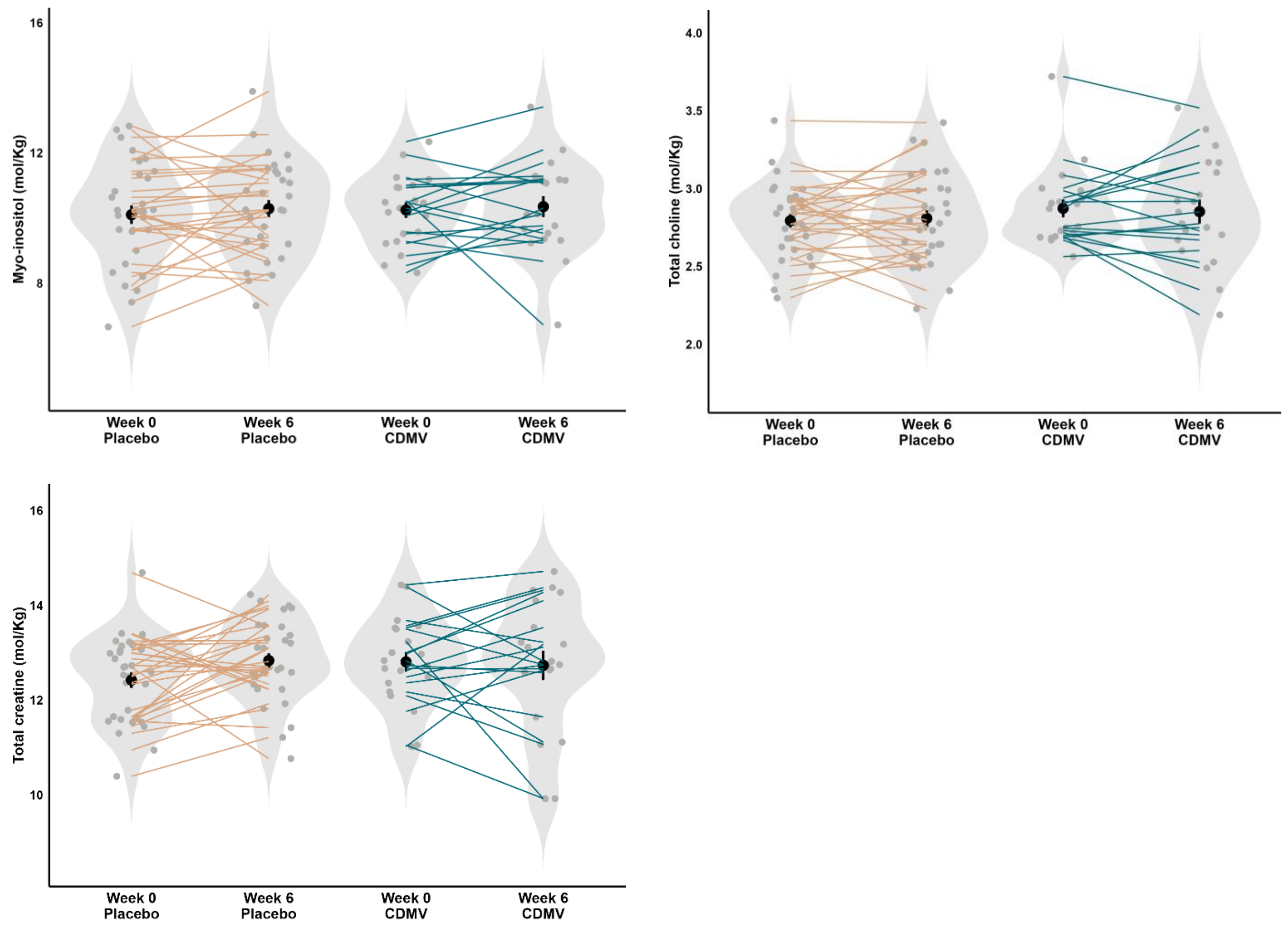


**Supplementary Figure 10. [1]H-MRS Neuroinflammation metabolites within the left dlPFC at baseline and after the 6-weeks intervention for placebo (*n*=31) and CDMV group (*n*=20).** *Myo-inositol, total choline, and total creatine are reported. Individual paired samples are connected by a line. The width of the violin shapes indicates the sample density, the circled shape inside indicates the group mean and SE. CDMV = colon-delivered multivitamin; dlPFC = dorsolateral prefrontal cortex; [1]H-MRS = proton magnetic resonance spectroscopy.*

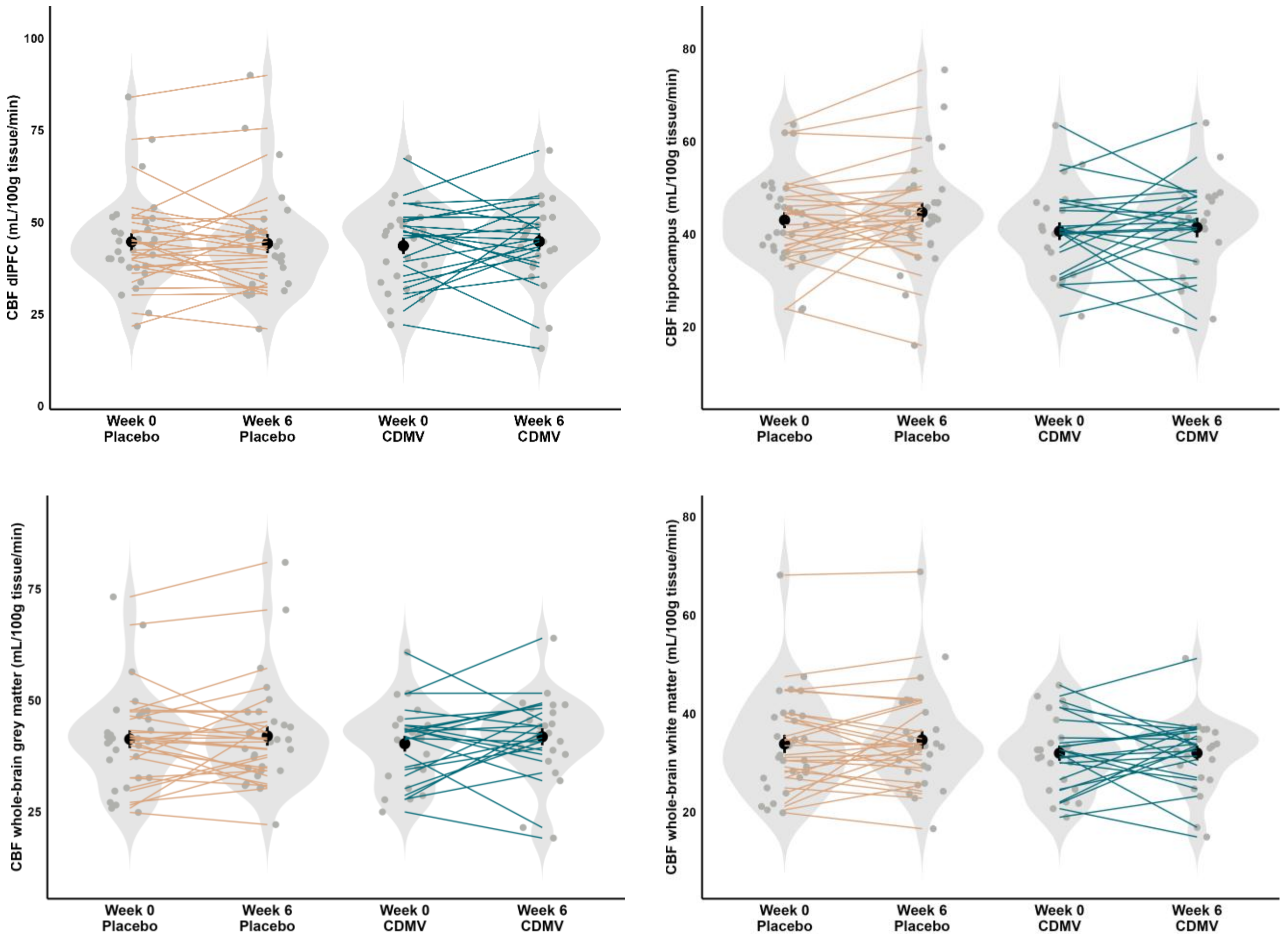


**Supplementary Figure 11. ASL cerebral blood flow levels at baseline and after the 6-weeks intervention for placebo (*n*=30) and CDMV group (*n*=24).** *CBF levels within the dlPFC, hippocampus, whole-brain grey matter and whole-brain white matter are reported. Individual paired samples are connected by a line. The width of the violin shapes indicates the sample density, the circled shape inside indicates the group mean and SE. ASL = arterial spin labeling; CBF = cerebral blood flow; CDMV = colon-delivered multivitamin; dlPFC = dorsolateral prefrontal cortex.*

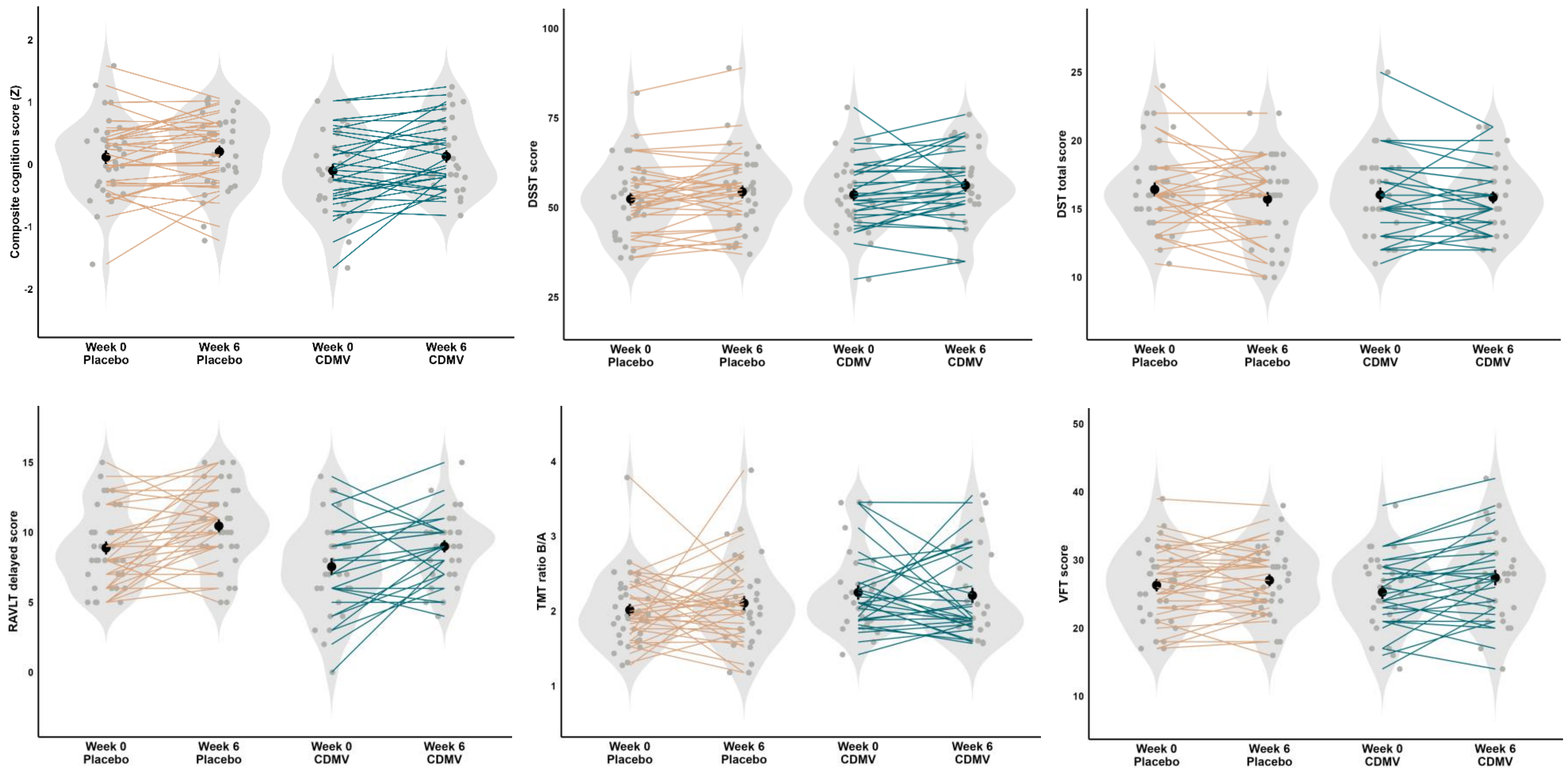


**Supplementary Figure 12. Neuropsychological test battery scores at baseline and after the 6-weeks intervention for placebo (*n*=35) and CDMV group (*n*=31).** *Composite cognition score and individual cognitive test scores for DSST, DST total, RAVLT delayed, TMT ratio B/A and VFT are reported. Individual paired samples are connected by a line. The width of the violin shapes indicates the sample density, the circled shape inside indicates the group mean and SE. CDMV = colon-delivered multivitamin; DST = digit span test; DSST = digit symbol substitution test; RAVLT = Rey auditory verbal learning test; TMT = trail making test; VFT = verbal fluency test.*

**Supplementary table 9. Intervention-induced effects in placebo and CDMV group on perceived stress, anxiety and depression scores in older adults at risk of cognitive decline**

| Outcome | Basic linear model | | | | | | Final linear model with covariates | | | | | |
|---|---|---|---|---|---|---|---|---|---|---|---|---|
| | Placebo | | CDMV | | Estimate group (CI) | P-value | Placebo | | CDMV | | Estimate group (CI) | P-value |
| | Mean$_{adj}$ (SE) | *n* | Mean$_{adj}$ (SE) | *n* | | | Mean$_{adj}$ (SE) | *n* | Mean$_{adj}$ (SE) | *n* | | |
| *PSS stress score* | 17.49 (0.41) | 36 | 17.47 (0.44) | 31 | -0.02 (-1.23; 1.19) | 0.97 | 17.46 (0.42) | 36 | 17.46 (0.46) | 31 | 0.00 (-1.25; 1.25) | 1.00 |
| *HADS anxiety score* | 3.42 (0.29) | 36 | 3.23 (0.31) | 31 | -0.19 (-1.03; 0.65) | 0.65 | 3.37 (0.29) | 36 | 3.18 (0.32) | 31 | -0.19 (-1.05; 0.66) | 0.65 |
| *HADS depression score* | 2.71 (0.29) | 36 | 2.96 (0.31) | 31 | 0.25 (-0.59; 1.19) | 0.55 | 2.71 (0.28) | 36 | 3.08 (0.31) | 31 | 0.38 (-0.47; 1.22) | 0.37 |

CDMV = colon-delivered multivitamin; PSS = Perceived Stress Scale; HADS = Hospital Anxiety and Depression Scale

Estimated marginal means with standard error at week 6 are reported for basic linear model (adjusted for baseline value only) and final linear model (adjusted for age, sex, and education). Parameter estimate and confidence interval for CDMV group are reported. Non-adjusted means at week 0 and week 6 can be found in **Supplementary table 10**.

### Non-adjusted group means

##### Supplementary table 10. Non-adjusted, non-transformed group means at week 0 and week 6 and within group change for all outcomes (gut-, immune- and neurocognitive-related) for placebo and CDMV group in older adults at risk of cognitive decline

| Outcome | Placebo | | | | CDMV | | | |
|---|---|---|---|---|---|---|---|---|
| | *n* | Week 0 Mean ± SD | Week 6 Mean ± SD | Within group change | *n* | Week 0 Mean ± SD | Week 6 Mean ± SD | Within group change |
| **Fecal short-chain fatty acids** | | | | | | | | |
| Total short-chain fatty acids (μmol/g feces) | 36 | 60.0 ± 22.4 | 55.1 ± 26.8 | -4.96 ± 23.3 | 31 | 50.9 ± 22.2 | 54.4 ± 18.4 | 3.50 ± 24.2 |
| Acetic acid (μmol/g feces) | 36 | 35.6 ± 12.9 | 33.2 ± 15.7 | -2.38 ± 13.8 | 31 | 30.4 ± 12.8 | 32 ± 10.4 | 1.62 ± 13.8 |
| Propionic acid (μmol/g feces) | 36 | 11.2 ± 5.32 | 10.4 ± 6.19 | -0.729 ± 4.77 | 31 | 9.45 ± 4.16 | 9.88 ± 3.51 | 0.432 ± 4.44 |
| Butyric acid (μmol/g feces) | 36 | 11.5 ± 5.93 | 9.72 ± 6.42 | -1.73 ± 6.22 | 31 | 9.55 ± 6.48 | 10.8 ± 5.42 | 1.24 ± 6.83 |
| Valeric acid (μmol/g feces) | 36 | 1.88 ± 0.857 | 1.75 ± 0.916 | -0.122 ± 0.869 | 31 | 1.56 ± 0.681 | 1.76 ± 0.741 | 0.207 ± 0.704 |
| **Fecal medium-chain fatty acids** | | | | | | | | |
| Total medium-chain fatty acids (μmol/g feces) | 36 | 0.913 ± 0.803 | 0.885 ± 1.15 | -0.0286 ± 0.818 | 31 | 0.883 ± 0.740 | 0.916 ± 0.713 | 0.0334 ± 0.474 |
| Hexanoic acid (μmol/g feces) | 36 | 0.745 ± 0.698 | 0.695 ± 0.940 | -0.0500 ± 0.665 | 31 | 0.697 ± 0.593 | 0.745 ± 0.602 | 0.0481 ± 0.414 |
| Heptanoic acid (μmol/g feces) | 36 | 0.169 ± 0.115 | 0.191 ± 0.238 | 0.0214 ± 0.185 | 31 | 0.187 ± 0.191 | 0.173 ± 0.132 | -0.0147 ± 0.119 |
| **Fecal branched-chain fatty acids** | | | | | | | | |
| Total branched-chain fatty acids (μmol/g feces) | 36 | 4.56 ± 1.73 | 4.28 ± 1.56 | -0.289 ± 1.73 | 31 | 3.78 ± 1.36 | 4.27 ± 1.57 | 0.491 ± 1.63 |

| | | | | | | | | |
|---|---|---|---|---|---|---|---|---|
| Isovaleric acid (µmol/g feces) | 36 | 2.93 ± 1.07 | 2.76 ± 0.997 | -0.17 ± 1.11 | 31 | 2.40 ± 0.877 | 2.72 ± 1.02 | 0.316 ± 1.06 |
| Isobutyric acid (µmol/g feces) | 36 | 1.56 ± 0.7 | 1.45 ± 0.593 | -0.113 ± 0.642 | 31 | 1.31 ± 0.479 | 1.48 ± 0.554 | 0.174 ± 0.587 |
| 4-methyl valeric acid (µmol/g feces) | 36 | 0.0786 ± 0.038 | 0.0731 ± 0.0421 | -0.00546 ± 0.0516 | 31 | 0.0724 ± 0.0443 | 0.0742 ± 0.0413 | 0.00180 ± 0.0467 |
| **Blood short-chain fatty acids** | | | | | | | | |
| Total short-chain fatty acids (µM) * | 34 | 127.29 ± 32.3 | 136.72 ± 39.6 | 9.43 ± 37.0 | 31 | 138.05 ± 42.9 | 135.38 ± 44.8 | -2.66 ± 42.7 |
| Acetic acid (µM) * | 34 | 124.48 ± 32.1 | 133.66 ± 39.0 | 9.18 ± 36.3 | 31 | 134.09 ± 42.0 | 131.92 ± 44.6 | -2.17 ± 41.8 |
| Propionic acid (µM) * | 34 | 1.89 ± 0.53 | 2.00 ± 0.62 | 0.11 ± 0.59 | 31 | 2.42 ± 1.62 | 2.14 ± 1.05 | -0.28 ± 1.07 |
| Butyric acid (µM) * | 34 | 0.87 ± 0.41 | 0.94 ± 0.50 | 0.07 ± 0.34 | 31 | 1.42 ± 1.02 | 1.22 ± 0.77 | -0.20 ± 0.54 |
| Valeric acid (µM) * | 10 | 0.12 ± 0.14 | 0.24 ± 0.19 | 0.12 ± 0.24 | 16 | 0.14 ± 0.15 | 0.18 ± 0.18 | 0.04 ± 0.23 |
| **Blood medium-chain fatty acids** | | | | | | | | |
| Total medium-chain fatty acids (µM) * | 34 | 3.68 ± 1.64 | 3.41 ± 1.25 | -0.27 ± 1.41 | 31 | 3.72 ± 1.23 | 3.67 ± 1.01 | -0.05 ± 1.45 |
| Hexanoic acid (µM) * | 34 | 0.54 ± 0.28 | 0.52 ± 0.19 | -0.02 ± 0.23 | 31 | 0.53 ± 0.18 | 0.49 ± 0.09 | -0.03 ± 0.20 |
| Heptanoic acid (µM) * | 34 | 0.08 ± 0.04 | 0.10 ± 0.08 | 0.02 ± 0.08 | 31 | 0.10 ± 0.10 | 0.08 ± 0.04 | -0.02 ± 0.09 |
| Octanoic acid (µM) * | 34 | 0.68 ± 0.28 | 0.63 ± 0.23 | -0.05 ± 0.22 | 31 | 0.70 ± 0.33 | 0.65 ± 0.20 | -0.05 ± 0.34 |
| Decanoic acid (µM) * | 34 | 0.76 ± 0.35 | 0.68 ± 0.28 | -0.08 ± 0.27 | 31 | 0.79 ± 0.33 | 0.78 ± 0.25 | -0.01 ± 0.36 |
| Dodecanoic acid (µM) * | 34 | 1.62 ± 1.12 | 1.47 ± 0.79 | -0.15 ± 0.85 | 31 | 1.60 ± 0.61 | 1.66 ± 0.67 | 0.06 ± 0.84 |
| **Blood branched-chain fatty acids** | | | | | | | | |
| Total branched-chain fatty acids (µM) * | 34 | 1.96 ± 0.90 | 2.06 ± 0.73 | 0.10 ± 0.86 | 31 | 2.01 ± 0.77 | 1.87 ± 0.46 | -0.14 ± 0.82 |
| Isovaleric acid (µM) * | 34 | 1.07 ± 0.58 | 1.13 ± 0.46 | 0.06 ± 0.50 | 31 | 1.09 ± 0.50 | 0.99 ± 0.29 | -0.10 ± 0.53 |

| | | | | | | | | |
|---|---|---|---|---|---|---|---|---|
| Isobutyric acid (μM) * | 34 | 0.52 ± 0.20 | 0.54 ± 0.19 | 0.02 ± 0.23 | 31 | 0.57 ± 0.21 | 0.54 ± 0.16 | -0.03 ± 0.19 |
| 2-methyl butyric acid (μM) * | 34 | 0.36 ± 0.21 | 0.39 ± 0.18 | 0.02 ± 0.23 | 31 | 0.35 ± 0.19 | 0.34 ± 0.14 | -0.01 ± 0.22 |
| **Fecal microbiota alpha-diversity** | | | | | | | | |
| Phylogenetic Diversity | 36 | 27.4 ± 4.65 | 28.2 ± 5.22 | 0.765 ± 2.24 | 31 | 26.8 ± 3.92 | 27.5 ± 4.33 | 0.697 ± 2.34 |
| Shannon Diversity | 36 | 4.33 ± 0.381 | 4.45 ± 0.386 | 0.116 ± 0.355 | 31 | 4.44 ± 0.379 | 4.41 ± 0.463 | -0.0217 ± 0.478 |
| Chao1 | 36 | 328 ± 64.4 | 344 ± 74.9 | 16.5 ± 35.7 | 31 | 322 ± 57.7 | 328 ± 60.8 | 6.10 ± 35.1 |
| **Fecal characteristics** | | | | | | | | |
| Fecal pH | 36 | 7.03 ± 0.32 | 7.15 ± 0.36 | 0.125 ± 0.434 | 31 | 7.08 ± 0.37 | 7.03 ± 0.42 | -0.048 ± 0.491 |
| Fecal redox potential (ORP) | 36 | 117.4 ± 49.9 | 89.5 ± 48.7 | -27.9 ± 52.6 | 31 | 110.8 ± 49.0 | 89.3 ± 58.4 | -21.5 ± 64.8 |
| **Intestinal permeability** | | | | | | | | |
| Blood zonulin (ng/mL) | 36 | 37.7 ± 19.0 | 34.6 ± 14.4 | -3.34 ± 10.9 | 31 | 30.2 ± 13.5 | 28.0 ± 13.8 | -2.24 ± 7.91 |
| Blood LBP (μg/mL) | 33 | 11.8 ± 3.74 | 11.8 ± 4.32 | 0.208 ± 4.42 | 31 | 12.6 ± 4.02 | 13.1 ± 4.71 | 0.494 ± 2.82 |
| **Intestinal inflammation** | | | | | | | | |
| Fecal lipocalin | 28 | 0.348 ± 0.150 | 0.377 ± 0.199 | 0.029 ± 0.160 | 24 | 0.413 ± 0.171 | 0.398 ± 0.210 | -0.015 ± 0.275 |
| **Gastro-intestinal symptoms, stool consistency and transit** | | | | | | | | |
| Fecal water content (%) | 32 | 73.0 ± 6.97 | 71.3 ± 6.47 | -1.66 ± 5.84 | 27 | 73.7 ± 7.25 | 72.8 ± 8.64 | -0.98 ± 6.71 |
| Stool consistency (Bristol stool score)[1] | 36 | 4 (1 - 7) | 4 (1 - 6) | 0 (-4 - 2) | 31 | 4 (2 - 7) | 3 (1 - 7) | -1 (-4 - 3) |
| Gastrointestinal transit time (hours) | 35 | 24.9 ± 14.6 | 29.0 ± 19.9 | 4.10 ± 15.5 | 30 | 25.6 ± 13.7 | 25.3 ± 16.5 | -0.31 ± 15.5 |

| | | | | | | | | |
|---|---|---|---|---|---|---|---|---|
| Stomach acid (score from 1 to 7)[1] | 36 | 1 (1- 3) | 1 (1 - 3) | 0 (-2 - 1) | 31 | 1 (1 - 4) | 1 (1 - 5) | 0 (-2 - 3) |
| Bloating (score from 1 to 7)[1] | 36 | 1 (1 - 5) | 1 (1 - 5) | 0 (-3 - 2) | 31 | 1 (1 - 4) | 1 (1 - 3) | 0 (-2 - 2) |
| Flatulence (score from 1 to 7)[1] | 36 | 2 (1 - 6) | 2 (1 - 5) | 0 (-4 - 2) | 31 | 2 (1 - 5) | 2 (1 - 4) | 0 (-2 - 3) |
| Stomach rumbling (score from 1 to 7)[1] | 36 | 1 (1 - 4) | 1 (1 - 3) | 0 (-2 - 2) | 31 | 1 (1 - 4) | 1 (1 - 4) | 0 (-2 - 2) |
| Constipation (score from 1 to 7)[1] | 36 | 1 (1 - 7) | 1 (1 - 5) | 0 (-3 - 4) | 31 | 1 (1 - 4) | 1 (1 - 4) | 0 (-3 - 2) |
| Diarrhoea (score from 1 to 7)[1] | 36 | 1 (1 - 4) | 1 (1 - 5) | 0 (-3 - 4) | 31 | 1 (1 - 5) | 1 (1 - 4) | 0 (-1 - 3) |
| Loose stools (score from 1 to 7)[1] | 36 | 1 (1 - 4) | 1 (1 - 5) | 0 (-1 - 1) | 31 | 1 (1 - 4) | 1 (1 - 4) | 0 (-2 - 1) |
| Hard stools (score from 1 to 7)[1] | 36 | 1 (1 - 4) | 1 (1 - 5) | 0 (-1 - 4) | 31 | 1 (1 - 3) | 1 (1 - 3) | 0 (-2 - 2) |
| **Blood immune markers** | | | | | | | | |
| Blood hs-CRP (mg/L) | 30 | 1.65 ± 0.69 | 1.76 ± 1.03 | 0.167 ± 0.867 | 27 | 2.29 ± 1.81 | 2.18 ± 1.53 | 0.059 ± 1.777 |
| Blood total WBC (counts×$10^9$/L) | 35 | 6.11 ± 1.6 | 5.98 ± 1.72 | -0.12 ± 1.96 | 29 | 6.16 ± 1.48 | 5.91 ± 1.3 | -0.283 ± 1.432 |
| Blood IFN-γ (pg/mL) | 31 | 4.56 ± 3.09 | 4.36 ± 2.94 | -0.044 ± 2.039 | 30 | 4.74 ± 4.18 | 4.1 ± 2.05 | -0.108 ± 3.355 |
| Blood IL-6 (pg/mL) | 31 | 0.71 ± 0.33 | 0.64 ± 0.39 | -0.033 ± 0.349 | 27 | 0.89 ± 0.52 | 0.74 ± 0.41 | -0.18 ± 0.529 |
| Blood IL-8 (pg/mL) | 33 | 4.46 ± 4.18 | 3.68 ± 1.02 | -0.705 ± 4.28 | 30 | 3.42 ± 0.82 | 3.59 ± 0.79 | 0.196 ± 0.688 |
| Blood IL-10 (pg/mL) | 32 | 0.21 ± 0.12 | 0.23 ± 0.16 | 0.02 ± 0.162 | 31 | 0.22 ± 0.09 | 0.26 ± 0.11 | 0.037 ± 0.104 |
| Blood TNF-α (pg/mL) | 33 | 1.03 ± 0.27 | 1.12 ± 0.31 | 0.09 ± 0.205 | 31 | 1.24 ± 0.35 | 1.27 ± 0.3 | 0.034 ± 0.244 |
| **Working memory fMRI responses** | | | | | | | | |
| dlPFC (bilateral) | 29 | 0.39 ± 0.15 | 0.41 ± 0.17 | 0.03 ± 0.16 | 27 | 0.36 ± 0.12 | 0.45 ± 0.17 | 0.09 ± 0.17 |
| dlPFC (right) | 29 | 0.34 ± 0.16 | 0.36 ± 0.18 | 0.02 ± 0.16 | 27 | 0.35 ± 0.13 | 0.42 ± 0.18 | 0.08 ± 0.18 |

| | | | | | | | |
|---|---|---|---|---|---|---|---|
| dlPFC (left) | 29 | 0.44 ± 0.17 | 0.46 ± 0.19 | 0.03 ± 0.18 | 27 | 0.38 ± 0.15 | 0.48 ± 0.20 | 0.10 ± 0.18 |
| Hippocampus (bilateral) | 29 | -0.04 ± 0.07 | -0.06 ± 0.07 | -0.02 ± 0.07 | 27 | -0.05 ± 0.07 | -0.02 ± 0.07 | 0.02 ± 0.08 |
| Hippocampus (right) | 29 | -0.03 ± 0.08 | -0.06 ± 0.07 | -0.03 ± 0.08 | 27 | -0.03 ± 0.07 | 0.00 ± 0.08 | 0.03 ± 0.10 |
| Hippocampus (left) | 29 | -0.05 ± 0.06 | -0.06 ± 0.09 | -0.01 ± 0.08 | 27 | -0.07 ± 0.07 | -0.05 ± 0.08 | 0.02 ± 0.08 |
| **Working memory fMRI functional connectivity with dlPFC** | | | | | | | | |
| Hippocampus (bilateral) | 29 | -0.06 ± 0.06 | -0.09 ± 0.08 | -0.03 ± 0.07 | 27 | -0.07 ± 0.07 | -0.06 ± 0.07 | 0.02 ± 0.09 |
| Hippocampus (right) | 29 | -0.06 ± 0.08 | -0.09 ± 0.07 | -0.04 ± 0.07 | 27 | -0.06 ± 0.07 | -0.04 ± 0.07 | 0.02 ± 0.10 |
| Hippocampus (left) | 29 | -0.07 ± 0.06 | -0.09 ± 0.09 | -0.02 ± 0.08 | 27 | -0.09 ± 0.08 | -0.08 ± 0.08 | 0.01 ± 0.09 |
| **Working memory performance** | | | | | | | | |
| Dprime 0-back | 34 | 4.24 ± 0.50 | 4.33 ± 0.29 | 0.09 ± 0.53 | 28 | 4.36 ± 0.30 | 4.43 ± 0.13 | 0.08 ± 0.31 |
| Dprime 1-back | 34 | 3.43 ± 0.80 | 3.83 ± 0.54 | 0.41 ± 0.92 | 28 | 3.32 ± 0.75 | 3.73 ± 0.81 | 0.41 ± 0.98 |
| Dprime 2-back | 34 | 2.47 ± 0.77 | 2.81 ± 0.69 | 0.34 ± 0.75 | 28 | 2.46 ± 0.66 | 2.79 ± 0.69 | 0.32 ± 0.54 |
| **$^{1}$H-MRS neuroinflammation metabolites in the left dlPFC** | | | | | | | | |
| Myo-inositol (mol/kg) | 31 | 10.11 ± 1.59 | 10.29 ± 1.48 | 0.18 ± 1.45 | 20 | 10.25 ± 1.09 | 10.35 ± 1.44 | 0.10 ± 1.31 |
| Total choline (mol/kg) | 31 | 2.79 ± 0.24 | 2.81 ± 0.29 | 0.02 ± 0.25 | 20 | 2.87 ± 0.26 | 2.85 ± 0.35 | -0.02 ± 0.24 |
| Total creatine (mol/kg) | 31 | 12.42 ± 0.91 | 12.84 ± 0.86 | 0.41 ± 1.03 | 20 | 12.81 ± 0.93 | 12.73 ± 1.40 | -0.07 ± 1.27 |
| **ASL Cerebral blood flow** | | | | | | | | |
| CBF whole-brain grey matter | 24 | 40.28 ± 8.72 | 41.83 ± 9.35 | 1.55 ± 9.15 | 30 | 41.30 ± 11.05 | 42.00 ± 11.88 | 0.70 ± 7.22 |

| | | | | | | | | |
|---|---|---|---|---|---|---|---|---|
| CBF whole-brain white matter | 24 | 31.99 ± 7.58 | 32.01 ± 7.40 | 0.02 ± 7.87 | 30 | 33.87 ± 10.11 | 34.68 ± 10.00 | 0.81 ± 6.31 |
| CBF dlPFC | 24 | 43.58 ± 11.07 | 44.77 ± 11.42 | 1.19 ± 11.03 | 30 | 44.67 ± 12.77 | 44.21 ± 14.20 | -0.46 ± 8.57 |
| CBF hippocampus | 24 | 40.66 ± 9.35 | 41.47 ± 10.37 | 0.80 ± 9.89 | 30 | 43.05 ± 9.54 | 44.72 ± 11.45 | 1.67 ± 7.98 |
| **Neuropsychological test battery** | | | | | | | | |
| DSST score | 35 | 52.40 ± 10.32 | 54.43 ± 10.68 | 2.03 ± 5.77 | 31 | 53.61 ± 9.72 | 56.26 ± 9.95 | 2.65 ± 6.71 |
| DST total score | 35 | 16.43 ± 3.01 | 15.71 ± 3.10 | -0.71 ± 2.63 | 31 | 16.03 ± 2.98 | 15.84 ± 2.42 | -0.19 ± 1.89 |
| RAVLT delayed score | 35 | 8.89 ± 2.76 | 10.46 ± 2.79 | 1.57 ± 2.76 | 31 | 7.55 ± 3.38 | 9.00 ± 2.42 | 1.45 ± 2.95 |
| TMT ratio B/A | 35 | 2.02 ± 0.48 | 2.11 ± 0.57 | 0.09 ± 0.71 | 31 | 2.25 ± 0.54 | 2.21 ± 0.57 | -0.04 ± 0.66 |
| VFT score | 35 | 26.31 ± 5.43 | 27.03 ± 5.22 | 0.71 ± 3.62 | 31 | 25.26 ± 5.39 | 27.42 ± 6.34 | 2.16 ± 4.11 |
| Composite cognition score (Z) | 35 | 0.12 ± 0.62 | 0.21 ± 0.58 | 0.09 ± 0.42 | 31 | -0.10 ± 0.64 | 0.13 ± 0.56 | 0.23 ± 0.45 |
| **Questionnaires** | | | | | | | | |
| PSS stress score | 36 | 18.11 ± 2.71 | 17.47 ± 2.78 | -0.64 ± 3.13 | 31 | 18.19 ± 2.40 | 17.48 ± 2.45 | -0.71 ± 2.67 |
| HADS anxiety score | 36 | 3.86 ± 2.72 | 3.47 ± 2.55 | -0.39 ± 1.99 | 31 | 3.71 ± 2.78 | 3.16 ± 3.02 | -0.55 ± 1.52 |
| HADS depression score | 36 | 2.78 ± 3.02 | 2.69 ± 3.21 | -0.08 ± 1.70 | 31 | 2.81 ± 2.57 | 2.97 ± 2.36 | 0.16 ± 1.88 |

ASL = arterial spin labelling; CBF = cerebral blood flow; CDMV = colon-delivered multivitamin; hs-CRP = high-sensitivity C-reactive protein; dlPFC = dorsolateral prefrontal cortex; DST = digit span test; DSST = digit symbol substitution test; HADS = Hospital Anxiety and Depression Scale; LBP = lipopolysaccharide binding protein; MRS = magnetic resonance spectroscopy; ORP = oxidation-reduction potential; PSS = Perceived Stress Scale; RAVLT = Rey auditory verbal learning test; TMT = trail making test; VFT = verbal fluency test; WBC = white blood cell.

[1] The non-transformed, non-adjusted median (range) are reported.

### Microbiome analysis

#### Beta-diversity

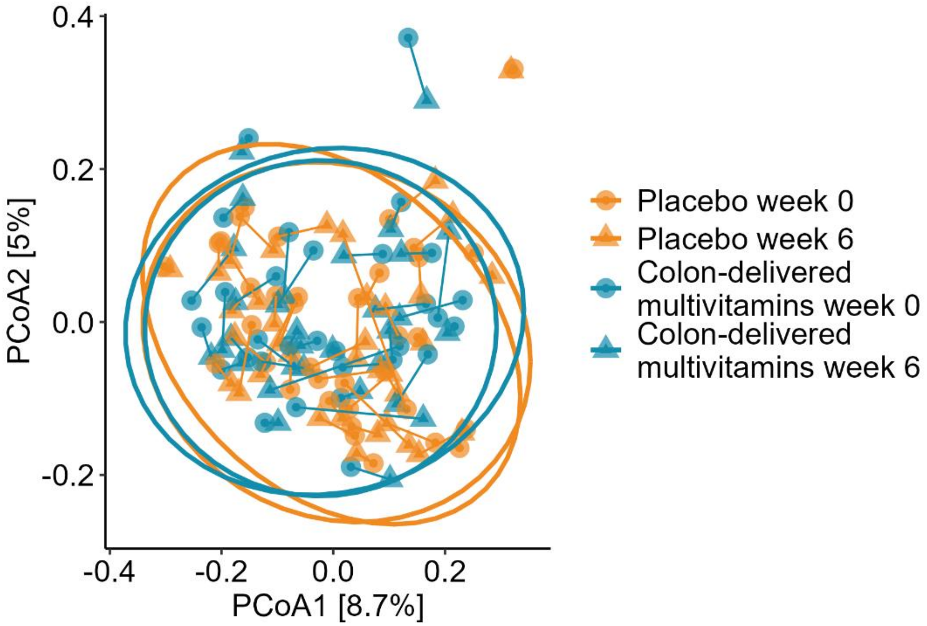


**Supplementary figure 13. Intervention-induced effects on the microbiota beta-diversity for placebo (*n*=36) and CDMV group (*n*=31).** *PCoA plot using Bray-Curtis dissimilarity to visualize the overall microbiota community variation. Color and shape highlight the groups before or after the 6-weeks intervention. The lines connect the within-person samples over time; 95% CIs are plotted. CDMV = colon-delivered multivitamin; PcoA = principal coordinate analysis.*

#### Genus level

**Supplementary Table 11. Fecal bacteria relative abundances at the genus level in placebo and CDMV group at baseline and after 6 weeks of intervention that were found to be significantly different between groups in older adults at risk of cognitive decline.** Only bacteria that passed the 10% prevalence cut-off (present in >13 samples) and had a P-values <0.05 are reported.

| | | Placebo | | CDMV | | Basic linear mixed model | | | Final linear mixed model | | |
|---|---|---|---|---|---|---|---|---|---|---|---|
| **Bacteria** | **MaAsLin3 model [1]** | **Week 0 Rel.ab. % (number)** | **Week 6 Rel. ab. % (number)** | **Week 0 Rel.ab. % (number)** | **Week 6 Rel. ab. % (number)** | **Coefficient (SD)** | **P-value Group x Time** | **FDR P-value Group x Time** | **Coefficient (SD)** | **P-value Group x Time** | **FDR P-value Group x Time** |
| ***Actinomyces*** | abundance | 0.01 ± 0.01 (*n*=14) | 0.01 ± 0.01 (*n*=16) | 0.01 ± 0.03 (*n*=10) | 0.01 ± 0.01 (*n*=13) | -1.13 (0.39) | 0.023 | 0.454 | -1.18 (0.41) | 0.022 | 0.553 |
| ***Anaerofilum*** | abundance | 0 ± 0.01 (*n*=7) | 0 ± 0.01 (*n*=11) | 0 ± 0.01 (*n*=10) | 0 ± 0.01 (*n*=7) | 1.02 (0.32) | 0.003 | 0.184 | 0.91 (0.35) | 0.011 | 0.540 |
| ***Anaeroplasma*** | abundance | 0.04 ± 0.13 (*n*=13) | 0.26 ± 0.8 (*n*=11) | 0.08 ± 0.29 (*n*=8) | 0.05 ± 0.15 (*n*=9) | -2.84 (0.95) | 0.013 | 0.382 | -2.94 (0.94) | 0.010 | 0.525 |
| ***Christensenellaceae_uncultured*** | prevalence | 0.03 ± 0.09 (*n*=22) | 0.04 ± 0.07 (*n*=29) | 0.02 ± 0.02 (*n*=21) | 0.02 ± 0.03 (*n*=20) | -2.43 (1.03) | 0.018 | 0.428 | -2.38 (1.01) | 0.018 | 0.549 |
| ***Catenibacillus*** | abundance | 0 ± 0.01 (*n*=4) | 0 ± 0.01 (*n*=4) | 0.01 ± 0.02 (*n*=7) | 0.01 ± 0.03 (*n*=7) | 1.45 (0.42) | 0.014 | 0.382 | 1.57 (0.40) | 0.006 | 0.421 |
| ***Cloacibacillus*** | abundance | 0.01 ± 0.05 (*n*=4) | 0.01 ± 0.02 (*n*=6) | 0 ± 0.01 (*n*=2) | 0.01 ± 0.04 (*n*=3) | 3.38 (0.83) | 0.021 | 0.447 | 3.90 (0.87) | 0.027 | 0.579 |
| ***Lachnospiraceae_FCS020_group*** | prevalence | 0.26 ± 0.16 (*n*=36) | 0.23 ± 0.17 (*n*=34) | 0.28 ± 0.18 (*n*=30) | 0.27 ± 0.18 (*n*=30) | 11.66 (3.29) | 0.000 | 0.058 | 9.07 (2.32) | 0.000 | **0.027*** |
| ***Rikenella*** | abundance | 0 ± 0 | 0 ± 0.01 | 0 ± 0 | 0.001 ± 0.002 | 1.21 (0.42) | 0.045 | 0.575 | 1.28 (0.41) | 0.036 | 0.591 |

| | | | | | | | | | | |
|---|---|---|---|---|---|---|---|---|---|---|
| | | (*n*=4) | (*n*=4) | (*n*=1) | (*n*=3) | | | | | |
| ***Turicibacter*** | prevalence | 0.11 ± 0.21 (*n*=27) | 0.1 ± 0.3 (*n*=27) | 0.13 ± 0.3 (*n*=28) | 0.08 ± 0.17 (*n*=19) | -2.66 (1.25) | 0.034 | 0.524 | -2.48 (1.18) | 0.035 | 0.582 |
| ***[Eubacterium]_ventriosum_groupD*** | abundance | 0.22 ± 0.27 (*n*=35) | 0.21 ± 0.18 (*n*=35) | 0.28 ± 0.34 (*n*=30) | 0.23 ± 0.18 (*n*=31) | NA. model failed | | | 3.98 (0.02) | 0.548 | **0.031*** |
| ***[Ruminococcus]_gauvreauii_group*** | abundance | 0.69 ± 0.54 (*n*=35) | 0.62 ± 0.49 (*n*=36) | 0.61 ± 0.63 (*n*=27) | 0.59 ± 0.64 (*n*=28) | NA. model failed | | | 1.52 (0.00) | 0.001 | **0.000*** |

CDMV = colon-delivered multivitamin.

Basic linear mixed model (Group x time interaction, adjusted for read depth) and extended linear mixed model (Group x time interaction, adjusted for age, sex, BMI, and habitual dietary fiber intake, and read depth). P-values group x time are reported. [1] MaAsLin3 runs two distinct models, the prevalence model that tests for the presence or absence of bacteria, and the abundance model that tests for the relative abundance of bacteria.

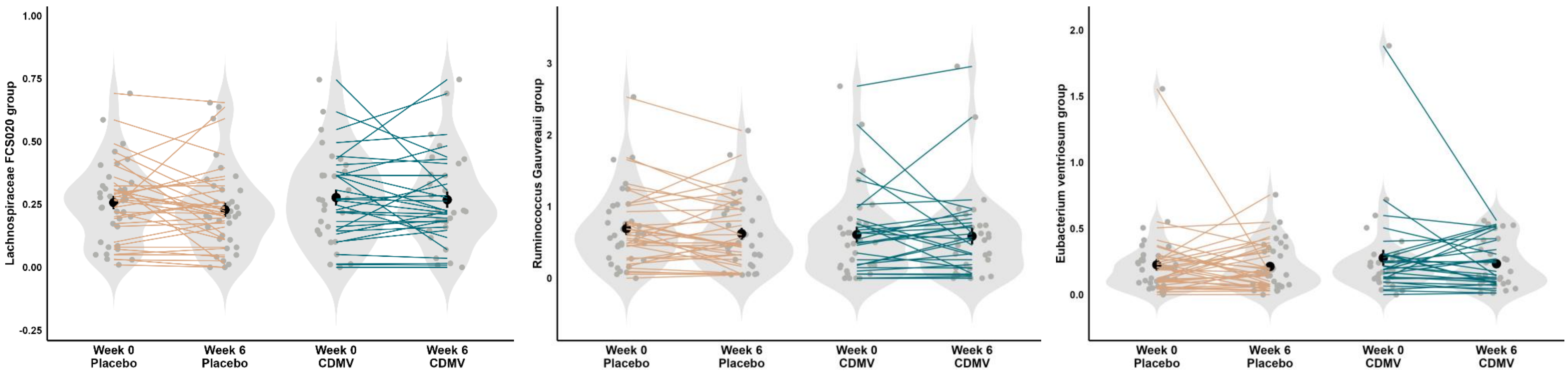


**Supplementary Figure 14. Significant (FDR-corrected p-values) fecal bacteria at baseline and after the 6-weeks intervention for placebo and CDMV group.** *The number of samples in which the bacteria was detected is mentioned in* ***Supplementary table 11.*** *Individual paired samples are connected by a line. The width of the violin shapes indicates the sample density, the circled shape inside indicates the group mean and SE.*

*CDMV = colon-delivered multivitamin.*

**Supplementary table 12. The relative fecal microbiota composition at baseline and after the 6-weeks intervention for placebo and CDMV group.** The mean ± SD of the relative bacteria abundance on genus level (%) is provided. Only the bacteria that had a prevalence of >10% in the total sample subset are included in the table.

| | **Placebo (*n*=36)** | | | | | **CDMV (*n*=31)** | | | | |
|---|---|---|---|---|---|---|---|---|---|---|
| | **Week 0** | **Week 0** | **Week 6** | **Week 6** | **Delta** | **Week 0** | **Week 0** | **Week 6** | **Week 6** | **Delta** |
| **Bacteria (genus)** | **Mean ± SD** | **Count > 0** | **Mean ± SD** | **Count > 0** | **Mean ± SD** | **Mean ± SD** | **Count > 0** | **Mean ± SD** | **Count > 0** | **Mean ± SD** |
| **Bacteroides** | 9.3 ± 8.66 | 36 | 8.35 ± 5.62 | 36 | -0.953 ± 6.944 | 9.09 ± 5.33 | 31 | 8.34 ± 4.67 | 31 | -0.749 ± 6.373 |
| **Unknown** | 4 ± 1.97 | 36 | 4.01 ± 1.74 | 36 | 0.01 ± 1.381 | 3.84 ± 1.57 | 31 | 3.7 ± 1.76 | 31 | -0.145 ± 1.243 |
| **Agathobacter** | 3.78 ± 5.01 | 36 | 2.9 ± 3.49 | 36 | -0.882 ± 4.454 | 2.78 ± 2.3 | 31 | 3.61 ± 3.43 | 31 | 0.826 ± 3.053 |
| **Roseburia** | 3.28 ± 2.32 | 36 | 2.89 ± 2.18 | 36 | -0.391 ± 1.682 | 4.37 ± 2.94 | 31 | 3.2 ± 2.06 | 31 | -1.173 ± 1.809 |
| **Collinsella** | 2.65 ± 2.22 | 36 | 2.23 ± 1.56 | 36 | -0.419 ± 1.567 | 2.92 ± 2.79 | 31 | 2.38 ± 1.93 | 31 | -0.531 ± 1.79 |
| **Bifidobacterium** | 2.64 ± 2.94 | 36 | 3.22 ± 6.8 | 36 | 0.581 ± 5.572 | 3.3 ± 5.42 | 31 | 2.83 ± 4.37 | 31 | -0.471 ± 4.833 |
| **Subdoligranulum** | 2.63 ± 2.28 | 36 | 2.24 ± 1.32 | 36 | -0.394 ± 2.308 | 2.94 ± 1.75 | 31 | 2.9 ± 2.03 | 31 | -0.04 ± 1.947 |

| | | | | | | | | | | |
|---|---|---|---|---|---|---|---|---|---|---|
| **Coprococcus** | 2.34 ± 1.36 | 36 | 2.14 ± 1.45 | 36 | -0.201 ± 0.9 | 1.81 ± 1.16 | 31 | 2.08 ± 1.92 | 31 | 0.27 ± 1.534 |
| **Fusicatenibacter** | 2.15 ± 1.78 | 36 | 2.04 ± 1.55 | 36 | -0.109 ± 1.12 | 2.36 ± 1.55 | 31 | 2.17 ± 1.52 | 31 | -0.188 ± 1.407 |
| **[Eubacterium]_hallii_group** | 2.01 ± 1.13 | 36 | 1.71 ± 0.92 | 36 | -0.295 ± 0.955 | 2.05 ± 1.14 | 31 | 1.67 ± 1 | 31 | -0.379 ± 1.075 |
| **Faecalibacterium** | 10.35 ± 5.84 | 36 | 12.67 ± 7.89 | 36 | 2.325 ± 6.591 | 10.04 ± 7.23 | 31 | 11.04 ± 6.08 | 31 | 0.995 ± 5.769 |
| **Blautia** | 10.18 ± 5.23 | 36 | 8.85 ± 4.59 | 36 | -1.324 ± 4.313 | 11.02 ± 4.85 | 31 | 10.23 ± 5 | 31 | -0.79 ± 4.332 |
| **Anaerostipes** | 1.85 ± 2.23 | 36 | 1.66 ± 1.65 | 36 | -0.19 ± 1.779 | 1.27 ± 0.78 | 31 | 1.42 ± 0.9 | 31 | 0.148 ± 0.844 |
| **Christensenellaceae_R-7_group** | 1.7 ± 2.08 | 36 | 2.04 ± 2.41 | 36 | 0.34 ± 1.817 | 1.69 ± 2.04 | 31 | 1.57 ± 1.87 | 31 | -0.118 ± 1.399 |
| **Oscillospiraceae UCG-002** | 1.62 ± 1.41 | 36 | 2.09 ± 1.8 | 36 | 0.469 ± 1.189 | 2.35 ± 2.54 | 31 | 2.05 ± 1.7 | 31 | -0.299 ± 1.914 |
| **Ruminococcus** | 1.39 ± 1.03 | 36 | 1.47 ± 1.19 | 36 | 0.08 ± 1.045 | 1.78 ± 1.47 | 31 | 1.79 ± 1.65 | 31 | 0.01 ± 1.448 |
| **Romboutsia** | 1.31 ± 3.18 | 36 | 1.22 ± 3.17 | 36 | -0.099 ± 1.491 | 1.04 ± 1.71 | 31 | 0.82 ± 1.01 | 31 | -0.225 ± 1.57 |
| **Dorea** | 1.16 ± 0.78 | 36 | 1.13 ± 0.81 | 36 | -0.038 ± 0.509 | 1.17 ± 0.82 | 31 | 1.06 ± 0.68 | 31 | -0.102 ± 0.464 |
| **[Ruminococcus]_torques_group** | 0.94 ± 0.7 | 36 | 0.88 ± 0.59 | 36 | -0.057 ± 0.707 | 1.19 ± 1.05 | 31 | 1.3 ± 1.62 | 31 | 0.114 ± 0.983 |
| **Clostridium_sensu_stricto_1** | 0.92 ± 1 | 36 | 0.64 ± 0.81 | 34 | -0.281 ± 1.122 | 1.51 ± 1.94 | 31 | 1.17 ± 1.74 | 29 | -0.343 ± 1.392 |

| | | | | | | | | | | |
|---|---|---|---|---|---|---|---|---|---|---|
| **Alistipes** | 0.91 ± 0.77 | 36 | 1.19 ± 1.3 | 36 | 0.276 ± 0.889 | 1.81 ± 3.07 | 31 | 1.71 ± 1.96 | 31 | -0.101 ± 2.747 |
| **Lachnospiraceae_NK4A136_group** | 0.85 ± 0.84 | 36 | 0.93 ± 1.02 | 36 | 0.085 ± 0.618 | 0.85 ± 0.86 | 30 | 0.84 ± 0.77 | 31 | -0.012 ± 0.721 |
| **Lachnospiraceae_ND3007_group** | 0.73 ± 0.69 | 36 | 0.78 ± 0.69 | 36 | 0.048 ± 0.33 | 0.76 ± 0.47 | 31 | 0.88 ± 0.67 | 31 | 0.116 ± 0.471 |
| **Oscillospiraceae NK4A214_group** | 0.72 ± 0.81 | 36 | 0.86 ± 1.09 | 36 | 0.141 ± 0.677 | 0.9 ± 1.17 | 30 | 0.85 ± 1.1 | 31 | -0.052 ± 0.768 |
| **Oscillospiraceae UCG-005** | 0.68 ± 0.77 | 36 | 0.75 ± 0.79 | 36 | 0.071 ± 0.698 | 0.77 ± 0.75 | 31 | 0.72 ± 0.65 | 29 | -0.05 ± 0.626 |
| **[Eubacterium]_coprostanoligenes_group** | 0.55 ± 0.5 | 36 | 0.68 ± 0.59 | 36 | 0.131 ± 0.426 | 0.83 ± 0.59 | 31 | 0.83 ± 0.62 | 31 | -0.002 ± 0.501 |
| **Parabacteroides** | 0.54 ± 0.47 | 36 | 0.59 ± 0.44 | 36 | 0.049 ± 0.402 | 0.71 ± 0.58 | 31 | 0.74 ± 0.52 | 31 | 0.025 ± 0.686 |
| **Lachnospira** | 0.49 ± 0.44 | 36 | 0.57 ± 0.52 | 36 | 0.08 ± 0.482 | 0.7 ± 0.67 | 31 | 0.69 ± 0.64 | 31 | -0.01 ± 0.493 |
| **Butyricicoccus** | 0.44 ± 0.3 | 36 | 0.42 ± 0.32 | 36 | -0.018 ± 0.235 | 0.43 ± 0.22 | 31 | 0.45 ± 0.32 | 30 | 0.016 ± 0.298 |
| **Monoglobus** | 0.35 ± 0.2 | 36 | 0.32 ± 0.25 | 36 | -0.023 ± 0.199 | 0.38 ± 0.59 | 31 | 0.3 ± 0.21 | 30 | -0.08 ± 0.555 |
| **Lachnoclostridium** | 0.26 ± 0.18 | 36 | 0.3 ± 0.38 | 36 | 0.033 ± 0.273 | 0.39 ± 0.57 | 31 | 0.3 ± 0.42 | 31 | -0.092 ± 0.261 |
| **Lachnospiraceae_FCS020_group** | 0.26 ± 0.16 | 36 | 0.23 ± 0.17 | 34 | -0.028 ± 0.136 | 0.28 ± 0.18 | 30 | 0.27 ± 0.18 | 30 | -0.008 ± 0.139 |
| **Incertae_Sedis** | 0.22 ± 0.2 | 36 | 0.24 ± 0.17 | 36 | 0.026 ± 0.217 | 0.21 ± 0.17 | 31 | 0.25 ± 0.21 | 31 | 0.039 ± 0.251 |
| **Ruminococcaceae_uncultured** | 0.21 ± 0.19 | 36 | 0.29 ± 0.29 | 35 | 0.074 ± 0.209 | 0.17 ± 0.12 | 31 | 0.23 ± 0.14 | 31 | 0.057 ± 0.105 |

| | | | | | | | | | |
|---|---|---|---|---|---|---|---|---|---|
| **Streptococcus** | 0.2 ± 0.25 | 36 | 0.21 ± 0.25 | 36 | 0.011 ± 0.318 | 0.97 ± 3.03 | 30 | 0.37 ± 0.72 | 31 | -0.603 ± 3.002 |
| **Oscillibacter** | 0.11 ± 0.11 | 36 | 0.13 ± 0.13 | 36 | 0.022 ± 0.077 | 0.11 ± 0.14 | 30 | 0.12 ± 0.2 | 30 | 0.011 ± 0.085 |
| **[Ruminococcus]_gauvreauii_group** | 0.69 ± 0.54 | 35 | 0.62 ± 0.49 | 36 | -0.07 ± 0.332 | 0.61 ± 0.63 | 27 | 0.59 ± 0.64 | 28 | -0.022 ± 0.477 |
| **Erysipelotrichaceae_UCG-003** | 0.4 ± 0.41 | 35 | 0.47 ± 0.49 | 35 | 0.068 ± 0.407 | 0.65 ± 0.86 | 31 | 0.69 ± 0.68 | 31 | 0.04 ± 0.555 |
| **Marvinbryantia** | 0.24 ± 0.24 | 35 | 0.22 ± 0.14 | 35 | -0.017 ± 0.161 | 0.25 ± 0.18 | 30 | 0.21 ± 0.19 | 28 | -0.038 ± 0.114 |
| **[Eubacterium]_ventriosum_group** | 0.22 ± 0.27 | 35 | 0.21 ± 0.18 | 35 | -0.012 ± 0.292 | 0.28 ± 0.34 | 30 | 0.23 ± 0.18 | 31 | -0.045 ± 0.295 |
| **Family_XIII_AD3011_group** | 0.15 ± 0.11 | 35 | 0.17 ± 0.14 | 34 | 0.016 ± 0.091 | 0.19 ± 0.18 | 31 | 0.15 ± 0.1 | 31 | -0.043 ± 0.136 |
| **Odoribacter** | 0.11 ± 0.09 | 35 | 0.14 ± 0.09 | 36 | 0.028 ± 0.106 | 0.12 ± 0.11 | 29 | 0.13 ± 0.11 | 30 | 0.008 ± 0.112 |
| **Family_XIII_UCG-001** | 0.09 ± 0.05 | 35 | 0.09 ± 0.06 | 35 | 0.001 ± 0.054 | 0.1 ± 0.06 | 30 | 0.1 ± 0.05 | 30 | -0.003 ± 0.046 |
| **Prevotella** | 7.58 ± 18.64 | 34 | 6.29 ± 16.25 | 32 | -1.295 ± 13.941 | 1.11 ± 3.3 | 28 | 3.59 ± 10.38 | 29 | 2.474 ± 8.157 |
| **Escherichia-Shigella** | 2.03 ± 7.96 | 34 | 1.52 ± 5.38 | 35 | -0.512 ± 2.728 | 1.74 ± 6.88 | 30 | 1.15 ± 3.73 | 29 | -0.592 ± 3.495 |
| **Barnesiella** | 0.42 ± 0.51 | 34 | 0.51 ± 0.59 | 33 | 0.091 ± 0.325 | 0.72 ± 1.16 | 30 | 0.55 ± 0.55 | 29 | -0.163 ± 0.998 |
| **Lachnospiraceae_UCG-001** | 0.31 ± 0.26 | 34 | 0.32 ± 0.25 | 33 | 0.006 ± 0.259 | 0.41 ± 0.43 | 27 | 0.44 ± 0.52 | 27 | 0.036 ± 0.382 |

| | | | | | | | | | |
|---|---|---|---|---|---|---|---|---|---|
| **Colidextribacter** | 0.15 ± 0.12 | 34 | 0.17 ± 0.1 | 36 | 0.013 ± 0.063 | 0.2 ± 0.17 | 31 | 0.18 ± 0.19 | 30 | -0.015 ± 0.124 |
| **Lachnospiraceae_uncultured** | 0.1 ± 0.07 | 34 | 0.11 ± 0.08 | 34 | 0.004 ± 0.055 | 0.14 ± 0.13 | 31 | 0.13 ± 0.13 | 31 | -0.013 ± 0.068 |
| **Intestinibacter** | 0.55 ± 0.86 | 33 | 0.36 ± 0.83 | 33 | -0.189 ± 0.736 | 0.55 ± 0.64 | 31 | 0.59 ± 1.28 | 31 | 0.047 ± 1.299 |
| **[Eubacterium]_siraeum_group** | 0.29 ± 0.49 | 33 | 0.51 ± 0.74 | 34 | 0.224 ± 0.645 | 0.3 ± 0.52 | 30 | 0.35 ± 0.46 | 30 | 0.046 ± 0.52 |
| **Sutterella** | 0.26 ± 0.37 | 33 | 0.38 ± 0.58 | 34 | 0.114 ± 0.562 | 0.8 ± 1.14 | 28 | 0.85 ± 1.36 | 26 | 0.05 ± 1.645 |
| **Holdemanella** | 0.58 ± 1.19 | 32 | 0.72 ± 1.76 | 27 | 0.139 ± 1.233 | 1.09 ± 2.45 | 26 | 0.8 ± 2.09 | 23 | -0.297 ± 1.917 |
| **Akkermansia** | 0.31 ± 1.02 | 32 | 0.25 ± 0.48 | 28 | -0.06 ± 0.934 | 0.49 ± 1.37 | 28 | 0.24 ± 0.48 | 25 | -0.245 ± 1.205 |
| **Eggerthellaceae_uncultured** | 0.19 ± 0.2 | 32 | 0.21 ± 0.26 | 32 | 0.02 ± 0.251 | 0.21 ± 0.3 | 28 | 0.23 ± 0.29 | 24 | 0.024 ± 0.12 |
| **Lachnospiraceae GCA-900066575** | 0.06 ± 0.04 | 32 | 0.07 ± 0.06 | 33 | 0.009 ± 0.041 | 0.06 ± 0.05 | 28 | 0.08 ± 0.08 | 30 | 0.022 ± 0.053 |
| **Clostridia_UCG-014** | 0.6 ± 0.81 | 31 | 0.6 ± 0.62 | 33 | -0.003 ± 0.676 | 0.52 ± 0.66 | 30 | 0.37 ± 0.31 | 27 | -0.151 ± 0.549 |
| **Rhodospirillales_uncultured_uncultured** | 0.36 ± 0.92 | 31 | 0.85 ± 1.39 | 34 | 0.49 ± 1.175 | 0.43 ± 0.92 | 25 | 0.35 ± 0.8 | 28 | -0.073 ± 0.53 |
| **[Eubacterium]_eligens_group** | 0.35 ± 0.46 | 31 | 0.38 ± 0.4 | 34 | 0.026 ± 0.506 | 0.48 ± 0.76 | 29 | 0.57 ± 0.7 | 30 | 0.089 ± 0.24 |
| **[Eubacterium]_xylanophilum_group** | 0.23 ± 0.24 | 31 | 0.32 ± 0.27 | 33 | 0.088 ± 0.207 | 0.28 ± 0.32 | 27 | 0.33 ± 0.32 | 28 | 0.044 ± 0.27 |
| **Oscillospiraceae UCG-003** | 0.17 ± 0.19 | 31 | 0.25 ± 0.32 | 31 | 0.076 ± 0.263 | 0.3 ± 0.31 | 28 | 0.37 ± 0.5 | 29 | 0.066 ± 0.511 |
| **Oscillospiraceae_uncultured** | 0.13 ± 0.13 | 31 | 0.15 ± 0.13 | 35 | 0.021 ± 0.069 | 0.2 ± 0.19 | 29 | 0.19 ± 0.16 | 30 | -0.009 ± 0.12 |

| | | | | | | | | | |
|---|---|---|---|---|---|---|---|---|---|
| **Phascolarctobacterium** | 0.58 ± 1.02 | 30 | 0.58 ± 0.83 | 34 | -0.008 ± 0.452 | 0.51 ± 0.72 | 30 | 0.76 ± 1.85 | 27 | 0.253 ± 1.894 |
| **Dialister** | 0.33 ± 0.57 | 30 | 0.36 ± 0.71 | 30 | 0.028 ± 0.508 | 0.54 ± 0.66 | 23 | 0.63 ± 0.84 | 27 | 0.088 ± 0.567 |
| **Lachnospiraceae CAG-56** | 0.29 ± 0.32 | 30 | 0.32 ± 0.3 | 31 | 0.026 ± 0.245 | 0.38 ± 0.43 | 25 | 0.33 ± 0.46 | 26 | -0.044 ± 0.237 |
| **Oscillospirales UCG-010** | 0.25 ± 0.36 | 29 | 0.44 ± 0.64 | 29 | 0.19 ± 0.406 | 0.31 ± 0.59 | 26 | 0.33 ± 0.54 | 26 | 0.021 ± 0.582 |
| **Bilophila** | 0.08 ± 0.12 | 29 | 0.09 ± 0.12 | 26 | 0.012 ± 0.067 | 0.09 ± 0.09 | 29 | 0.09 ± 0.09 | 26 | 0.002 ± 0.083 |
| **Adlercreutzia** | 0.07 ± 0.1 | 29 | 0.07 ± 0.1 | 26 | 0.003 ± 0.057 | 0.06 ± 0.07 | 23 | 0.06 ± 0.06 | 25 | -0.004 ± 0.043 |
| **Butyricimonas** | 0.04 ± 0.04 | 29 | 0.09 ± 0.11 | 28 | 0.046 ± 0.085 | 0.13 ± 0.2 | 24 | 0.13 ± 0.16 | 28 | -0.001 ± 0.156 |
| **Muribaculaceae** | 1.08 ± 3.67 | 28 | 0.59 ± 1.86 | 26 | -0.484 ± 2.956 | 0.43 ± 1.17 | 24 | 0.6 ± 1.36 | 21 | 0.171 ± 0.789 |
| **Lachnospiraceae_UCG-004** | 0.12 ± 0.11 | 28 | 0.18 ± 0.13 | 34 | 0.06 ± 0.127 | 0.18 ± 0.15 | 25 | 0.17 ± 0.11 | 26 | -0.012 ± 0.149 |
| **Lachnospiraceae_UCG-010** | 0.08 ± 0.09 | 28 | 0.08 ± 0.1 | 27 | 0.006 ± 0.096 | 0.11 ± 0.14 | 28 | 0.13 ± 0.17 | 23 | 0.016 ± 0.15 |
| **Paraprevotella** | 0.4 ± 0.89 | 27 | 0.29 ± 0.5 | 28 | -0.109 ± 0.814 | 0.42 ± 0.58 | 25 | 0.9 ± 1.81 | 24 | 0.485 ± 1.667 |
| **Parasutterella** | 0.14 ± 0.29 | 27 | 0.15 ± 0.28 | 28 | 0.017 ± 0.16 | 0.15 ± 0.24 | 27 | 0.13 ± 0.2 | 25 | -0.018 ± 0.177 |
| **Turicibacter** | 0.11 ± 0.21 | 27 | 0.1 ± 0.3 | 27 | -0.008 ± 0.235 | 0.13 ± 0.3 | 28 | 0.08 ± 0.17 | 19 | -0.051 ± 0.322 |

| | | | | | | | | | |
|---|---|---|---|---|---|---|---|---|---|
| **Moryella** | 0.02 ± 0.02 | 27 | 0.02 ± 0.02 | 27 | 0.001 ± 0.017 | 0.02 ± 0.02 | 21 | 0.02 ± 0.02 | 21 | 0 ± 0.018 |
| **Gastranaerophilales** | 0.17 ± 0.31 | 26 | 0.55 ± 1.11 | 26 | 0.381 ± 1.006 | 0.19 ± 0.32 | 22 | 0.26 ± 0.57 | 22 | 0.076 ± 0.618 |
| **Enterorhabdus** | 0.11 ± 0.18 | 26 | 0.12 ± 0.2 | 30 | 0.012 ± 0.088 | 0.18 ± 0.3 | 26 | 0.16 ± 0.28 | 25 | -0.019 ± 0.104 |
| **Lachnospiraceae_UCG-008** | 0.06 ± 0.06 | 26 | 0.07 ± 0.07 | 25 | 0.007 ± 0.064 | 0.07 ± 0.09 | 18 | 0.06 ± 0.07 | 17 | -0.014 ± 0.04 |
| **[Eubacterium]_ruminantium_group** | 0.18 ± 0.34 | 25 | 0.34 ± 1 | 26 | 0.162 ± 0.988 | 0.16 ± 0.28 | 24 | 0.23 ± 0.59 | 24 | 0.076 ± 0.546 |
| **Senegalimassilia** | 0.13 ± 0.16 | 25 | 0.15 ± 0.2 | 23 | 0.021 ± 0.183 | 0.12 ± 0.18 | 19 | 0.07 ± 0.09 | 17 | -0.046 ± 0.11 |
| **Negativibacillus** | 0.07 ± 0.11 | 25 | 0.09 ± 0.13 | 26 | 0.015 ± 0.061 | 0.11 ± 0.19 | 23 | 0.12 ± 0.15 | 26 | 0.016 ± 0.109 |
| **Prevotellaceae_NK3B31_group** | 0.7 ± 3.33 | 23 | 0.48 ± 1.64 | 22 | -0.225 ± 2.085 | 0.23 ± 0.74 | 21 | 0.3 ± 0.92 | 18 | 0.073 ± 0.484 |
| **Bacilli RF39** | 0.18 ± 0.42 | 23 | 0.28 ± 0.78 | 24 | 0.099 ± 0.652 | 0.1 ± 0.28 | 16 | 0.07 ± 0.1 | 18 | -0.034 ± 0.284 |
| **Desulfovibrio** | 0.18 ± 0.31 | 23 | 0.28 ± 0.45 | 24 | 0.1 ± 0.375 | 0.29 ± 0.62 | 20 | 0.29 ± 0.5 | 19 | -0.006 ± 0.231 |
| **Clostridia_vadinBB60_group** | 0.27 ± 0.98 | 22 | 0.38 ± 0.87 | 30 | 0.11 ± 0.396 | 0.25 ± 0.64 | 21 | 0.29 ± 0.57 | 22 | 0.044 ± 0.733 |
| **Ruminococcaceae UBA1819** | 0.05 ± 0.1 | 22 | 0.05 ± 0.15 | 24 | 0.005 ± 0.087 | 0.02 ± 0.03 | 21 | 0.02 ± 0.03 | 18 | -0.002 ± 0.025 |
| **Christensenellaceae_uncultured** | 0.03 ± 0.09 | 22 | 0.04 ± 0.07 | 29 | 0.006 ± 0.046 | 0.02 ± 0.02 | 21 | 0.02 ± 0.03 | 20 | 0.002 ± 0.023 |
| **Ruminococcaceae CAG-352** | 0.5 ± 0.98 | 21 | 0.53 ± 1.15 | 24 | 0.028 ± 1.363 | 0.32 ± 0.73 | 12 | 0.43 ± 0.96 | 15 | 0.112 ± 0.628 |
| **Catenibacterium** | 0.35 ± 1.17 | 21 | 0.46 ± 1.35 | 26 | 0.111 ± 0.676 | 0.13 ± 0.57 | 14 | 0.09 ± 0.3 | 17 | -0.049 ± 0.313 |

| | | | | | | | | | | |
|---|---|---|---|---|---|---|---|---|---|---|
| **Coprobacter** | 0.04 ± 0.06 | 21 | 0.07 ± 0.18 | 24 | 0.033 ± 0.141 | 0.1 ± 0.3 | 22 | 0.06 ± 0.09 | 25 | -0.038 ± 0.223 |
| **Defluviitaleaceae_UCG-011** | 0.02 ± 0.03 | 21 | 0.02 ± 0.02 | 20 | -0.002 ± 0.024 | 0.02 ± 0.02 | 16 | 0.01 ± 0.02 | 19 | -0.002 ± 0.022 |
| **Slackia** | 0.11 ± 0.19 | 20 | 0.06 ± 0.09 | 19 | -0.043 ± 0.15 | 0.09 ± 0.19 | 16 | 0.06 ± 0.1 | 16 | -0.036 ± 0.103 |
| **Barnesiellaceae_uncultured** | 0.05 ± 0.11 | 20 | 0.1 ± 0.25 | 22 | 0.054 ± 0.162 | 0.06 ± 0.13 | 18 | 0.13 ± 0.28 | 19 | 0.063 ± 0.245 |
| **Erysipelatoclostridium** | 0.04 ± 0.08 | 20 | 0.05 ± 0.08 | 22 | 0.003 ± 0.071 | 0.03 ± 0.05 | 22 | 0.03 ± 0.04 | 22 | 0.001 ± 0.027 |
| **Flavonifractor** | 0.04 ± 0.06 | 20 | 0.04 ± 0.05 | 23 | -0.002 ± 0.039 | 0.03 ± 0.06 | 17 | 0.03 ± 0.09 | 18 | 0.005 ± 0.039 |
| **Terrisporobacter** | 0.11 ± 0.22 | 19 | 0.09 ± 0.18 | 15 | -0.026 ± 0.219 | 0.19 ± 0.37 | 15 | 0.19 ± 0.44 | 13 | -0.005 ± 0.347 |
| **Peptococcus** | 0.04 ± 0.09 | 19 | 0.05 ± 0.11 | 17 | 0.013 ± 0.042 | 0.06 ± 0.17 | 13 | 0.03 ± 0.06 | 13 | -0.034 ± 0.129 |
| **Tyzzerella** | 0.07 ± 0.13 | 17 | 0.13 ± 0.27 | 19 | 0.066 ± 0.23 | 0.09 ± 0.29 | 15 | 0.08 ± 0.25 | 15 | -0.019 ± 0.068 |
| **Butyricicoccaceae UCG-009** | 0.01 ± 0.01 | 17 | 0.02 ± 0.02 | 21 | 0.006 ± 0.015 | 0.01 ± 0.03 | 9 | 0.01 ± 0.02 | 16 | 0.004 ± 0.021 |
| **Prevotellaceae_uncultured** | 0.27 ± 0.75 | 16 | 0.16 ± 0.45 | 16 | -0.117 ± 0.467 | 0.21 ± 0.68 | 9 | 0.16 ± 0.4 | 9 | -0.055 ± 0.649 |
| **Megamonas** | 0.25 ± 1.32 | 16 | 0.42 ± 2.23 | 14 | 0.173 ± 0.904 | 0.02 ± 0.07 | 18 | 0.09 ± 0.34 | 10 | 0.066 ± 0.307 |
| **Holdemania** | 0.01 ± 0.01 | 16 | 0.01 ± 0.01 | 19 | -0.001 ± 0.01 | 0.01 ± 0.01 | 17 | 0.01 ± 0.01 | 19 | 0.001 ± 0.009 |

| | | | | | | | | | |
|---|---|---|---|---|---|---|---|---|---|
| **Olsenella** | 0.06 ± 0.18 | 15 | 0.06 ± 0.15 | 13 | -0.003 ± 0.067 | 0.07 ± 0.14 | 16 | 0.05 ± 0.08 | 16 | -0.022 ± 0.11 |
| **Coriobacteriales_uncultured_uncultured** | 0.05 ± 0.18 | 15 | 0.04 ± 0.11 | 13 | -0.013 ± 0.082 | 0.01 ± 0.03 | 6 | 0.01 ± 0.03 | 4 | -0.002 ± 0.019 |
| **Peptococcaceae_uncultured** | 0.01 ± 0.02 | 15 | 0.02 ± 0.02 | 20 | 0.004 ± 0.018 | 0.02 ± 0.04 | 17 | 0.02 ± 0.04 | 16 | 0.002 ± 0.023 |
| **Butyrivibrio** | 0.12 ± 0.36 | 14 | 0.3 ± 0.75 | 19 | 0.177 ± 0.527 | 0.09 ± 0.24 | 15 | 0.11 ± 0.29 | 13 | 0.02 ± 0.076 |
| **Erysipelotrichaceae_uncultured** | 0.02 ± 0.03 | 14 | 0.02 ± 0.04 | 14 | 0.004 ± 0.026 | 0.01 ± 0.02 | 7 | 0.02 ± 0.04 | 11 | 0.007 ± 0.038 |
| **Incertae_Sedis_uncultured** | 0.01 ± 0.03 | 14 | 0.02 ± 0.03 | 15 | 0.005 ± 0.017 | 0.03 ± 0.07 | 14 | 0.03 ± 0.06 | 12 | -0.005 ± 0.048 |
| **Chloroplast** | 0.01 ± 0.02 | 14 | 0.04 ± 0.1 | 11 | 0.027 ± 0.103 | 0.04 ± 0.11 | 14 | 0.06 ± 0.29 | 12 | 0.027 ± 0.202 |
| **Ruminococcaceae DTU089** | 0.01 ± 0.02 | 14 | 0.01 ± 0.01 | 17 | 0.001 ± 0.014 | 0.02 ± 0.02 | 17 | 0.02 ± 0.03 | 17 | 0.002 ± 0.029 |
| **Oxalobacter** | 0.01 ± 0.02 | 14 | 0.01 ± 0.02 | 15 | 0.001 ± 0.018 | 0.02 ± 0.02 | 16 | 0.01 ± 0.02 | 13 | -0.004 ± 0.016 |
| **Actinomyces** | 0.01 ± 0.01 | 14 | 0.01 ± 0.01 | 16 | 0.002 ± 0.01 | 0.01 ± 0.03 | 10 | 0.01 ± 0.01 | 13 | -0.004 ± 0.022 |
| **Anaerotruncus** | 0.01 ± 0.01 | 14 | 0.01 ± 0.02 | 14 | 0.001 ± 0.011 | 0.01 ± 0.02 | 10 | 0.01 ± 0.02 | 12 | 0.002 ± 0.013 |
| **Candidatus_Soleaferrea** | 0.01 ± 0.01 | 14 | 0.01 ± 0.02 | 16 | 0.004 ± 0.013 | 0.01 ± 0.01 | 17 | 0.01 ± 0.01 | 14 | -0.001 ± 0.011 |
| **Veillonella** | 0.04 ± 0.18 | 13 | 0.02 ± 0.1 | 8 | -0.019 ± 0.088 | 0.02 ± 0.04 | 14 | 0.01 ± 0.05 | 6 | -0.008 ± 0.034 |
| **Anaeroplasma** | 0.04 ± 0.13 | 13 | 0.26 ± 0.8 | 11 | 0.217 ± 0.744 | 0.08 ± 0.29 | 8 | 0.05 ± 0.15 | 9 | -0.03 ± 0.251 |

| | | | | | | | | | |
|---|---|---|---|---|---|---|---|---|---|
| **[Clostridium]_methylpentosum_group** | 0.01 ± 0.01 | 13 | 0.01 ± 0.03 | 12 | 0.003 ± 0.019 | 0.01 ± 0.01 | 13 | 0.01 ± 0.01 | 17 | 0.004 ± 0.015 |
| **Izemoplasmatales** | 0.04 ± 0.14 | 12 | 0.11 ± 0.26 | 16 | 0.065 ± 0.142 | 0.04 ± 0.11 | 8 | 0.04 ± 0.08 | 15 | -0.005 ± 0.124 |
| **Howardella** | 0.03 ± 0.06 | 12 | 0.03 ± 0.06 | 11 | -0.001 ± 0.039 | 0.05 ± 0.09 | 9 | 0.03 ± 0.06 | 11 | -0.013 ± 0.044 |
| **[Eubacterium]_brachy_group** | 0.01 ± 0.03 | 12 | 0.01 ± 0.02 | 14 | -0.004 ± 0.022 | 0.02 ± 0.03 | 11 | 0.01 ± 0.02 | 12 | -0.003 ± 0.019 |
| **Alloprevotella** | 0.1 ± 0.27 | 11 | 0.12 ± 0.29 | 11 | 0.015 ± 0.171 | 0.1 ± 0.41 | 3 | 0.07 ± 0.28 | 4 | -0.031 ± 0.138 |
| **Eisenbergiella** | 0.02 ± 0.07 | 11 | 0.03 ± 0.08 | 11 | 0.007 ± 0.101 | 0.05 ± 0.19 | 11 | 0.06 ± 0.24 | 11 | 0.01 ± 0.053 |
| **Desulfovibrionaceae_uncultured** | 0.01 ± 0.02 | 11 | 0.02 ± 0.04 | 18 | 0.01 ± 0.031 | 0.01 ± 0.04 | 4 | 0.02 ± 0.05 | 11 | 0.008 ± 0.015 |
| **Lactococcus** | 0.01 ± 0.02 | 11 | 0.02 ± 0.06 | 11 | 0.009 ± 0.057 | 0.03 ± 0.11 | 9 | 0 ± 0.01 | 6 | -0.028 ± 0.107 |
| **Asteroleplasma** | 0.14 ± 0.79 | 10 | 0.19 ± 0.99 | 14 | 0.046 ± 0.205 | 0.31 ± 1.64 | 12 | 1.83 ± 9.46 | 19 | 1.518 ± 9.461 |
| **Gordonibacter** | 0.01 ± 0.02 | 10 | 0.01 ± 0.02 | 14 | 0.003 ± 0.009 | 0.01 ± 0.01 | 11 | 0.01 ± 0.01 | 9 | -0.002 ± 0.007 |
| **Intestinimonas** | 0.01 ± 0.02 | 10 | 0.01 ± 0.02 | 17 | 0.004 ± 0.019 | 0.02 ± 0.03 | 11 | 0.01 ± 0.02 | 15 | -0.003 ± 0.026 |
| **Rikenellaceae_RC9_gut_group** | 0.1 ± 0.29 | 9 | 0.27 ± 1.23 | 10 | 0.173 ± 1.07 | 0.01 ± 0.03 | 7 | 0.01 ± 0.02 | 5 | -0.004 ± 0.024 |
| **Atopobiaceae_uncultured** | 0.02 ± 0.06 | 9 | 0.02 ± 0.06 | 9 | 0.003 ± 0.035 | 0.03 ± 0.09 | 7 | 0.03 ± 0.11 | 6 | 0 ± 0.049 |

| | | | | | | | | | |
|---|---|---|---|---|---|---|---|---|---|
| **Eggerthella** | 0.01 ± 0.04 | 9 | 0.01 ± 0.02 | 12 | -0.005 ± 0.03 | 0.02 ± 0.04 | 8 | 0.02 ± 0.03 | 14 | 0.001 ± 0.021 |
| **Haemophilus** | 0.01 ± 0.03 | 9 | 0.01 ± 0.02 | 9 | -0.003 ± 0.024 | 0.01 ± 0.03 | 9 | 0.01 ± 0.01 | 7 | -0.004 ± 0.021 |
| **Coriobacteriaceae_UCG-003** | 0.01 ± 0.02 | 9 | 0.01 ± 0.02 | 4 | -0.002 ± 0.009 | 0.01 ± 0.06 | 1 | 0 ± 0.03 | 1 | -0.007 ± 0.037 |
| **Merdibacter** | 0.01 ± 0.02 | 9 | 0.01 ± 0.02 | 8 | -0.001 ± 0.014 | 0.01 ± 0.02 | 9 | 0.02 ± 0.04 | 12 | 0.007 ± 0.034 |
| **Lachnospiraceae_NK4B4_group** | 0.01 ± 0.02 | 9 | 0.01 ± 0.03 | 7 | 0.002 ± 0.023 | 0.12 ± 0.59 | 5 | 0.03 ± 0.08 | 5 | -0.096 ± 0.513 |
| **Victivallis** | 0.01 ± 0.02 | 9 | 0.02 ± 0.04 | 13 | 0.011 ± 0.028 | 0.04 ± 0.15 | 10 | 0.05 ± 0.09 | 15 | 0.003 ± 0.165 |
| **Prevotellaceae_UCG-001** | 0.09 ± 0.37 | 8 | 0.18 ± 0.82 | 6 | 0.087 ± 0.454 | 0.03 ± 0.19 | 3 | 0.05 ± 0.26 | 3 | 0.014 ± 0.071 |
| **Megasphaera** | 0.05 ± 0.23 | 8 | 0.08 ± 0.37 | 8 | 0.032 ± 0.156 | 0.11 ± 0.58 | 5 | 0.1 ± 0.52 | 9 | -0.009 ± 0.054 |
| **[Clostridium]_innocuum_group** | 0.02 ± 0.12 | 8 | 0.02 ± 0.06 | 10 | -0.006 ± 0.085 | 0.02 ± 0.08 | 10 | 0.01 ± 0.05 | 12 | -0.008 ± 0.034 |
| **[Ruminococcus]_gnavus_group** | 0.01 ± 0.03 | 8 | 0.03 ± 0.1 | 6 | 0.018 ± 0.083 | 0.13 ± 0.33 | 11 | 0.15 ± 0.44 | 11 | 0.02 ± 0.21 |
| **Oscillospirales_uncultured_uncultured** | 0.01 ± 0.02 | 8 | 0.01 ± 0.02 | 8 | 0.001 ± 0.012 | 0 ± 0.01 | 3 | 0 ± 0.01 | 6 | 0.001 ± 0.017 |
| **Incertae_Sedis DTU014** | 0 ± 0.01 | 8 | 0.01 ± 0.01 | 9 | 0.002 ± 0.006 | 0 ± 0.01 | 6 | 0 ± 0.01 | 7 | 0 ± 0.01 |
| **Faecalitalea** | 0.05 ± 0.27 | 7 | 0.05 ± 0.28 | 6 | 0.001 ± 0.016 | 0 ± 0.01 | 6 | 0.01 ± 0.04 | 5 | 0.007 ± 0.04 |
| **Lactobacillus** | 0.02 ± 0.07 | 7 | 0.01 ± 0.01 | 15 | -0.008 ± 0.073 | 0.02 ± 0.06 | 8 | 0.01 ± 0.02 | 15 | -0.009 ± 0.056 |

| | | | | | | | | | |
|---|---|---|---|---|---|---|---|---|---|
| **Fournierella** | 0.02 ± 0.06 | 7 | 0.02 ± 0.04 | 8 | -0.004 ± 0.04 | 0.03 ± 0.09 | 6 | 0.03 ± 0.08 | 7 | -0.006 ± 0.062 |
| **Allisonella** | 0.01 ± 0.05 | 7 | 0.01 ± 0.02 | 8 | -0.005 ± 0.039 | 0.02 ± 0.04 | 7 | 0.02 ± 0.04 | 8 | -0.001 ± 0.025 |
| **Pseudomonas** | 0.01 ± 0.04 | 7 | 0 ± 0 | 5 | -0.008 ± 0.038 | 0 ± 0 | 3 | 0.01 ± 0.04 | 7 | 0.008 ± 0.037 |
| **Flavobacteriaceae_uncultured** | 0.01 ± 0.03 | 7 | 0.01 ± 0.03 | 6 | -0.001 ± 0.027 | 0.01 ± 0.06 | 7 | 0.01 ± 0.04 | 7 | -0.001 ± 0.054 |
| **[Eubacterium]_nodatum_group** | 0.01 ± 0.03 | 7 | 0.01 ± 0.02 | 9 | -0.002 ± 0.013 | 0 ± 0.01 | 4 | 0 ± 0.01 | 3 | 0 ± 0.008 |
| **Oscillospira** | 0.01 ± 0.02 | 7 | 0.01 ± 0.01 | 9 | -0.001 ± 0.018 | 0.01 ± 0.01 | 6 | 0.01 ± 0.05 | 7 | 0.009 ± 0.037 |
| **Mogibacterium** | 0.01 ± 0.02 | 7 | 0 ± 0.01 | 5 | -0.004 ± 0.012 | 0.01 ± 0.02 | 8 | 0.01 ± 0.02 | 8 | -0.004 ± 0.009 |
| **Anaerofilum** | 0 ± 0.01 | 7 | 0 ± 0.01 | 11 | 0.001 ± 0.006 | 0 ± 0.01 | 10 | 0 ± 0.01 | 7 | 0 ± 0.009 |
| **Christensenella** | 0 ± 0 | 7 | 0 ± 0 | 3 | -0.001 ± 0.005 | 0 ± 0 | 3 | 0 ± 0 | 4 | 0 ± 0.004 |
| **Phocea** | 0 ± 0 | 7 | 0 ± 0.01 | 9 | 0.001 ± 0.006 | 0 ± 0.01 | 8 | 0 ± 0.01 | 6 | -0.001 ± 0.004 |
| **Klebsiella** | 0.12 ± 0.52 | 6 | 0.04 ± 0.18 | 4 | -0.085 ± 0.366 | 0.01 ± 0.03 | 4 | 0 ± 0 | 0 | -0.008 ± 0.03 |

| | | | | | | | | | |
|---|---|---|---|---|---|---|---|---|---|
| **Lachnospiraceae_UCG-003** | 0.03 ± 0.09 | 6 | 0.04 ± 0.13 | 5 | 0.011 ± 0.14 | 0.1 ± 0.32 | 8 | 0.08 ± 0.22 | 8 | -0.022 ± 0.115 |
| **Libanicoccus** | 0.02 ± 0.07 | 6 | 0.02 ± 0.05 | 6 | 0 ± 0.058 | 0.08 ± 0.3 | 5 | 0.05 ± 0.19 | 4 | -0.036 ± 0.122 |
| **Acidaminococcus** | 0.02 ± 0.06 | 6 | 0.02 ± 0.06 | 8 | 0.003 ± 0.059 | 0.08 ± 0.18 | 7 | 0.12 ± 0.29 | 8 | 0.039 ± 0.145 |
| **Enorma** | 0.01 ± 0.03 | 6 | 0.02 ± 0.08 | 3 | 0.008 ± 0.051 | 0 ± 0.01 | 2 | 0.01 ± 0.03 | 3 | 0.006 ± 0.029 |
| **Hungatella** | 0.01 ± 0.03 | 6 | 0 ± 0.01 | 6 | -0.003 ± 0.032 | 0.01 ± 0.05 | 7 | 0.01 ± 0.05 | 4 | 0 ± 0.007 |
| **Anaerofustis** | 0 ± 0.01 | 6 | 0 ± 0.01 | 4 | -0.001 ± 0.006 | 0 ± 0 | 2 | 0 ± 0.01 | 4 | 0.002 ± 0.006 |
| **Solobacterium** | 0.04 ± 0.14 | 5 | 0.03 ± 0.09 | 6 | -0.009 ± 0.103 | 0.06 ± 0.18 | 7 | 0.04 ± 0.12 | 7 | -0.017 ± 0.111 |
| **Mitsuokella** | 0.03 ± 0.12 | 5 | 0.04 ± 0.14 | 4 | 0.006 ± 0.051 | 0.03 ± 0.1 | 3 | 0.16 ± 0.59 | 5 | 0.133 ± 0.531 |
| **Frisingicoccus** | 0.01 ± 0.04 | 5 | 0.02 ± 0.06 | 5 | 0.007 ± 0.045 | 0.03 ± 0.12 | 5 | 0.04 ± 0.14 | 6 | 0.003 ± 0.05 |
| **Paludicola** | 0.01 ± 0.03 | 5 | 0.01 ± 0.04 | 5 | 0.004 ± 0.019 | 0 ± 0.02 | 4 | 0 ± 0.02 | 1 | -0.001 ± 0.023 |
| **Mailhella** | 0.01 ± 0.02 | 5 | 0.02 ± 0.05 | 5 | 0.009 ± 0.027 | 0 ± 0.01 | 2 | 0.01 ± 0.04 | 2 | 0.007 ± 0.031 |
| **Erysipelatoclostridiaceae UCG-004** | 0 ± 0.02 | 5 | 0.02 ± 0.07 | 9 | 0.02 ± 0.063 | 0.01 ± 0.03 | 6 | 0.03 ± 0.08 | 6 | 0.015 ± 0.079 |
| **Caproiciproducens** | 0 ± 0.01 | 5 | 0.01 ± 0.01 | 9 | 0.004 ± 0.01 | 0 ± 0.01 | 5 | 0.01 ± 0.01 | 7 | 0.001 ± 0.009 |
| **Cloacibacillus** | 0.01 ± 0.05 | 4 | 0.01 ± 0.02 | 6 | -0.002 ± 0.032 | 0 ± 0.01 | 2 | 0.01 ± 0.04 | 3 | 0.008 ± 0.029 |

| | | | | | | | | | | |
|---|---|---|---|---|---|---|---|---|---|---|
| **Coprobacillus** | 0.01 ± 0.03 | 4 | 0.01 ± 0.04 | 4 | -0.002 ± 0.049 | 0.04 ± 0.2 | 3 | 0.02 ± 0.13 | 5 | -0.012 ± 0.07 |
| **Sellimonas** | 0.01 ± 0.02 | 4 | 0 ± 0.01 | 4 | -0.001 ± 0.013 | 0.03 ± 0.08 | 6 | 0.03 ± 0.08 | 6 | -0.001 ± 0.024 |
| **Victivallales vadinBE97** | 0.01 ± 0.02 | 4 | 0.01 ± 0.03 | 9 | 0.005 ± 0.021 | 0.03 ± 0.1 | 4 | 0.01 ± 0.02 | 8 | -0.016 ± 0.085 |
| **[Eubacterium]_fissicatena_group** | 0 ± 0.01 | 4 | 0 ± 0 | 2 | -0.001 ± 0.005 | 0 ± 0.01 | 7 | 0 ± 0.01 | 3 | -0.002 ± 0.006 |
| **Catenibacillus** | 0 ± 0.01 | 4 | 0 ± 0.01 | 4 | -0.001 ± 0.007 | 0.01 ± 0.02 | 7 | 0.01 ± 0.03 | 7 | 0.005 ± 0.019 |
| **Shuttleworthia** | 0 ± 0.01 | 4 | 0 ± 0.01 | 2 | -0.001 ± 0.012 | 0.01 ± 0.02 | 7 | 0.01 ± 0.01 | 6 | -0.002 ± 0.017 |
| **Hydrogenoanaerobacterium** | 0 ± 0.01 | 4 | 0 ± 0.01 | 8 | 0.001 ± 0.007 | 0 ± 0.01 | 4 | 0 ± 0 | 3 | -0.001 ± 0.008 |
| **Bacillus** | 0 ± 0.02 | 3 | 0.01 ± 0.02 | 8 | 0.006 ± 0.018 | 0 ± 0.01 | 2 | 0.02 ± 0.02 | 11 | 0.014 ± 0.02 |
| **Enterococcus** | 0 ± 0.01 | 3 | 0 ± 0.01 | 7 | 0.002 ± 0.007 | 0 ± 0.01 | 3 | 0.01 ± 0.01 | 13 | 0.006 ± 0.015 |
| **Oscillospiraceae UCG-007** | 0 ± 0 | 3 | 0 ± 0 | 5 | 0.001 ± 0.005 | 0 ± 0 | 4 | 0 ± 0 | 3 | 0 ± 0.003 |
| **Succinivibrio** | 0.05 ± 0.31 | 2 | 0.01 ± 0.06 | 10 | -0.039 ± 0.25 | 0 ± 0 | 0 | 0 ± 0.01 | 5 | 0.002 ± 0.006 |
| **Listeria** | 0 ± 0.01 | 2 | 0 ± 0.01 | 7 | 0.003 ± 0.007 | 0 ± 0 | 1 | 0.01 ± 0.01 | 10 | 0.008 ± 0.014 |
| **Staphylococcus** | 0 ± 0.01 | 2 | 0 ± 0.01 | 8 | 0.003 ± 0.007 | 0 ± 0.01 | 1 | 0.01 ± 0.01 | 9 | 0.005 ± 0.011 |
| **Lachnospiraceae UC5-1-2E3** | 0 ± 0.01 | 2 | 0 ± 0.01 | 4 | 0 ± 0.007 | 0.01 ± 0.03 | 5 | 0.01 ± 0.03 | 6 | 0 ± 0.014 |
| **Dielma** | 0 ± 0 | 1 | 0 ± 0.01 | 4 | 0.002 ± 0.006 | 0 ± 0 | 5 | 0 ± 0.01 | 6 | 0.002 ± 0.005 |

CDMV = colon-delivered multivitamin.

#### Whole-brain fMRI analysis

**Supplementary table 13. Uncorrected, exploratory whole-brain fMRI peaks for 2b-0b contrast for both group-level contrasts (Clusters of k>3 are reported).**

##### CDMV > Placebo

| Region | MNI peak coordinates | | | Cluster | Peak statistics | |
|---|---|---|---|---|---|---|
| | x | y | z | k | T | p (unc) |
| **Insula, R** | 34 | -22 | 30 | 22 | 4.54 | 0.000 |
| **Sup Temp Pole, L** | -42 | 24 | -20 | 13 | 4.51 | 0.000 |
| **Sup Front G (Orb), R** | 18 | 46 | -16 | 42 | 4.49 | 0.000 |
| **Mid Temp G, L** | -54 | -34 | 2 | 44 | 4.46 | 0.000 |
| *Mid Temp G, L* | -62 | -34 | 6 | | 3.56 | 0.000 |
| **Mid Temp G, R** | 52 | -28 | -14 | 13 | 4.41 | 0.000 |
| **Inf Temp G, R** | 66 | -18 | -32 | 65 | 4.38 | 0.000 |
| *Inf Temp G, R* | 56 | -24 | -32 | | 3.99 | 0.000 |
| **Cerebellum, L** | -48 | -56 | -48 | 23 | 4.35 | 0.000 |
| **Hippocampus, L** | -16 | -18 | -16 | 19 | 4.2 | 0.000 |
| **Cerebellum, L** | -52 | -58 | -54 | 24 | 4.18 | 0.000 |
| **Sup Occ G, R** | 26 | -76 | 34 | 11 | 4.12 | 0.000 |

| | | | | | | |
|---|---|---|---|---|---|---|
| **Cuneus, L** | 2 | -78 | 28 | 18 | 4.11 | 0.000 |
| **Mid Front G, R** | 30 | 28 | 50 | 28 | 4.07 | 0.000 |
| **Mid Front G, R** | 52 | 56 | 0 | 25 | 4.03 | 0.000 |
| **Sup Occ G, L** | -12 | -94 | 10 | 13 | 4.03 | 0.000 |
| **Thalamus, R** | 10 | -16 | -4 | 7 | 3.99 | 0.000 |
| **Ant Cing G, L** | -6 | 20 | 26 | 21 | 3.99 | 0.000 |
| **Precentral G, R** | 52 | 4 | 46 | 25 | 3.95 | 0.000 |
| **Supp Motor Area, R** | 6 | -10 | 50 | 11 | 3.92 | 0.000 |
| **Fusiform, R** | 26 | 4 | -52 | 7 | 3.86 | 0.000 |
| **Inf Front G (Orb), R** | 50 | 42 | -12 | 22 | 3.86 | 0.000 |
| **Rolandic Operculum, R** | 52 | 6 | 4 | 12 | 3.85 | 0.000 |
| **Mid Front G, R** | 40 | 50 | 22 | 11 | 3.85 | 0.000 |
| **Putamen, L** | -28 | 8 | 0 | 5 | 3.84 | 0.000 |
| **Mid Front G, R** | 24 | 10 | 44 | 13 | 3.83 | 0.000 |
| **Supp Motor Area, L** | 2 | 14 | 44 | 13 | 3.79 | 0.000 |
| **Mid Front G, R** | 40 | 24 | 34 | 8 | 3.79 | 0.000 |
| **Ant Cing G, R** | 10 | 50 | 14 | 7 | 3.78 | 0.000 |
| **Mid Cing G, R** | 10 | -26 | 38 | 10 | 3.77 | 0.000 |
| **Precuneus, L** | -6 | -64 | 40 | 5 | 3.74 | 0.000 |
| **Sup Front G (Orb), L** | -20 | 34 | -26 | 5 | 3.73 | 0.000 |

| | | | | | | |
|---|---|---|---|---|---|---|
| **Insula, L** | -32 | -20 | 18 | 10 | 3.72 | 0.000 |
| **Mid Cing G, R** | 8 | 14 | 32 | 9 | 3.72 | 0.000 |
| **Cerebellum, R** | 4 | -50 | -34 | 9 | 3.71 | 0.000 |
| **Precentral G, R** | 46 | -14 | 50 | 5 | 3.71 | 0.000 |
| **Rectus G, L** | -14 | 24 | -10 | 6 | 3.7 | 0.000 |
| **Sup Temp G, L** | -36 | -32 | 10 | 13 | 3.69 | 0.000 |
| **Sup Occ G, L** | -24 | -82 | 26 | 8 | 3.69 | 0.000 |
| **Calcarine, L** | -20 | -74 | 18 | 4 | 3.69 | 0.000 |
| **Mid Cing G, L** | -10 | -32 | 28 | 6 | 3.65 | 0.000 |
| **Sup Temp G, R** | 58 | -2 | -4 | 8 | 3.61 | 0.000 |
| **Inf Front G (Tri), L** | -54 | 20 | 20 | 7 | 3.61 | 0.000 |
| **Inf Front G (Orb), L** | 30 | 34 | -22 | 7 | 3.59 | 0.000 |
| **Cerebellum, L** | -4 | -52 | -24 | 8 | 3.58 | 0.000 |
| **Ant Cing G, L** | -4 | 32 | 30 | 7 | 3.57 | 0.000 |
| **Precentral G, L** | -52 | 6 | 44 | 4 | 3.55 | 0.000 |
| **Inf Temp G, R** | 56 | -58 | -22 | 8 | 3.55 | 0.000 |
| **Postcentral G, L** | -18 | -34 | 66 | 6 | 3.54 | 0.000 |
| **Inf Front G (Tri), R** | 48 | 28 | 4 | 4 | 3.49 | 0.001 |
| **ParaHippocampal G, R** | 18 | -16 | -20 | 4 | 3.47 | 0.001 |
| **Sup Temp Pole, L** | -44 | 16 | -10 | 5 | 3.46 | 0.001 |

| | | | | | | |
|---|---|---|---|---|---|---|
| **Inf Front G (Orb), R** | 26 | 18 | -22 | 4 | 3.45 | 0.001 |
| **Inf Front G (Tri), L** | -46 | 26 | 0 | 4 | 3.42 | 0.001 |
| **Precentral G, R** | 38 | -26 | 60 | 5 | 3.39 | 0.001 |
| **ParaHippocampal G, R** | 14 | -28 | -12 | 5 | 3.36 | 0.001 |

#### Placebo > CDMV

| **Region** | **MNI peak coordinates** | | | **Cluster** | **Peak statistics** | |
|---|---|---|---|---|---|---|
| | **x** | **y** | **z** | **k** | **T** | **p (unc)** |
| **Sup Front G, L** | -24 | 74 | 8 | 35 | 3.96 | 0.000 |
| *Sup Front G, L* | -14 | 74 | 8 | | 3.92 | 0.000 |
| **Inf Temp G, L** | -44 | -22 | -34 | 6 | 3.92 | 0.000 |
| **Caudate, L** | -26 | 2 | 26 | 4 | 3.81 | 0.000 |
| **Cerebellum, L** | -24 | -62 | -60 | 10 | 3.6 | 0.000 |
| **Precuneus, L** | -14 | -56 | 48 | 5 | 3.58 | 0.000 |

Ant = anterior; CDMV = colon-delivered multivitamin; Cing = cingulate; Front = frontal; G = gyrus; Inf = inferior; L = left; Mid = middle; Occ = occipital; Orb = pars orbitalis; R = right; Sup = superior; Supp = Supplementary; Temp = Temporal; Tri = pars triangularis

Height threshold: T = 3.26, p = 0.001 (1.000); Extent threshold: k = 0 voxels; Expected voxels per cluster, <k> = 7.170; Expected number of clusters, <c> = 46.46; Degrees of freedom = [1.0, 51.0]; FWHM = 8.5 8.2 8.1 mm mm mm; 4.2 4.1 4.0 (voxels); Volume: 2503912 = 312989 voxels = 4234.2 resels; Voxel size: 2.0 2.0 2.0 mm mm mm; (resel = 69.85 voxels)

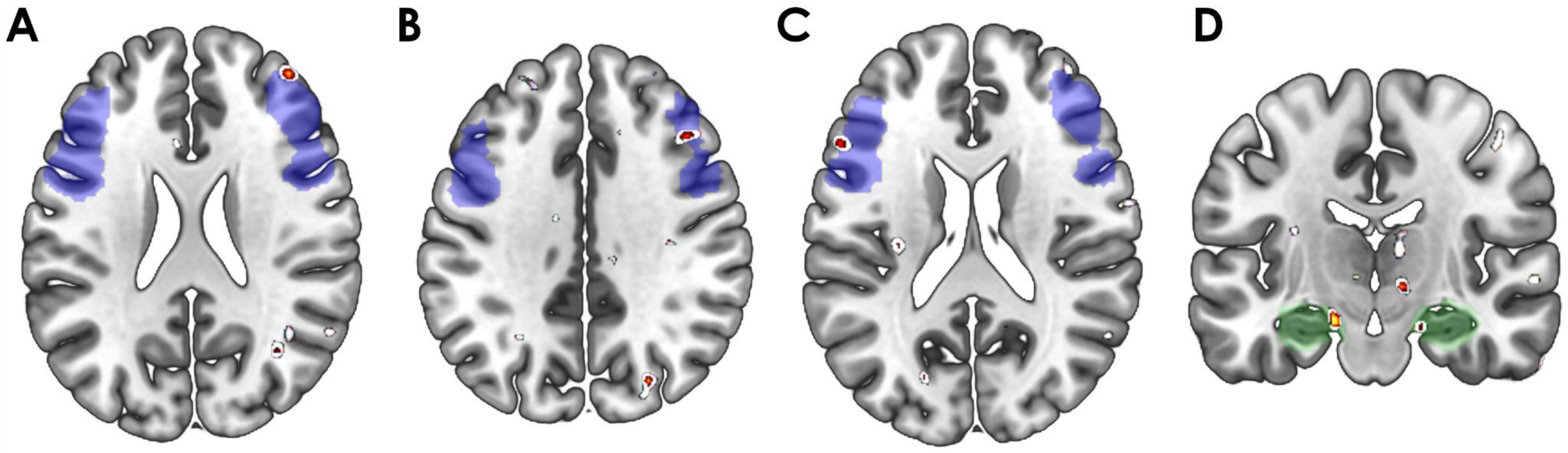


**Supplementary Figure 15. Whole-brain fMRI CDMV > Placebo map for 2b-0b contrast, for illustration purposes.** *The dlPFC (**A, B, C**) and hippocampus (**D**) masks are indicated by the semi-transparent blue (dlPFC) and green (hippocampus) regions. Red-to-yellow clusters show significant peaks for p=0.001 uncorrected; White clusters show significant peaks for p=0.005 uncorrected. CDMV = colon-delivered multivitamin; dlPFC = dorsolateral prefrontal cortex.*

#### Gut-brain correlations over time

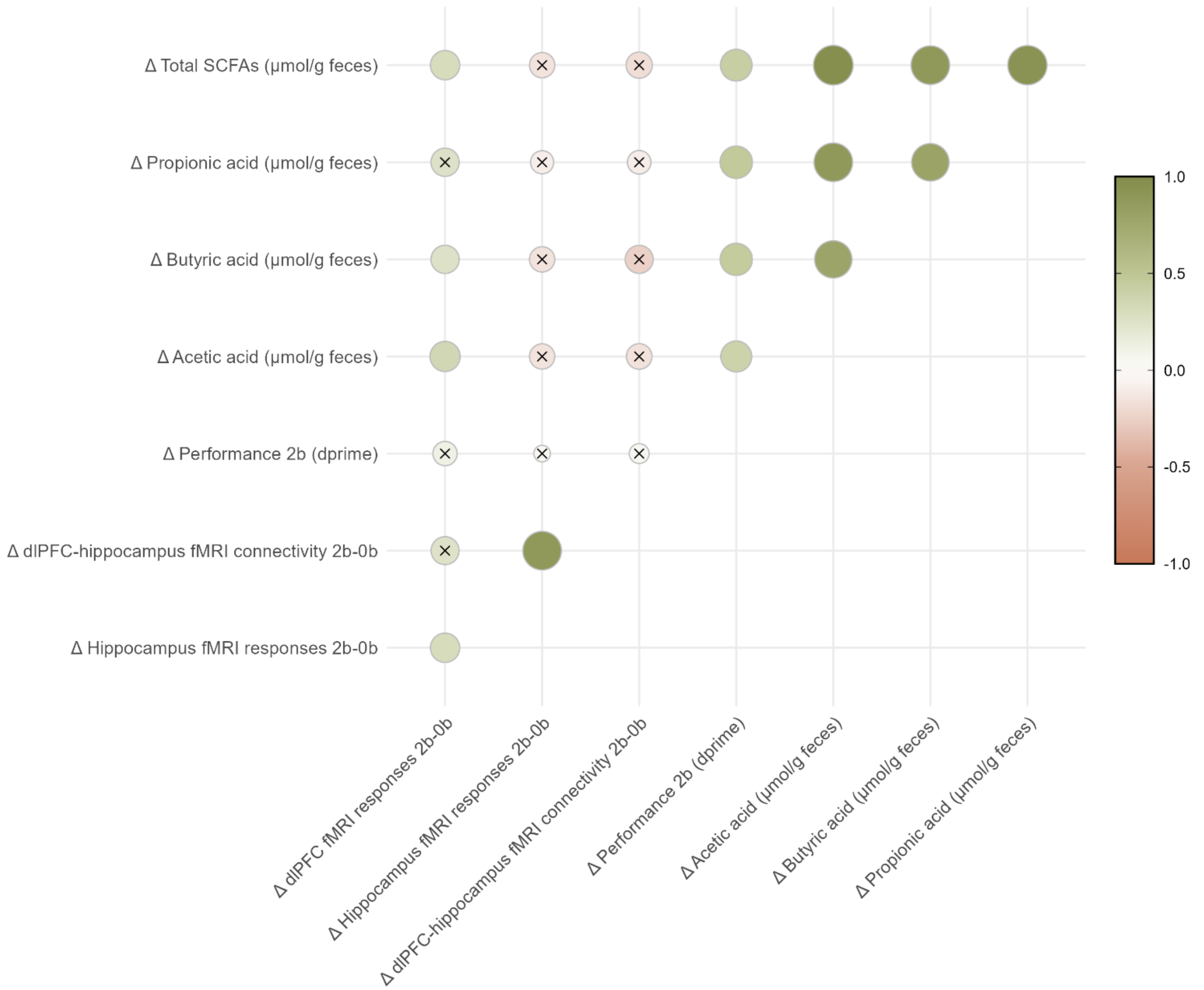

**Supplementary Figure 16. Spearman correlation heatmap of delta primary outcomes: SCFAs, working memory fMRI responses and performance.** Heatmap representing correlations between delta scores (post-value minus pre-value) of SCFAs, working memory-related fMRI responses (n-back task 2-back minus 0-back contrast), working memory-related fMRI connectivity between dlPFC and hippocampus (n-back task 2-back minus 0-back contrast) and working memory performance (n-back task 2-back condition). All delta scores are corrected for baseline (pre) value. Green colors indicate a positive correlation, red colors indicate a negative correlation. Circle size and color intensity indicate strength of the correlation (Spearman's rho). Absence of a cross indicates significance ($p \leq 0.05$) after FDR correction. dlPFC = dorsolateral prefrontal cortex; FDR = false discovery rate; SCFAs = short-chain fatty acids.

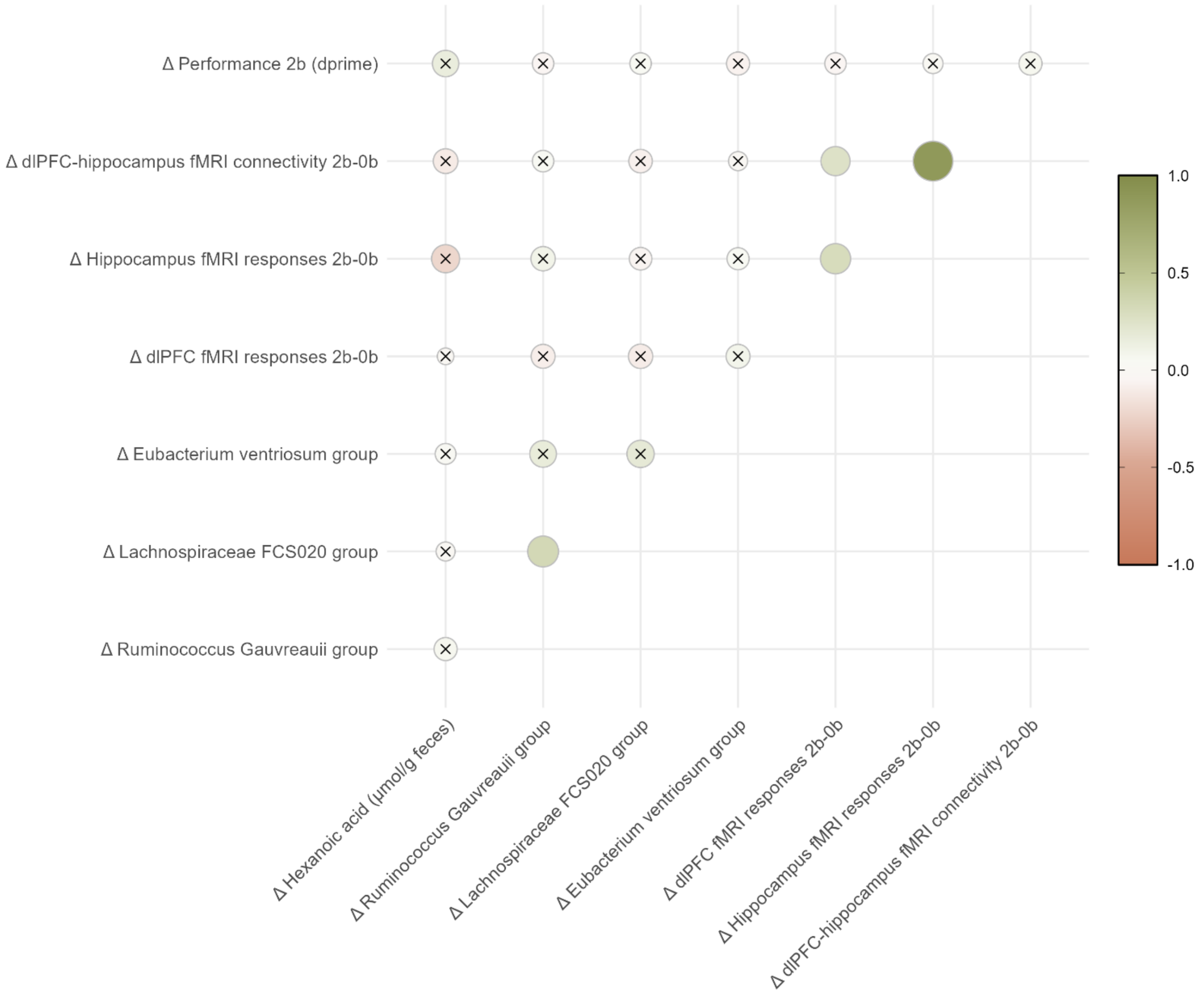

Δ Performance 2b (dprime)
Δ dlPFC-hippocampus fMRI connectivity 2b-0b
Δ Hippocampus fMRI responses 2b-0b
Δ dlPFC fMRI responses 2b-0b
Δ Eubacterium ventriosum group
Δ Lachnospiraceae FCS020 group
Δ Ruminococcus Gauvreauii group
Δ Hexanoic acid (µmol/g feces)
Δ Ruminococcus Gauvreauii group
Δ Lachnospiraceae FCS020 group
Δ Eubacterium ventriosum group
Δ dlPFC fMRI responses 2b-0b
Δ Hippocampus fMRI responses 2b-0b
Δ dlPFC-hippocampus fMRI connectivity 2b-0b
1.0
0.5
0.0
-0.5
-1.0

**Supplementary Figure 17. Spearman correlation heatmap of delta significant secondary gut outcomes with delta working memory fMRI responses and performance.**

Heatmap representing correlations between delta scores (post-value minus pre-value) of fecal hexanoic acid, *Lachnospiraceae FCS020 group*, *Eubacterium ventriosum group* and *Ruminococcus gauvreauii group*, with working memory-related fMRI responses (n-back task 2-back minus 0-back contrast), working memory-related fMRI connectivity between dlPFC and hippocampus (n-back task 2-back minus 0-back contrast) and working memory performance (n-back task 2-back condition). All delta scores are corrected for baseline (pre) value. Green colors indicate a positive correlation, red colors indicate a negative correlation. Circle size and color intensity indicate strength of the correlation (Spearman's rho). Absence of a cross indicates significance ($p \leq 0.05$) after FDR correction. dlPFC = dorsolateral prefrontal cortex; FDR = false discovery rate.

#### References preprocessing with fMRIPrep

Abraham, Alexandre, Fabian Pedregosa, Michael Eickenberg, Philippe Gervais, Andreas Mueller, Jean Kossaifi, Alexandre Gramfort, Bertrand Thirion, and Gael Varoquaux. 2014. "Machine Learning for Neuroimaging with Scikit-Learn." *Frontiers in Neuroinformatics* 8. https://doi.org/10.3389/fninf.2014.00014.

Avants, B. B., C. L. Epstein, M. Grossman, and J. C. Gee. 2008. "Symmetric Diffeomorphic Image Registration with Cross-Correlation: Evaluating Automated Labeling of Elderly and Neurodegenerative Brain." *Medical Image Analysis* 12 (1): 26–41. https://doi.org/10.1016/j.media.2007.06.004.

Behzadi, Yashar, Khaled Restom, Joy Liau, and Thomas T. Liu. 2007. "A Component Based Noise Correction Method (CompCor) for BOLD and Perfusion Based fMRI." *NeuroImage* 37 (1): 90–101. https://doi.org/10.1016/j.neuroimage.2007.04.042.

Ciric, R., William H. Thompson, R. Lorenz, M. Goncalves, E. MacNicol, C. J. Markiewicz, Y. O. Halchenko, et al. 2022. "TemplateFlow: FAIR-Sharing of Multi-Scale, Multi-Species Brain Models." *Nature Methods* 19: 1568–71. https://doi.org/10.1038/s41592-022-01681-2.

Dale, Anders M., Bruce Fischl, and Martin I. Sereno. 1999. "Cortical Surface-Based Analysis: I. Segmentation and Surface Reconstruction." *NeuroImage* 9 (2): 179–94. https://doi.org/10.1006/nimg.1998.0395.

Esteban, Oscar, Ross Blair, Christopher J. Markiewicz, Shoshana L. Berleant, Craig Moodie, Feilong Ma, Ayse Ilkay Isik, et al. 2018. “fMRIPrep 23.2.0.” *Software*. https://doi.org/10.5281/zenodo.852659.

Esteban, Oscar, Christopher Markiewicz, Ross W Blair, Craig Moodie, Ayse Ilkay Isik, Asier Erramuzpe Aliaga, James Kent, et al. 2019. “fMRIPrep: A Robust Preprocessing Pipeline for Functional MRI.” *Nature Methods* 16: 111–16. https://doi.org/10.1038/s41592-018-0235-4.

Evans, AC, AL Janke, DL Collins, and S Baillet. 2012. “Brain Templates and Atlases.” *NeuroImage* 62 (2): 911–22. https://doi.org/10.1016/j.neuroimage.2012.01.024.

Fonov, VS, AC Evans, RC McKinstry, CR Almli, and DL Collins. 2009. “Unbiased Nonlinear Average Age-Appropriate Brain Templates from Birth to Adulthood.” *NeuroImage* 47, Supplement 1: S102. https://doi.org/10.1016/S1053-8119(09)70884-5.

Gorgolewski, K., C. D. Burns, C. Madison, D. Clark, Y. O. Halchenko, M. L. Waskom, and S. Ghosh. 2011. “Nipype: A Flexible, Lightweight and Extensible Neuroimaging Data Processing Framework in Python.” *Frontiers in Neuroinformatics* 5: 13. https://doi.org/10.3389/fninf.2011.00013.

Gorgolewski, Krzysztof J., Oscar Esteban, Christopher J. Markiewicz, Erik Ziegler, David Gage Ellis, Michael Philipp Notter, Dorota Jarecka, et al. 2018. “Nipype.” *Software*. https://doi.org/10.5281/zenodo.596855.

Greve, Douglas N, and Bruce Fischl. 2009. “Accurate and Robust Brain Image Alignment Using Boundary-Based Registration.” *NeuroImage* 48 (1): 63–72. https://doi.org/10.1016/j.neuroimage.2009.06.060.

Jenkinson, Mark, Peter Bannister, Michael Brady, and Stephen Smith. 2002. “Improved Optimization for the Robust and Accurate Linear Registration and Motion Correction of Brain Images.” *NeuroImage* 17 (2): 825–41. https://doi.org/10.1006/nimg.2002.1132.

Klein, Arno, Satrajit S. Ghosh, Forrest S. Bao, Joachim Giard, Yrjö Häme, Eliezer Stavsky, Noah Lee, et al. 2017. "Mindboggling Morphometry of Human Brains." *PLOS Computational Biology* 13 (2): e1005350. https://doi.org/10.1371/journal.pcbi.1005350.

Patriat, Rémi, Richard C. Reynolds, and Rasmus M. Birn. 2017. "An Improved Model of Motion-Related Signal Changes in fMRI." *NeuroImage* 144, Part A (January): 74–82. https://doi.org/10.1016/j.neuroimage.2016.08.051.

Power, Jonathan D., Anish Mitra, Timothy O. Laumann, Abraham Z. Snyder, Bradley L. Schlaggar, and Steven E. Petersen. 2014. "Methods to Detect, Characterize, and Remove Motion Artifact in Resting State fMRI." *NeuroImage* 84 (Supplement C): 320–41. https://doi.org/10.1016/j.neuroimage.2013.08.048.

Reuter, Martin, Herminia Diana Rosas, and Bruce Fischl. 2010. "Highly Accurate Inverse Consistent Registration: A Robust Approach." *NeuroImage* 53 (4): 1181–96. https://doi.org/10.1016/j.neuroimage.2010.07.020.

Satterthwaite, Theodore D., Mark A. Elliott, Raphael T. Gerraty, Kosha Ruparel, James Loughead, Monica E. Calkins, Simon B. Eickhoff, et al. 2013. "An improved framework for confound regression and filtering for control of motion artifact in the preprocessing of resting-state functional connectivity data." *NeuroImage* 64 (1): 240–56. https://doi.org/10.1016/j.neuroimage.2012.08.052.

Tustison, N. J., B. B. Avants, P. A. Cook, Y. Zheng, A. Egan, P. A. Yushkevich, and J. C. Gee. 2010. "N4ITK: Improved N3 Bias Correction." *IEEE Transactions on Medical Imaging* 29 (6): 1310–20. https://doi.org/10.1109/TMI.2010.2046908.

Zhang, Y., M. Brady, and S. Smith. 2001. "Segmentation of Brain MR Images Through a Hidden Markov Random Field Model and the Expectation-Maximization Algorithm." *IEEE Transactions on Medical Imaging* 20 (1): 45–57. https://doi.org/10.1109/42.906424.
